# Species ecology, behavior, and substrate type shape eDNA detection of mammals at savanna waterholes

**DOI:** 10.64898/2026.10.12.758424

**Authors:** Irmgard Seldmayr, Janine Baxter, Rebecca Garbett, Itumeleng Letlojane, Monica Mwale, Byron Weckworth, Tamara Schenekar

## Abstract

Environmental DNA (eDNA) is increasingly used to monitor terrestrial mammal biodiversity, yet factors that influence detection probability in such systems remain poorly understood. Substrates in and around freshwater drinking areas represent particularly promising eDNA sampling points to capture terrestrial biodiversity, especially in arid and semi-arid landscapes. However, eDNA detection probabilities are expected to depend strongly on the frequency and intensity of animal interaction with the sampled substrate at these watering holes.

We combined camera trapping with eDNA metabarcoding to investigate drivers of mammal eDNA detection at savanna waterholes in South Africa (Botsalano Game Reserve) over several weeks during both wet and dry seasons and evaluated the effects of species behavior (interaction intensity with water), species traits, environmental covariates and substrate type (water versus sediment) on eDNA detection probabilities. Camera trapping detected 30 mammal species, whereas eDNA metabarcoding detected 40 species, with 25 species being shared between methods.

Body size increased detection probability using camera traps but not using eDNA metabarcoding. eDNA detection was strongly influenced by trophic level, with herbivores exhibiting significantly higher eDNA detection rates than omnivores and carnivores. eDNA detection probability further increased with behavioral water interaction intensity, social group size, visitation frequency, the number of observed individuals, and declined with time elapsed since the last visit of the respective species. A fine-scale spatial analysis within the water bodies revealed that eDNA deposition is highly localized. Substrate type had a pronounced effect, with sediment samples reflecting more long-term mammal activity, whereas water samples reflected more recent interactions and were more sensitive to environmental conditions. Samples collected at wildlife trails exhibited similar patterns, with trophic level and spatial proximity to the trail both increasing detection rates.

## 1. Introduction

Effective biodiversity monitoring methods provide the foundational data information for many key biodiversity conservation and management decisions (IPBES, 2019). However, all monitoring and assessment approaches of species and ecosystems are subject to biases and limitations, which vary depending on the target taxon and the need for the monitoring (Deiner et al., 2017; Iacaruso et al., 2026). Consequently, it is crucial to choose the most appropriate monitoring method for the study objectives while considering its specific caveats and limitations when interpreting results for implementation.

Camera trapping (CT) and environmental DNA (eDNA)-based monitoring are both non-invasive and now widely adopted methods in biodiversity monitoring (Burton et al., 2015; Deiner et al., 2017). Camera traps are widely used for monitoring mammals, allowing the assessment of species presence, species behavior and activity patterns. With individual identification via unique individual markings, CT data can be used to estimate relative abundance and population density with mark-recapture or mark-resight models (Alonso et al., 2015; O’Brien et al., 2003).

Nevertheless, deployment and maintenance of camera traps can be time- and labor-intensive. It requires regular field visits to service cameras and to replace batteries and storage media to ensure proper long-term functioning. Camera traps must be positioned carefully, with improper placement substantially influencing detection results (Tanwar et al., 2021). They are a novel element in the environment, and often must be concealed and protected against animal and human disturbance. The camera trap emits light, sound and human scent, which may influence animal behavior to a varying extend, depending on the species. Large volumes of image data require substantial storage capacity and processing time. Deep learning, particularly convolutional neural networks (CNNs) has been increasingly used to handle such data (Osner, 2022/2026). However, training efficient models for a specific research question requires substantial amounts of preprocessed data before they can be applied to the biological system under study (Vélez et al., 2023).

The analysis of eDNA has emerged as a promising, cost-effective technique for efficient large-scale biomonitoring (Deiner et al., 2017; Thomsen & Willerslev, 2015). Monitoring using eDNA methods is non-invasive has low or no impact on the target species (Tsuji et al., 2017), providing information on species presence that can be retrieved in a comparatively short timescale (Ishige et al., 2017) with only minimal disruption to the habitat. eDNA metabarcoding, which couples a taxon-group-specific PCR on eDNA samples with high-throughput sequencing, allows the identification of multiple species from a single environmental sample, when supported by well-curated reference databases (Taberlet et al., 2012). A significant advantage of monitoring terrestrial mammals via eDNA versus camera trapping is the detection of small-bodied, cryptic species that are often missed by camera traps (Burton et al., 2015; Leempoel et al., 2020).

However, eDNA analyses are also susceptible to multiple sources of error, such as imperfect detection or (cross-) contamination and incomplete reference databases that can generate both false positive and false negative presence data results (Burian et al., 2021). As eDNA approaches are still developing, many biological, environmental, and methodological factors that influence eDNA detection accuracy are the focus of ongoing research. To reliably assess the risks of false positives and false negatives in eDNA-based surveys, it is crucial to better understand eDNA dynamics in space and time in natural environments. This is essential for efficient study design and for ensuring robust, ecologically meaningful interpretation of eDNA results for each specific region under study.

### Environmental DNA-based monitoring of mammals

Mammals, especially in arid or semi-arid habitats, are well-suited to be monitored by camera trapping and eDNA, as their dependence on water concentrates their activity around waterbodies and their surroundings, which serve as a robust source of terrestrial mammalian eDNA (Rodgers & Mock, 2015; Schenekar et al., 2026; Ushio et al., 2017; Williams et al., 2018).

However, monitoring terrestrial mammals with eDNA it is not yet as established as the monitoring of aquatic organisms (Newton et al., 2025). Several studies have compared camera trapping and eDNA approaches for terrestrial mammal monitoring (Leempoel et al., 2020; Lyet et al., 2021; Mas-Carrió et al., 2022; Sales et al., 2020; Sherub et al., 2026). From these, various factors potentially influencing eDNA detection probabilities have been identified.

Survey duration and sampling effort of eDNA surveys are important survey design factors to consider to effectively compare both survey methods (Lyet et al., 2021; Mas-Carrió et al., 2022). The choice of both the eDNA sample substrate and the precise sampling location is critical, as eDNA is often heterogeneously distributed within environments (Xiao et al., 2025). Abiotic factors such as pH, temperature, conductivity, UV radiation and water volume have been suggested to influence DNA persistence and its rate of degradation (Barnes & Turner, 2016; Joseph et al., 2022) while oxygen saturation has been proposed as an indirect predictor, due to its effect on microbial activity (Barnes et al., 2014) and seasonal differences in eDNA detection of terrestrial animals have been reported (Bai et al., 2025). Additionally, the dynamics of eDNA also vary in differing substrates (Allen et al., 2023; Ionescu et al., 2022; Joseph et al., 2022) and the eDNA degradation rate of sediment is reportedly slower than in the water column in savanna waterholes (Sedlmayr & Schenekar, 2024).

Moreover, several factors derived from species-specific traits of the target taxa that are likely to influence eDNA detection rates, as well. Several studies have suggested that high biomass and body mass, large body volume, and a high number of individuals increase eDNA detection probability (Lyet et al., 2021; Lynggaard et al., 2022; Newton et al., 2024; Yates et al., 2025 ). Temporal dynamics further influence detection, as the number of individuals and visitation history in connection with sampling time and -frequency have been demonstrated to affect eDNA detection (Harper et al., 2019; Lyet et al., 2021; Sales et al., 2020). These patterns likely reflect a combination of increased DNA input due to more mammal activity and eDNA degradation before sampling. eDNA shedding rates also vary by taxon (Andruszkiewicz Allan et al., 2021) with taxa-dependent variation in eDNA detection rates being reported for otters, felids, and reptiles (Adams et al., 2019; Harper et al., 2019; Sales et al., 2020), and a lower eDNA detection signal has been reported for carnivores as opposed to omnivores or herbivores (Lyet et al., 2021). Finally, in contrast to aquatic animals, which live in permanent contact with water, eDNA shedding rates of terrestrial animals are heavily influenced by the type and duration of interaction with the sampled substrate. For example, water interaction experiments with domestic dogs revealed that water interaction intensity increase the amount of eDNA shed into water bodies by individuals (Sauseng & Schenekar, 2025). Harper et al., (2019) investigated how mammal behavior influences eDNA metabarcoding signal strength in captive (zoo) environments. The study recorded the focal species behavior over five days, using camera traps and compared with proportional read counts obtained through eDNA metabarcoding but found no significant link between animal interaction and eDNA signal strength. The authors suggested that the water sources in the study environment may have been saturated with eDNA, potentially masking behavioral effects. Therefore, an assessment of how mammals’ behavior and water interaction intensity affect their eDNA detection probabilities in more natural settings is still needed.

### Study goals

In this study, we link mammalian interactions with waterholes, recorded via camera traps, to the molecular signals detected through eDNA metabarcoding from water and sediment samples collected from these waterholes in a savanna ecosystem.

Specifically, we aim to (1) compare the sensitivity of eDNA metabarcoding with camera trapping by applying both methods in a natural environment at the same location over the same time period, (2) assess how factors such as species-specific traits (body mass, body volume, trophic level, social group size), species behavior (interaction with the sampled substrate), visitation frequencies, spatial distribution of sampling spots, and environmental variables influence the probability of eDNA detection and (3) assess whether these factors affected eDNA detection rate differently in water and sediment samples. To do this, we employed both monitoring methods at three waterholes in a savanna ecosystem in South Africa for a total of 14 weeks (seven weeks in the wet season and seven weeks in the dry season). We then compared species counts of each method, and used a Generalized Linear Mixed Modelling (GLMM) to assess how species traits and activity metrics, as well as environmental variables, influenced eDNA detection probability in general, and separately in water and sediment.

## 2. Methods

### Study site

The study was conducted in Botsalano Game Reserve, a 5800-ha game reserve situated in the North West Province of South Africa. The area is characterized by a semi-arid climate with a wet, rainy season (November-April) and a dry season (May-October). The reserve contains dams along ephemeral streams to retain water during the dry season, forming semi-natural waterholes. The streams typically dry out shortly after the main rainfall period (November–February), resulting in hydrologically independent waterholes. The game reserve is fenced and home to over 20 larger-bodied mammal species (Morris, 2022).

Field work was conducted during two seasons of the same year: during the end of the wet season (08.03.2023 - 23.04.2023) and during the end of the dry season (07.09.2023 - 23.10.2023).

During each sampling season, three waterholes were monitored by both camera trapping and eDNA metabarcoding. In addition, transects on three wildlife trails in proximity (within 60 m) of one waterhole (Monong, see below) were examined similarly for three weeks (1.10.2023-21.10.2023) during the dry season. Although the initial sampling design intended to use the same waterholes across both seasons, one waterhole (Dam 3) dried up at the start of the dry season and had to be replaced with an alternative waterhole (Monong). The waterholes that were sampled and monitored were Kolobe, Mofela, and Dam 3 in the wet season, and Kolobe, Mofela, and Monong during the dry season. Surface areas of all waterholes were measured approximately midway through each sampling period using coarse field measurements. During the wet season, surface areas were approximately 2800 m^2^ for Kolobe, 4500 m^2^ for Mofela and 5900 m^2^ for Dam 3, whereas during the dry season they were 100 m^2^ for Kolobe, 1400 m^2^ for Mofela and 2900 m^2^ for Monong.

### Camera trapping setup

For the monitoring of the waterholes, three to four cameras (10x Bushnell Prime Low Glow 20 MP, 1x Spypoint Solar-Dark No Glow), depending on water body size, were placed around each water body to maximize coverage of all eight eDNA sampling points (Figure 1, Table 1). The cameras were set to take photos at regular intervals (one picture per minute). The infrared motion detection trigger function was not used due to the distances across the waterholes being too great to reliably trigger a recording. Three sampling transects of 5 m of wildlife trails adjacent to Monong waterhole were equipped with one camera each (Bushnell Prime Low Glow 20 MP), set to motion detection and three images per trigger. Cameras were mounted on metal poles approximately 50 cm above ground level and were covered by tree branches and thorns to protect them from animal damage. The area in front of the camera trap was cleared regularly to maintain an unobstructed field of view.

**Figure 1:**
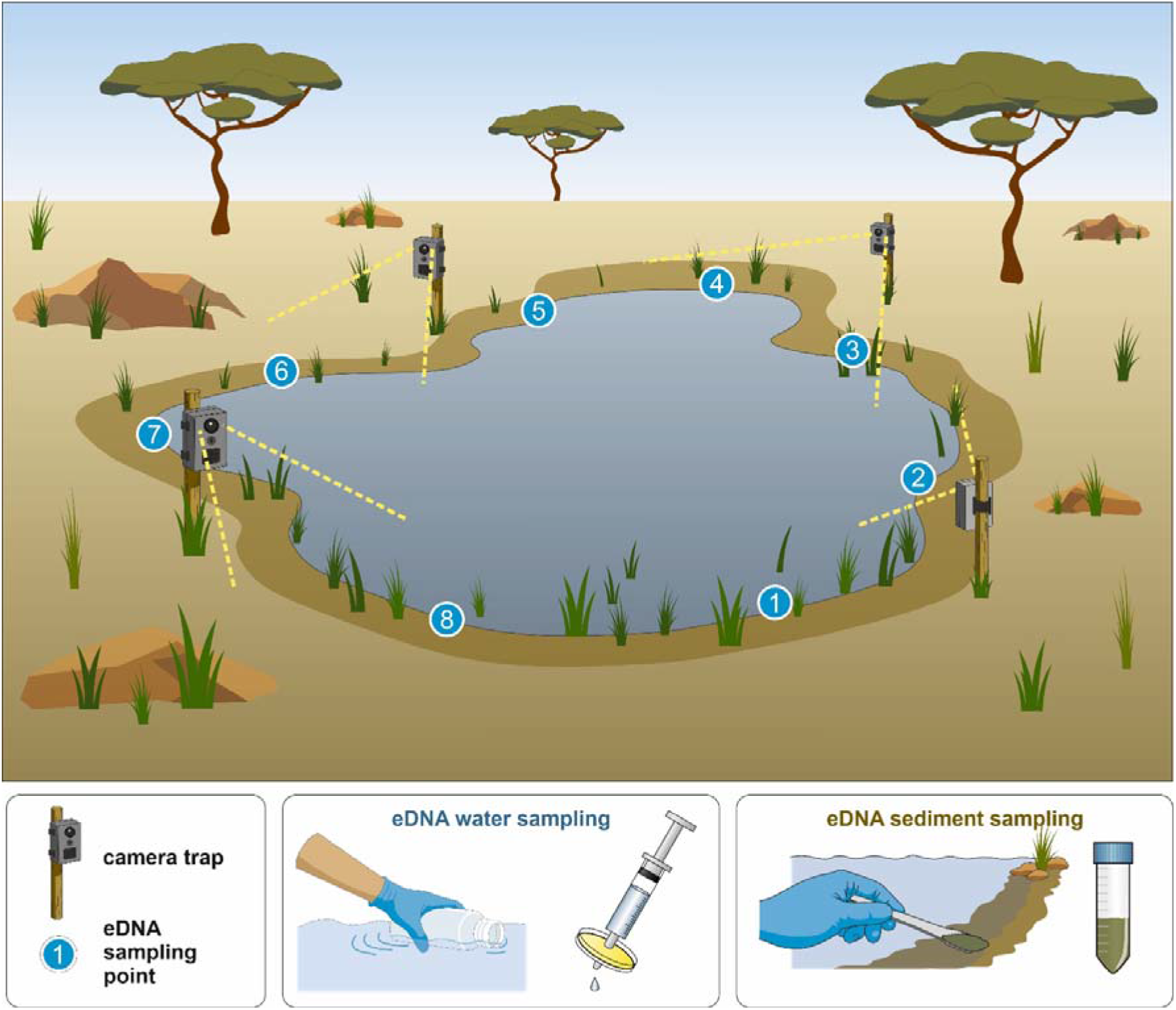
Schematic of the eDNA sampling and camera trapping setup at a waterhole. At each waterhole, eight eDNA sampling points were distributed as evenly as possible along the shore line of the water body. At each sampling point, both eDNA water and sediment samples were collected at each sampling event. Additionally, 3 – 4 camera traps were mounted at a height of approx. 50 cm around the waterhole, aiming to maximize shoreline cover in their views. In addition to water and sediment sampling directly at the waterholes, a limited number of soil samples were also collected along wildlife trails leading to waterholes, which were likewise equipped with camera traps (not shown).

**Table 1:**
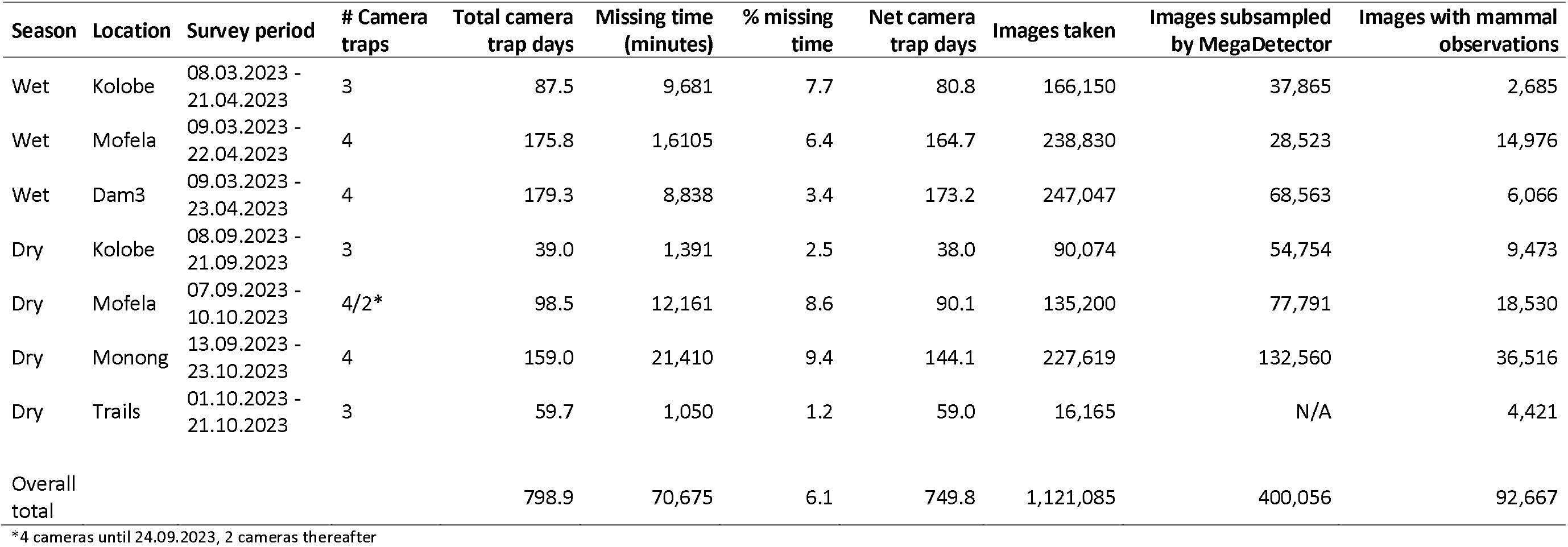
Camera trap survey effort by season and location in Botsalano, South Africa. Table shows survey period, number of deployed camera traps, total camera trapping days (sum of active camera days across all devices), missing time (total minutes when cameras were non-functional or data were lost), percentage of missing time relative to total potential recording time, and net camera trap days (total camera trap days minus missing time). Furthermore, the number of images taken at each survey, as well as the number of images presorted by MegaDetector and the final images containing mammal observations that were exported from Timelapse for further analysis are shown.

| Season | Location | Survey period | # Camera traps | Total camera trap days | Missing time (minutes) | % missing time | Net camera trap days | Images taken | Images subsampled by MegaDetector | Images with mammal observations |
| --- | --- | --- | --- | --- | --- | --- | --- | --- | --- | --- |
| Wet | Kolobe | 08.03.2023 - 21.04.2023 | 3 | 87.5 | 9,681 | 7.7 | 80.8 | 166,150 | 37,865 | 2,685 |
| Wet | Mofela | 09.03.2023 - 22.04.2023 | 4 | 175.8 | 1,6105 | 6.4 | 164.7 | 238,830 | 28,523 | 14,976 |
| Wet | Dam3 | 09.03.2023 - 23.04.2023 | 4 | 179.3 | 8,838 | 3.4 | 173.2 | 247,047 | 68,563 | 6,066 |
| Dry | Kolobe | 08.09.2023 - 21.09.2023 | 3 | 39.0 | 1,391 | 2.5 | 38.0 | 90,074 | 54,754 | 9,473 |
| Dry | Mofela | 07.09.2023 - 10.10.2023 | 4/2* | 98.5 | 12,161 | 8.6 | 90.1 | 135,200 | 77,791 | 18,530 |
| Dry | Monong | 13.09.2023 - 23.10.2023 | 4 | 159.0 | 21,410 | 9.4 | 144.1 | 227,619 | 132,560 | 36,516 |
| Dry | Trails | 01.10.2023 - 21.10.2023 | 3 | 59.7 | 1,050 | 1.2 | 59.0 | 16,165 | N/A | 4,421 |
| Overall total |  |  |  | 798.9 | 70,675 | 6.1 | 749.8 | 1,121,085 | 400,056 | 92,667 |

\*4 cameras until 24.09.2023, 2 cameras thereafter

### Analysis of raw camera trap data

All 1,104,920 photos taken at the waterholes were preclassified with the automated object-detection model MegaDetector (Beery et al., 2018/2023) to categorize photos as empty images and images containing an animal. The “custom recognition sensitivity”, which sets the threshold of an image being categorized as containing an animal was set to 0.04. After the presorting with MegaDetector, a total of 400,056 photos (36%) were retained, which were then analyzed manually using the software Timelapse (https://timelapse.ucalgary.ca, Greenberg et al., 2019). For each image containing mammal detections the following details were recorded (Figure 2): (1) the species, (2) the number of individuals, (3) the closest eDNA sampling point, and (4) their behavior/ interaction with the water body in the following categories: (a) standing nearby (< 10 m distance from the waterbody) (b) drinking (c) minimum of the animal’s feet submerged in the water and (d) minimum of the animal’s at least feet submerged in the water while drinking. If multiple mammal species were present in the same image, the image received separate entries for each species. Only mammals within a 10 m radius of the waterhole were scored.

**Figure 2:**
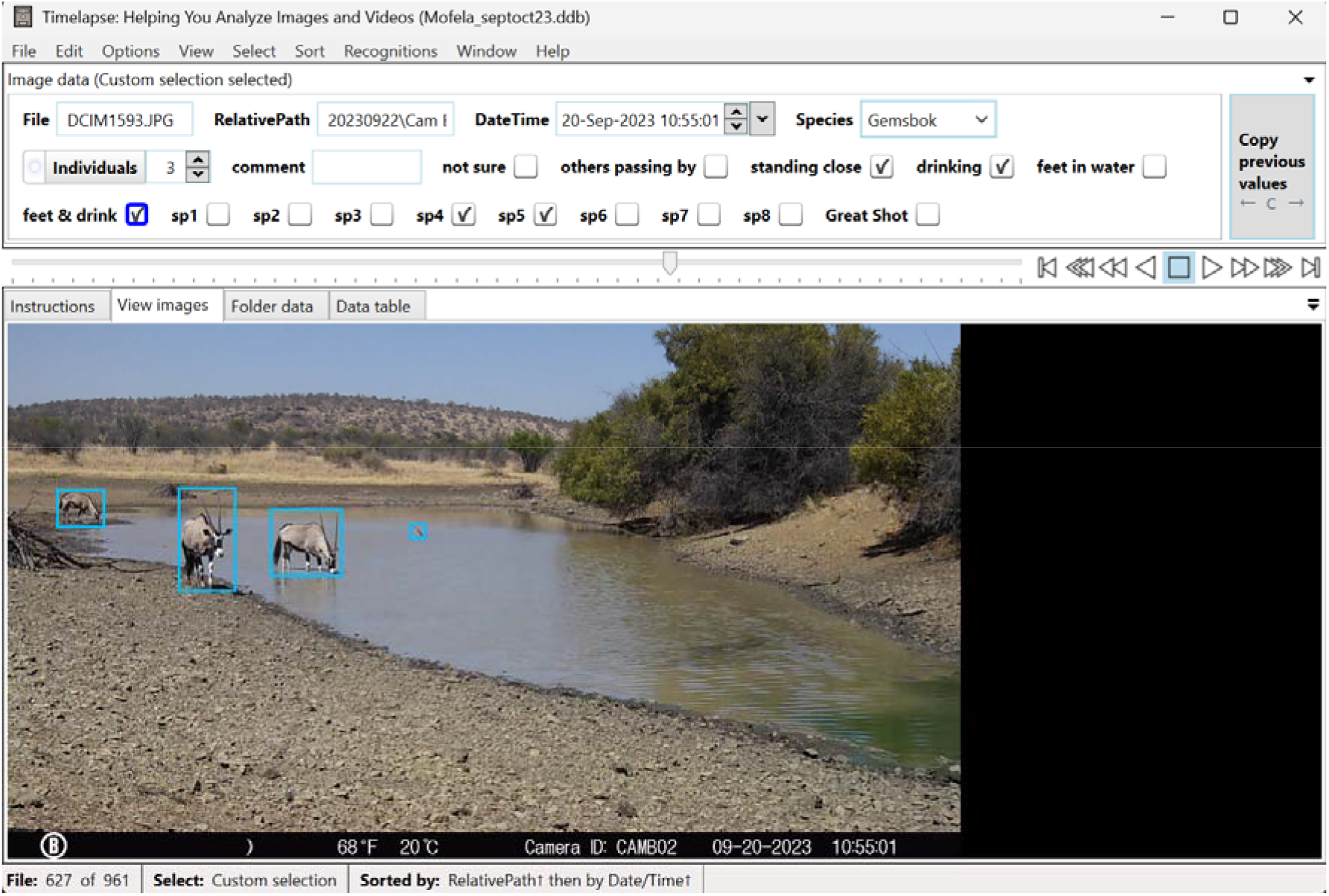
Screenshot of data processing in Timelapse (Greenberg et al., 2019). Each picture in which MegaDetector scored an animal to be present (blue rectangles) was scored for the following: Species (Gemsbok), number of individuals (3), water interaction type (standing close, drinking, feet & drink) and the closest sampling point(s) (sp4 & sp5).

The dams became progressively smaller during the dry periods during the course of the study affecting the site data that was analyzed. On 24.09.2023, Mofela dam became small enough to be covered by one single camera trap (CamB2). Consequently, the analysis of CamB2 was prioritized and the images from other cameras were only analyzed if required (e.g., if CamB2’s field of view was obstructed). For the 16,165 images taken by cameras at wildlife trails, the following parameters were scored for each mammal detected: (1) mammal species (2) number of individuals (3) proximity to the trail in terms of whether the animal was (a) on the transect or (b) within 5 m of the transect.

#### eDNA metabarcoding

The eDNA metabarcoding data included 725 field samples (+ 153 control samples) from Schenekar et al. (2026). In addition, 19 field samples not analyzed in that initial dataset were later subsequently sequenced and processed using the same laboratory and bioinformatic workflow. A brief method overview is given here, with the detailed eDNA workflow described in Schenekar et al. (2026). Eight sampling points were distributed as evenly as possible along the shoreline of each waterhole. At each sampling point, one water (500 ml) and one sediment (3 ml) sample were taken in six-day intervals. In the wet season, each waterhole was sampled eight times. During the dry season, the number of sampling events had to be reduced due to low water levels (Kolobe—water 2×, sediment 5×; Mofela—water 5×, sediment 8×; Monong water and sediment 7×). Wildlife trail samples were collected along a 5 m transect at six evenly distributed points in 6-day intervals, resulting in a total of four sampling events in the dry season.

Water samples were filtered through a 3 μm MCE filter, 8 μM MCE filter or 20-μm nylon filter, depending on water turbidity. All samples were stored in Longmire’s lysis buffer (Longmire et al., 1997) at 8°C in the field for up to seven weeks until they were transported to the genetic laboratory where they were stored at −20°C until DNA extraction. DNA extractions and PCR setup were performed in a dedicated low-template DNA laboratory, using the DNeasy PowerSoil Pro Kit (Qiagen), following the protocol described by Schenekar et al. (2024). For metabarcoding library preparation we used the MiMammal primer set (Ushio et al., 2017), optimized for amplifying mammalian DNA, and (2) the 12SV5 primer set (Riaz et al., 2011), which targets vertebrate sequences. Four PCR replicates were conducted for each sample. The libraries were sequenced on an Illumina NovaSeq X platform using 150 bp paired-end reads, with a target output of 80 million reads per library (approximately 417,000 reads per sample per primer set). Raw reads were quality-checked, demultiplexed, length filtered and dereplicated using cutadapt (Martin, 2011) and vsearch (Rognes et al., 2016). Taxonomic assignment was performed using the MIDORI2 database (Leray et al., 2022) with vsearch cutoff = 0.9, and ASVs assigned to the same species were merged for downstream analyses. For each sample, species detections from the two primer pairs were merged. During each sampling event at a waterhole, pH, conductivity, and dissolved oxygen levels were measured using a 500 ml measuring cup and a multimeter (pHenomenal Multimeter MU6100 H). A datalogger (OnsetHOBO Pendant MX Temperature/Light Data Logger) was placed in each waterhole to log temperature and light intensity at 5-min intervals.

#### Data preparation for statistical modelling – camera trap data

All camera-trap observations, defined as a single image containing at least one individual of a mammal species, were exported from Timelapse. When a species was recorded at multiple sampling points within the same observation, the record was duplicated such that each duplicate corresponded to one sampling point. The total number of individuals in the original observation was then divided by the number of sampling points checked, thereby assuming an equal distribution of individuals across the checked sampling points. This assumption is a simplification of the true spatial distribution, but it was necessary given how the camera-trap data were recorded. Similarly, if multiple behaviors were recorded for the same species within a single observation, the record was duplicated so that each duplicate corresponded to one sampling point behavior. The number of individuals per sampling point (as derived in the previous step) was then divided by the number of behaviors recorded, yielding for each record the sampling point and recorded behavior of each species and the “number of individuals per sampling point and behavior”. When the same species, sampling point, and behavior were recorded simultaneously by multiple cameras at the same waterhole, only one record was retained. Specifically, we kept the record with the highest number of observed individuals, assuming that this camera had the least obstructed view of the sampling point. If individual counts were tied, we retained the record with the greatest number of interaction types scored. If a tie still remained, one of these identical records was selected at random. Each camera-trap observation was assigned to an “observation period.” An observation period was defined as the six days preceding an eDNA sampling event at a given location, that is, the interval between the previous eDNA sampling and the current one. Observation periods were classified as regular when camera-trap coverage was available for a full six days interval and the eDNA dataset comprised eight water samples and eight sediment samples. Observation periods were classified as non-regular if (1) camera traps were active for less than six days before to the eDNA sampling event, (2) data were collected on trails, where only soil samples were taken or (3) only sediment or no eDNA samples were available for this period. Several dry season observation periods were excluded because only sediment or no eDNA samples could be collected due to the waterholes drying up. After these exclusions, a total of 34 regular observation periods were retained for the final analyses (Table 2).

**Table 2:** Number of observation periods for which data were collected (# Obs. periods) and number of regular observation periods retained for statistical analyses (# Regular obs. periods”). Fewer data were available during the dry period because several waterholes dried up during fieldwork.

| Location | Season | # Obs. periods | # Regular obs. periods |
| --- | --- | --- | --- |
| Dam 3 | wet | 8 | 7 |
| Kolobe | wet | 8 | 7 |
| Mofela | wet | 8 | 7 |
| Kolobe | dry | 4 | 2 |
| Mofela | dry | 6 | 5 |
| Monong | dry | 7 | 6 |
| Trail 1 | dry | 4 | 0 |
| Trail 2 | dry | 4 | 0 |
| Trail 3 | dry | 4 | 0 |
| Total |  | 53 | 34 |

“Visitation events” were defined as: All cameralZItrap observations of the same species from the same location, separated by less than 30 minutes were grouped into a single visitation event. This 30lZIminute interval threshold followed the approach described in O’Brien et al. (2003) and Zimmermann & Rovero (2016).

#### Data preparation for statistical modelling – eDNA data

Read counts per sample and species were converted to binary species detections (presence/absence) at two levels: per sample and per sampling event. A sampling event was defined as all eDNA samples collected at a given location on a given day, with eDNA detections combined across these samples. For waterhole locations, each sampling event comprised 16 samples (eight sediment and eight water samples), whereas for trail locations, each sampling event comprised six soil samples. For the per–samplinglZIevent dataset, we calculated the number of samples in which each species was detected. The dataset was then restricted to mammal detections only, and the corresponding observation period was assigned to each sample and sampling event, linking each eDNA record to the relevant cameralZItrapping observation period. Detections of elephant and nyala were removed because they were presumed to originate from secondary DNA input (Schenekar et al., 2026). Savanna hare (Lepus capensis) detections were assigned to Lepus sp. (“hare”), as the cameralZItrap data could not reliably distinguish Savannah hare from scrub hare (Lepus saxatilis) and were treated as one species for downstream analysis.

#### Covariates/predictors for modelling

We included species-specific ecological traits (species traits), camera-trap-derived activity metrics and environmental parameters as predictors in our subsequent models.

Species traits: For each species, we compiled four ecological traits: (1) adult body mass, (2) adult head–body length, (3) social group size and (4) trophic level (1 – herbivore, 2-omnivore, 3 – carnivore). Primary data were obtained from the PanTHERIA database (Jones et al., 2009), and supplemented, where necessary, with information from GBIF (GBIF.org, 2025, accessed 23.03.2026), the Encyclopedia of Life (Parr et al., 2014), and species information from Kingdon’s Field Guide to African Mammals (Kingdon, 2015). Because the generalized linear models used in subsequent analyses cannot accommodate missing values, we created a binary indicator variable (5) Social group size missing, for cases where social group size could not be obtained for a species or a closely related species.

Camera trap-derived activity metrics: For each species and each observation period, we derived the following activity metrics from the camera-trap data: (1) Number of individual observations: calculated as the sum of “individuals per sampling point and behavior” across all images for a given species within an observation period. (2) Number of visitation events: calculated as the sum of visitation events for that species within that period. (3) Water interaction intensity: Water interaction behavior types were converted to ordinal scores as follows: standing close - 1, drinking - 2, feet in water – 3, and feet in water while drinking - 4. For each species and observation period, we calculated the 75% percentile of these behavioral scores across all images. This behavioral percentile served as an index of water interaction intensity per species and period, with higher values indicating greater water interaction (4) Time since last visit - same period: Defined as the time (elapsed in minutes) between the last recorded image of that species within this observation period and the eDNA sampling time point (fixed at 07:45 on the corresponding sampling day). (5) Time since last visit - overall: Defined as the time elapsed since the last recorded image of a species during the current or in any previous observation period, and the current eDNA sampling time point. (6) No previous visit - same period: Analogous to accommodating for missing data in the Social Group Size predictor, we created this binary indicator variable for cases where no image of a given species was recorded in this observation period. (7) No previous visit – overall. Analogous to the previous variable for cases where the given species had not been recorded at the waterhole at any time before to the eDNA sampling time point. (8) Water interaction missing: Analogous to the two previous variables, for cases where the given species had not been recorded in this period.

Environmental variables: In addition to species traits and camera-trap–derived metrics, we included a set of environmental variables in the models. (1) Season: A binary factor indicating whether the observation period occurred in the wet or dry season. (2) Temperature: The mean water temperature [°C] for the corresponding observation period and location, calculated from datalogger records. (3) pH: The water pH measured with a handheld multimeter (VWR) at the time of eDNA sampling. (4) Light intensity: The mean light intensity [lux] for the corresponding observation period and location, calculated from datalogger measurements. (5) Conductivity: Electrical conductivity of the water [total dissolved solids in mg/l], measured with a handheld multimeter at the time of eDNA sampling. (6) Oxygen concentration - Dissolved oxygen concentration [mbar] of the water, measured with a handheld multimeter at the time of eDNA sampling.

#### Statistical modelling

All continuous predictors were standardized (z-transformed) before statistical modelling. All analysis were conducted in R version 4.4.1 (R Core Team, 2023) using RStudio (Posit team, 2024). In addition to the below-mentioned packages, the following packages were used for data handling and visualization: dplyr (Wickham et al., 2023), readr (Wickham, Hester, et al., 2024), tidyr (Wickham, Vaughan, et al., 2024), lubridate (Grolemund & Wickham, 2011), stringr (Wickham, 2023) , eulerr (Larsson, 2024), purrr (Wickham & Henry, 2023) and ggplot2 (Wickham, 2016). All R scripts used in this study are available in Supplementary Material 2 (will be made available through a github repository upon publication).

#### Logistic regression of species traits on eDNA and camera trapping detection overall

To evaluate whether species-level traits influence detection probabilities across the entire dataset, we fitted separate logistic regression models for eDNA and camera-trap data. For both eDNA and camera-trap datasets, species-level detection (detected vs. not detected at least once during the study) was modelled as a function of these traits using a binomial logit model. Trophic level was included as a categorical predictor, while adult body mass, adult head–body length, and social group size were treated as continuous predictors. Logistic regression models were fitted using the glm function with a binomial error distribution and logit link function. We used type III analyses of deviance (ANOVA) to obtain overall tests for each predictor. Collinearity between adult body mass and adult head–body length was assessed using Pearson correlation coefficients and variance inflation factors (VIFs). As VIF values were low, both traits were retained in the final model. Model fit and predictive performance was evaluated with the Hosmer–Lemeshow goodness-of-fit test using the hoslem.test function in the ResourceSelection package of R (Lele et al., 2023) and receiver operating characteristic (ROC) curves with the area under the curve (AUC) values using the roc function of the pROC package (Robin et al., 2011). Differences in mean detection rates among trophic levels were analyzed using a Kruskal–Wallis test followed by Dunn’s post hoc comparisons with Holm-adjusted p-values.

#### Mixed-effects modelling of species traits, activity metrics and environmental variables on eDNA detection probability

For this model, only regular observation periods were included. We modelled how species traits, camera trap-derived activity metrics and environmental variables influenced eDNA detection probability at sampling events using a binomial generalized linear mixed-effects model with a logit link. The number of eDNA-positive samples for each species out of the total number of samples at a sampling event was used as the response variable. Fixed effects in the initial full model included the species traits adult body mass, adult head–body length, social group size, the indicator for social group size missing and trophic level. Camera trap-derived activity metrics included the number of individual observations, the number of visitation events, time since last visit – same period, the indicator of no previous visit – same period, time since last visit – overall, the indicator of no previous visit – overall, water interaction intensity and the indicator of water interaction intensity missing. Environmental variables included season, temperature, pH, light intensity, conductivity and oxygen concentration. Random effects included species, location (waterhole) and observation period. The model was fitted using the glmer function of the R package lme4 (Bates et al., 2015) using the bobyqa optimizer with an increased iteration limit to aid convergence. After running the full model, the missingness indicators for social group size and behavioral metrics, which were fully collinear with species, were removed. We then applied stepwise simplification of the fixed-effects structure, comparing nested models using likelihood-ratio tests and Akaike’s Information Criterion (AIC), sequentially evaluating the contribution of environmental variables, season, species traits (adult body mass and head–body length) and time since last visit within the same period. For the final model, we assessed goodness-of-fit using simulation-based residual diagnostics from the DHARMa package (Hartig, 2024), including tests for overdispersion and zero inflation.

#### Mixed-effects modelling incorporating substrate (water versus sediment)

To test whether the effects of the covariates identified in the final model differed between substrates, we refitted the binomial GLMM on a dataset in which each sampling event was split by medium and a factor medium was added to indicate substrate type. Starting from the final model structure of the previous modelling, we introduced medium as a fixed effect and allowed it to interact with all biological, activity, and environmental predictors. This extended model, enabled explicit assessment of whether eDNA detection patterns differed between water and sediment samples, while retaining the same random-effects structure as in the original model. We also applied a stepwise simplification of the fixed-effects structure as described above. However, the ANOVA showed a significant reduction in model fit of the reduced model such that the full model was kept.

#### Spatial mixed-effects modelling on eDNA detection at the individual sample level

To investigate whether eDNA detection at a given sampling position was driven by the fine-scale spatial distribution of eDNA resulting from localized water interactions within waterholes, we extended the initial mixed-effects model to the level of individual samples and explicitly incorporated spatial predictors. We used a binomial generalized linear mixed-effects model with a logit link, again fitted with the glmer function of the package lme4 and retained the same structure of fixed and random effects as in the previous full model (species traits, camera trap-derived activity metrics and environmental variables). Additionally, we included the following spatially explicit covariates: For each eDNA sample and species, we derived counts of water interactions of that species at the same sampling spot, and its neighboring sampling spots. Sampling spots were arranged in a circular layout around each waterhole (eight positions, see Figure 1), and for each sample we calculated (1) “number of same-spot interactions”, which is the number of recorded water interactions of the focal species at that specific spot in the corresponding observation period, (2) “number of neighboring spots interactions”, which were the number of water interactions at the two neighboring sampling points and (3) “number of local interactions”, which was the sum of same-spot and neighboring spot interactions. We first fitted a base model without spatial terms, corresponding to the earlier detection model but using the binary detection response at the sample level. We then fitted a spatial model that included also “number of same-spot interactions” and “number of local interactions” as predictors. Both models were fitted with a binomial error distribution and logit link, again using bobyqa optimizer. To assess the added value of the spatial covariates, we compared the base and spatial models using a likelihood-ratio test. Model fit and residual structure were also evaluated using simulation-based diagnostics from the DHARMa package.

#### Mixed-effects modelling of soil samples on trails

We assessed the effects of species traits and activity metrics on eDNA detection rates using the data from soil samples collected on wildlife trails during the dry season. Because no environmental covariates were measured for this dataset, they were not included in the model. Instead of a behavioral percentile, we calculated a “proximity percentile” analogous to the behavioral metric used for waterholes, based on three ordinal levels of scored proximity for each observation (3 = on trail transect, 2 = within 5 m of transect, and 1 = farther than 5 m from transect). The model structure otherwise followed the mixed-effects model used for eDNA detection probability at waterholes. The initial full model included species traits (adult head–body length, adult body mass, social group size, an indicator for missing social group size, trophic level) and activity metrics (number of individual observations, number of visitation events, time since last visit within the same period, time since last visit overall, proximity percentile, and an indicator for missing proximity percentile) as fixed effects, and species, location, and observation period as random intercepts. We first fitted this full model and then simplified it by removing statistically weak and/or collinear predictors. Full and reduced models were compared using likelihood-ratio tests, and the final model was checked for overdispersion and zero inflation using DHARMa residual diagnostics.

## 3. Results

### Species detections from camera trapping and eDNA data sets

Across both seasons and all waterholes (including trails), the camera traps recorded a total of 1,121,085 images over 749.8 net camera-trap days, of which 92,667 contained mammal observations (Table 1). From these images, 30 mammal species were identified. From the 744 field samples in the eDNA dataset (320 water, 352 sediment and 72 soil samples), 40 mammal species were detected. Overall, unique 45 mammal species belonging to 10 orders and 19 families were detected across both datasets, of which 25 were shared between the camera-trap and eDNA datasets (Figure 3, Table 3).

**Figure 3:**
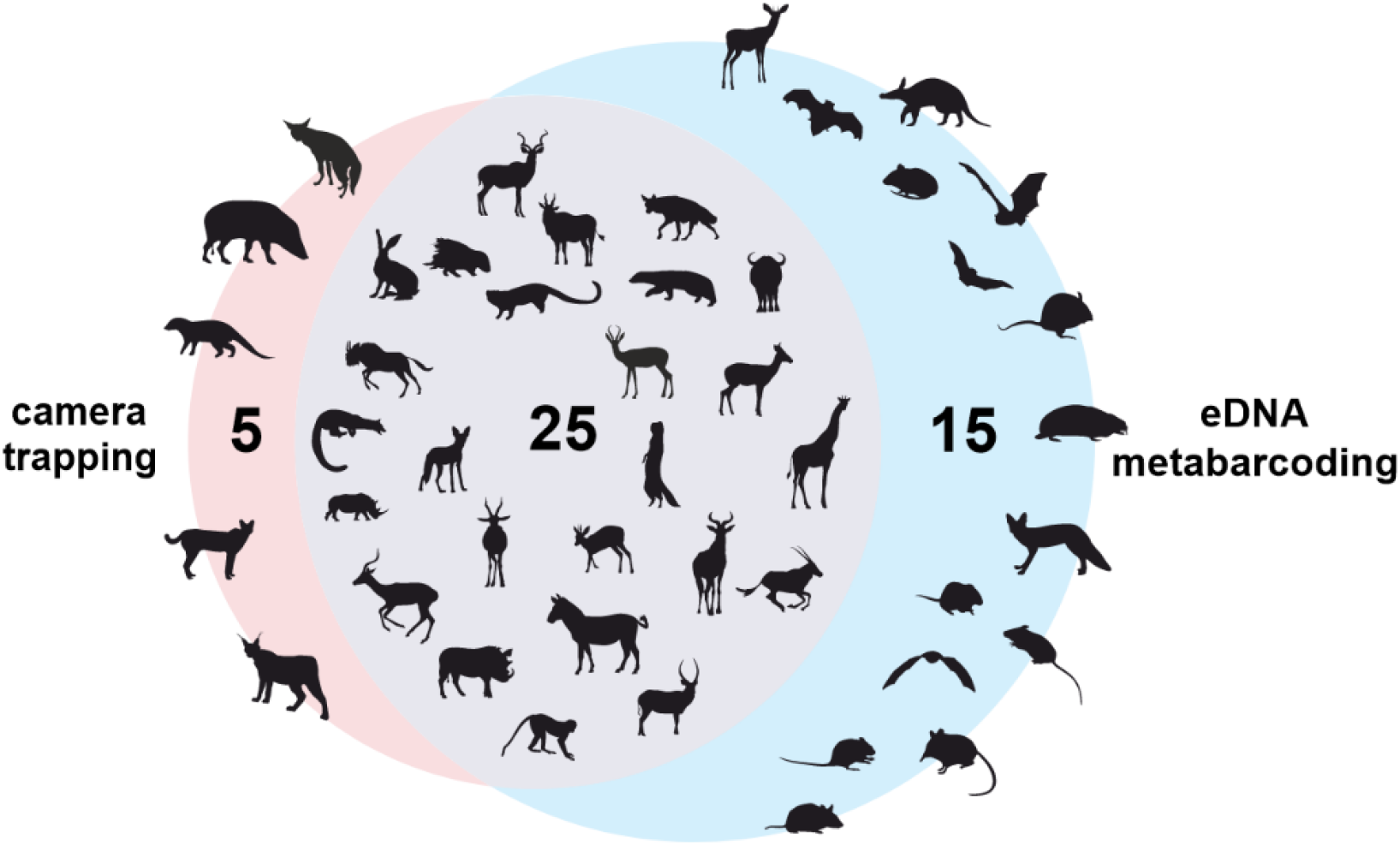
Euler diagram of the detected mammal species by the two monitoring approaches (camera trapping and eDNA metabarcoding). Numbers indicate the detected species numbers by the respective approach.

**Table 3:** Detections (✓) and non-detections (-) of the 45 mammal species by the two monitoring approaches camera trapping (CT) and eDNA metabarcoding (eDNA). For each species, the scientific and common name, as well as respective order and family are given.

| Species | Common name | Order | Family | CT | eDNA |
| --- | --- | --- | --- | --- | --- |
| *Aepyceros melampus* | impala | Artiodactyla | Bovidae | ✓ | ✓ |
| *Aethomys chrysophilus* | red rock rat | Rodentia | Muridae | - | ✓ |
| *Alcelaphus buselaphus* | red hartebeest | Artiodactyla | Bovidae | ✓ | ✓ |
| *Antidorcas marsupialis* | springbok | Artiodactyla | Bovidae | ✓ | ✓ |
| *Caracal caracal* | caracal | Carnivora | Felidae | ✓ | - |
| *Ceratotherium simum* | white rhinoceros | Perissodactyla | Rhinocerotidae | ✓ | ✓ |
| *Chlorocebus pygerythrus* | vervet monkey | Primates | Cercopithecidae | ✓ | ✓ |
| *Connochaetes gnou* | black wildebeest | Artiodactyla | Bovidae | ✓ | ✓ |
| *Cryptomys holosericeus* | common mole-rat | Rodentia | Muridae | - | ✓ |
| *Cynictis penicillata* | yellow mongoose | Carnivora | Herpestidae | ✓ | ✓ |
| *Damaliscus pygargus* | blesbok | Artiodactyla | Bovidae | ✓ | ✓ |
| *Elephantulus brachyrhynchus* | short-snouted elephant shrew | Macroscelidea | Macroscelididae | - | ✓ |
| *Equus quagga* | plains zebra | Perissodactyla | Equidae | ✓ | ✓ |
| *Galerella sanguinea* | slender mongoose | Carnivora | Herpestidae | ✓ | ✓ |
| *Genetta genetta* | small-spotted genet | Carnivora | Viverridae | ✓ | ✓ |
| *Gerbilliscus leucogaster* | bushveld gerbil | Rodentia | Muridae |  | ✓ |
| *Giraffa giraffa* | giraffe | Artiodactyla | Giraffidae | ✓ | ✓ |
| *Hystrix africaeaustralis* | cape porcupine | Rodentia | Hystricidae | ✓ | ✓ |
| *Kobus ellipsiprymnus* | waterbuck | Artiodactyla | Bovidae | ✓ | ✓ |
| *Laephotis capensis* | cape serotine | Chiroptera | Vespertilionidae | - | ✓ |
| *Lemniscomys rosalia* | single-striped grass mouse | Rodentia | Muridae | - | ✓ |
| *Leptailurus serval* | serval | Carnivora | Felidae | ✓ | - |
| *Lepus sp* | hare | Lagomorpha | Leporidae | ✓ | ✓ |
| *Lupulella mesomelas* | black-backed jackal | Carnivora | Canidae | ✓ | ✓ |
| *Mastomys coucha* | southern multimammate mouse | Rodentia | Muridae | - | ✓ |
| *Mellivora capensis* | honey badger | Carnivora | Mustelidae | ✓ | ✓ |
| *Miniopterus fraterculus* | lesser long-fingered bat | Chiroptera | Miniopteridae | - | ✓ |
| *Miniopterus natalensis* | Natal long-fingered bat | Chiroptera | Miniopteridae | - | ✓ |
| *Mungos mungo* | banded mongoose | Carnivora | Herpestidae | ✓ | - |
| *Mus minutoides* | southern African pygmy mouse | Rodentia | Muridae | - | ✓ |
| *Orycteropus afer* | aardvark | Tubulidentata | Orycteropodidae | - | ✓ |
| *Oryx gazella* | gemsbok | Artiodactyla | Bovidae | ✓ | ✓ |
| *Otomys angoniensis* | Angoni vlei rat | Rodentia | Muridae | - | ✓ |
| *Parahyaena brunnea* | brown hyena | Carnivora | Hyaenidae | ✓ | ✓ |
| *Phacochoerus africanus* | common warthog | Artiodactyla | Suidae | ✓ | ✓ |
| *Potamochoerus larvatus* | bushpig | Artiodactyla | Suidae | ✓ | - |
| *Proteles cristatus* | aardwolf | Carnivora | Hyaenidae | ✓ | - |
| *Raphicerus campestris* | steenbok | Artiodactyla | Bovidae | - | ✓ |
| *Redunca fulvorufula* | mountain reedbuck | Artiodactyla | Bovidae | ✓ | ✓ |
| *Scotophilus viridis* | greenish yellow bat | Chiroptera | Vespertilionidae | - | ✓ |
| *Sylvicapra grimmia* | bush duiker | Artiodactyla | Bovidae | ✓ | ✓ |
| *Syncerus caffer* | African buffalo | Artiodactyla | Bovidae | ✓ | ✓ |
| *Tragelaphus oryx* | eland | Artiodactyla | Bovidae | ✓ | ✓ |
| *Tragelaphus strepsiceros* | greater kudu | Artiodactyla | Bovidae | ✓ | ✓ |
| *Vulpes chama* | Cape fox | Carnivora | Canidae | - | ✓ |

### Logistic regression of species traits on eDNA and camera trapping detection overall

For camera trapping, logistic regression revealed adult head-body length as the only species trait to have a clear association with detection, with longer-bodied species having a higher probability of being detected by camera traps (Table 4). For eDNA, trophic level was a significant predictor, with ANOVA indicating a strong overall effect of trophic level on eDNA detection probabilities. Both logistic regression models showed adequate fit according to the Hosmer–Lemeshow test (camera trapping: χ^2^ = 7.97, df = 8, P = 0.44; eDNA: χ^2^ = 0.91, df=7, P = 1.00) and high discriminatory ability (camera trapping: AUC = 0.896; eDNA: AUC = 0.964). Kruskal-Wallis test showed significant differences among trophic levels χ² = 14.85, p < 0.001), and post-hoc showed that all pairwise significant differences between herbivores and omnivores and herbivores and carnivores, but not between omnivores and carnivores.

**Table 4:** Results of type III ANOVA on logistic regression models testing the effects of species traits on camera-trap and eDNA detection. LR χ^2^ : likelihood-ratio statistic; df: degrees of freedom. * indicate a P-value < 0.05

| Monitoring approach | Species trait | LR $\chi^2$ | df | P |
| --- | --- | --- | --- | --- |
| Camera trapping | Adult body mass | 0.09 | 1 | 0.764 |
|  | Adult head-body length | 5.87 | 1 | 0.015* |
|  | Social group size | 1.96 | 1 | 0.161 |
|  | Trophic level | 1.73 | 2 | 0.421 |
| eDNA metabarcoding | Adult body mass | 0.88 | 1 | 0.348 |
|  | Adult head-body length | 2.10 | 1 | 0.148 |
|  | Social group size | 0.72 | 1 | 0.396 |
|  | Trophic level | 14.47 | 2 | 0.0007* |

### Mixed-effects modelling of species traits, activity metrics and environmental variables on eDNA detection probability

At the level of sampling events, eDNA detection probability was strongly associated with a subset of the species traits and camera trap-derived activity metrics, whereas environmental variables and season did not improve model fit (Table 5). In the final binomial mixed-effects logistic regression model, the probability that a species was detected by eDNA increased with social group size, number of visitation events, the number of individual observations and water interaction intensity. Trophic level showed a strong negative association with eDNA detection. Relative to herbivores, omnivores and carnivores had lower detection probabilities. Furthermore, detection probability decreased with increasing time since the last visit across all periods and was lower when no previous visit had ever been recorded overall. Counterintuitively, the model also predicted a higher detection probability when no visit had occurred based on camera trap data in the current period. In contrast, adult body mass, adult head–body length, season, and all environmental variables (temperature, pH, light intensity, conductivity, and oxygen concentration) did not significantly improve model fit and were excluded during stepwise simplification based on likelihood-ratio tests and AIC. Random intercepts captured substantial among-species variation in baseline detection probability, with smaller contributions of period and location. DHARMa residual diagnostics indicated no evidence of overdispersion (dispersion = 0.89) or zero inflation (ratio of observed to expected zeros = 0.97), suggesting that the final model provided an adequate description of the data (for DHARMa diagnostic plots, see Supporting Information 1, Figs S1-S3).

**Table 5:** Summary of fixed effects in the final binomial mixedlZIeffects model of eDNA detection probability at sampling events. For each predictor, the estimate (on logit scale), the standard error of the estimated coefficient (Std. Error), Wald test statistic (z-value) and P-value are given. *: P < 0.05, **: P < 0.01, ***: P < 0.001

| Predictor | Estimate | Std. Error | z-value | P |
| --- | --- | --- | --- | --- |
| (Intercept) | -2.94 | 0.44 | -6.66 | 0.001*** |
| Social group size | 0.92 | 0.28 | 3.30 | 0.0009*** |
| Trophic level 2 (omnivore) | -2.29 | 0.69 | -3.29 | 0.001** |
| Trophic level 3 (carnivore) | -2.65 | 0.80 | -3.32 | 0.0009*** |
| Number of visitation events | 0.12 | 0.03 | 3.75 | 0.0002*** |
| Number of individual observations | 0.05 | 0.02 | 2.09 | 0.036** |
| time since last visit - overall | -0.35 | 0.07 | -4.99 | <0.0001*** |
| No previous visit - same period | 0.78 | 0.20 | 3.83 | 0.0001*** |
| No previous visit - overall | -1.34 | 0.26 | -5.20 | <0.0001*** |
| Water interaction intensity | 0.12 | 0.04 | 2.86 | 0.004** |

In summary, the final model indicates that variation in detection probability is primarily driven by species’ social and trophic characteristics, water interaction intensity and number of interactions, rather than by environmental covariates or body-size traits.

### Mixed-effects modelling incorporating substrate (water versus sediment)

In the extended model including substrate we found a strong effect of medium on eDNA detection and clear substrate-specific differences in how species and environmental predictors influence detection (Table S2). Overall, the eDNA detection probability was significantly lower in water than in sediment samples. Several key predictors showed significant interactions with substrate type. Concerning species traits, while social group size and low trophic level generally had a positive effect on detection probability, these relationships were stronger in sediment than in water samples that were collected. In addition, body size showed a more positive association with detections in water versus sediment. In terms of activity metrics, the number of observed individuals and their water interaction had significantly stronger positive effects on detection probabilities in water samples. Finally, environmental variables also differed between the substrates: temperature and dissolved oxygen were positively associated with detection in water, whereas higher pH, conductivity and light intensity reduced water detection probability relative to sediment samples. Model checking with DHARMa indicated adequate fit and no evidence of problematic zero inflation (for DHARMa diagnostic plots, see Supporting Information 1, Figs S4-S6).

### Spatial mixed-effects modelling on eDNA detection at the individual sample level

At the level of individual eDNA samples, the base mixed-effects model without spatial terms indicated that eDNA detection probability increased with social group size and several camera trap-derived activity metrics, and decreased with trophic level and time since last visit, largely mirroring the results of the models applied at the sampling event level (Supporting information 1, Tables S1 & S2). Adding the spatial covariates significantly improved model fit (likelihood-ratio test: χ^2^ = 29.486, df = 2, P < 0.001, AIC_no_spatial_= 10472, AIC_spatial_ = 10446), indicating that within-waterhole spatial structure in eDNA deposition is non-random. In the spatial model, same-spot camera trap activity had a strong positive effect on detection probability (z = 3.48, P < 0.001, Supporting information 1, Table S3), while the local interaction activity was not significant. Other significant effects in the spatial model largely mirrored those of the base model. DHARMa diagnostics did not indicate substantial overdispersion or zero inflation for either model (Supporting information 1, Figs S7-S12).

### Mixed-effects modelling of soil samples on trails

This model revealed that eDNA detection rate from soil samples on wildlife trails was strongly associated with certain species traits and selected camera-trap activity metrics (Table 6). In the final mixed-effects model, adult head–body length had a positive effect on detection probability, indicating that larger species were more likely to be detected. Trophic level also influenced detection, with carnivores showing markedly lower detection probabilities compared to herbivores. Among the activity metrics, the number of visitation events had a positive effect on detection probability, and the proximity percentile was positively associated with detection, suggesting that species whose observations were more frequently recorded directly on or very close to the trail were more likely to yield detectable eDNA. Social group size and time since last visit did not show clear effects on detection. Random intercepts for species, location, and observation period indicated additional unexplained variation at these levels, but model diagnostics (DHARMa) confirmed an adequate fit without evidence of overdispersion or zero-inflation (for DHARMa diagnostic plots, see Supporting Information 1, Figs S13-15).

**Table 6:** Summary of fixed effects in the final mixedlZIeffects model of eDNA detection probability at sampling events of soil samples from wildlife trails. For each predictor, the estimate (on logit scale), the standard error of the estimated coefficient (Std. Error), the Wald test statistic (z-value) and P-values (P) are given. *: P < 0.05, **: P < 0.01, ***: P < 0.001

| Predictor | Estimate | Std. Error | z-value | P |
| --- | --- | --- | --- | --- |
| (Intercept) | -2.75 | 0.59 | -4.68 | 0.000003*** |
| Adult head-body length | 2.27 | 0.50 | 4.53 | 0.000006*** |
| Social group size | -0.46 | 0.40 | -1.15 | 0.25 |
| Trophic level 2 (omnivore) | -0.17 | 0.87 | -0.20 | 0.84 |
| Trophic level 3 (carnivore) | -4.61 | 1.60 | -2.88 | 0.004** |
| Number of visitation events | 0.17 | 0.08 | 2.13 | 0.033* |
| Time since last visit – same period | -0.01 | 0.07 | -0.16 | 0.88 |
| Proximity percentile | 0.26 | 0.10 | 2.53 | 0.011* |

## 4. Discussion

This study conducted extensive monitoring using camera trapping and eDNA metabarcoding to assess the effect of species traits, activity metrics and environmental variables on eDNA detection probabilities. To our knowledge Harper et al. (2019) is the only similar study directly linking terrestrial mammal behavior to eDNA detection in water bodies though the joint analysis of these two monitoring approaches. In contrast to their findings, we found eDNA detection probability rising with water interaction intensity, particularly in water samples. While the effect of behavior was strong on water detections, it was weaker and not statistically significant in sediment. This result suggests water samples in semi-arid areas are more sensitive to the extent of more recent water interactions of mammals. Harper et al. (2019) suggested that the artificial pond in the British Zoo may have been saturated with eDNA, masking any effect of animal behavior. Consequently, we hypothesize that the effect was visible in our study predominantly in the water samples because the animals occurred in lower densities in the 5800-ha study area. Whilst the largest waterbody in Harper et al. (2019) had a size of 100 m^2^, our estimations of the surveyed waterholes are mostly between 1000 m^2^ and 6000 m^2^ (though strongly fluctuating throughout the seasons). Moreover or alternatively, DNA degradation dynamics are likely to be accelerated due to higher temperatures that typify the subtropical climate of our study (Mas-Carrió et al., 2022) and our study system featured a higher species diversity. However, as deposited eDNA reportedly persists longer in sediment than in water samples (Sedlmayr & Schenekar, 2024), a weak, but similar saturation effect might have caused the weaker effect of behavior in the sediment samples.

### Sensitivity of camera trapping and eDNA metabarcoding and species traits affecting detection by either method

There was a significant overlap of the species detected by the two methods, even though also a substantial number of species were only detected by one or the other. As reported in earlier work (Leempoel et al., 2020), body size measured in adult head-body length was a significant predictor for detection via camera trapping whilst eDNA was also able to detect small and elusive animals missed by cameras.

Trophic level was a significant species trait for species detection across the eDNA dataset, with several carnivores and omnivores not being detected by the eDNA dataset but by camera trapping, potentially due to their lower occurrences or due to their limited water interactions compared to herbivores. This corroborates the recommendations of previous studies, which suggest that eDNA and camera trapping are complementary approaches, with mixed or approaches showing the highest sensitivity in terms of the number of species detected (Leempoel et al., 2020; Mas-Carrió et al., 2022). Given the extensive eDNA and camera-trapping datasets in this study, this pattern is unlikely to be a mere sampling artefact and instead appears to be robust even when based on comparatively exhaustive data.

### Species-specific traits affecting eDNA detection probability

Of the tested species-specific traits, social group size had the largest effect on eDNA detection probability. This partly reflects its correlation with a higher probability of observing more individuals of a more social species. While the applied GLMMs can accommodate correlation among predictors, the moderately high correlation between social group size and the number of individual observations (r = 0.72) in our dataset could have led to a small and less precisely estimated unique effect of social group size after adjusting for observation frequency. Consistent with this, previous work has shown that the number of individuals interacting with a water body influences eDNA shedding rates and consequently detection probability (Williams et al., 2018).

Although individual body mass and body volume are suggested as indicators of eDNA detection probability many times (Barnes & Turner, 2016; Thomsen & Willerslev, 2015), our findings showed no effect of either traits. Conversely, many studies indicated biomass as a significant predictor, be it airborne (Lynggaard et al., 2022), in lentic systems (Mas-Carrió et al., 2022; Yates et al., 2025) or in streams (Lyet et al., 2021). There are unfortunately no available estimations of the number of individuals of any species present in Botsalano game reserve to include the factor “biomass” in our model as a predictor. It seems probable, however, that the total biomass of a species, which is expected to be high for the social antelopes, may contribute to the large effect of social group size as the most significant predictor.

Trophic level was a strong predictor of eDNA detection probability, with carnivores and omnivores having a substantially lower detection rate than herbivores. Carnivores usually occur in lower abundances at waterholes. Consequently, the reduced eDNA detection rates observed for carnivores may partly reflect their lower overall biomass in the study area.

Of the 11 species of the Order Carnivora, four were not detected by eDNA metabarcoding, including both felids (Caracal Caracal caracal and Serval Leptailurus serval). Previous related studies also report lower detectability of felids with eDNA methods (Bai et al., 2025; Broadhurst et al., 2021; Lyet et al., 2021). In addition to lower abundances and reduced water interactions of carnivores and omnivores relative to herbivores - in our study herbivores stayed on average longer at the waterhole and had more intense water interactions than omnivores and carnivores. Additional behavioral differences that we did not score, like defecation at the waterhole, may have further contributed to these differences as defecation is a significant source of eDNA in such systems (Sauseng & Schenekar, 2025). The large effect of species as a random effect further suggests that, aside from the species-specific traits considered in our models, there might be additional species-specific traits affecting eDNA detection.

### Activity metrics-influence of mammal behavior on eDNA detection probability

Several camera-trap derived activity metrics showed a strong influence on eDNA detection probability. The positive effect of water interaction intensity corroborates the few existing studies that explicitly link behavior to eDNA deposition (Harper et al., 2019; Sauseng & Schenekar, 2025). Additionally, both the number of visitation events and the total number of individuals observed increased detection probability positively. This is in line with the mechanistic expectations and matches results from others that report higher detection probabilities with more observed individuals (Harper et al., 2019; Lyet et al., 2021). The strong effect of “time since last visit overall” is in line with eDNA degradation reducing detection probabilities of species as the time span between eDNA deposition and eDNA sampling increases (Barnes & Turner, 2016; Buxton et al., 2017; Strickler et al., 2015). Overall, our findings corroborate the view that integrating behavioral and activity data from camera traps is crucial for interpreting eDNA signals and for understanding false non-detections in terrestrial mammal monitoring

### Environmental factors

In contrast to species traits and activity metrics, environmental variables showed little contribution to explaining variation in eDNA detection probabilities in our study. Water temperature, conductivity, dissolved oxygen, pH, light intensity, and season did not improve model fit. This suggests that, within the environmental gradients encompassed by our study, physico–chemical conditions were relatively minor and did not strongly affect eDNA persistence. Previous work has demonstrated that temperature, UV exposure, and pH can markedly influence eDNA degradation rates and detection probabilities, particularly along steep environmental gradients or under extreme conditions (McCartin et al., 2022; Strickler et al., 2015). However, in our study, the variation in eDNA detection appears more strongly driven by species ecology, activity and occupancy rather than by moderate fluctuations in abiotic conditions. However, individual environmental variables did have an effect in the model, accounting for substrate type (see below).

### Differing effects of species traits, activity metrics and environmental factors in sediment and water samples

The inclusion of substrate type (water vs. sediment) revealed that eDNA detection probabilities differ markedly between them and that the influence of species traits, activity metrics, and environmental variables is substrate-dependent. Overall, detection probabilities were substantially lower in water than in sediment. This pattern is consistent with the notion that sediments act as a longerlZIterm sink and integrator of eDNA, whereas water samples capture more transient or recent signals (Turner et al., 2015), also a previous observation from the same study site (Schenekar et al., 2026). Additionally, several significant interaction terms between substrate and multiple predictors demonstrate that water and sediment respond differently to biological and environmental drivers. For example, the positive main effects of social group size and low trophic level (herbivores) on detection are much stronger in sediment, while their interactions with water are strongly negative, indicating that the positive effect of sociality and herbivory on eDNA detection probability is stronger in the water column. The interactions between activity metrics and substrate further support the suggestion that water and sediment integrate mammal activity over different temporal scales. In sediment, detection is already positively related to the number of visitation events and negatively to the time since last visit, consistent with sediment acting as a temporal integrator. In water, the additional positive interaction for individual observations suggests that recent high-intensity activity is particularly important to generate reliable eDNA detection signals in the water column. Additionally, the positive effect of the interaction of substrate water and high water interaction intensity indicates that behaviors such as drinking, or submerging have a disproportionate effect on water eDNA detection compared to sediment detection. These results echo experimental work showing that behaviors involving direct contact with water can strongly enhance detection in water samples (Sauseng & Schenekar, 2025). Finally, the environmental variables showed pronounced substratelZIdependent effects. Several abiotic predictors had relatively weak or inconsistent main effects, but their interactions with substrate were often strong and highly significant. In sediment, light intensity showed a modest but significant positive association with detection, whereas in water it was strongly and negatively related to detection probability. The latter is consistent with experimental evidence that UV and high light accelerate degradation of free eDNA in the water column, but at this point we cannot explain the presumable positive effect of light in sediment samples, aside from the fact that sediment-bound DNA may be shielded from direct radiation damage, reducing the negative effect of UV light. The pH, conductivity and dissolved oxygen levels likewise had weak main effects in sediment but strong, often opposing, interaction terms with water, indicating that these variables affect eDNA persistence more strongly in the water column than in sediments. Overall, these patterns suggest that abiotic drivers primarily shape waterlZIcolumn eDNA signals, while sediment signals are comparatively buffered and more strongly driven by cumulative biological inputs from species traits and activity.

### Spatial distribution of eDNA detection signals

The inclusion of spatial predictors significantly improved model fit, indicating that eDNA detection is spatially structured within waterholes rather than homogeneous. Specifically, same-spot interactions had a significant effect, whereas local nearby interactions did not, a pattern suggesting that eDNA signals in our lentic systems are strongly localized. Detection probability was highest where animals directly interact with the substrate and drops off over very short distances. Such fine-scale spatial structuring is consistent with earlier work reporting patchy eDNA distributions in lentic systems (Harper et al., 2019; Sales et al., 2020). From a practical perspective, these results highlight that location of sampling within a lentic waterbody critically affects which species are detected, and that sampling designs aiming to maximize detection should either target known activity hotspots or include spatially replicated sampling to capture smalllZIscale heterogeneity in eDNA distribution.

### Detection probabilities at wildlife trails

Detection probability increased with higher visitation frequency and closer proximity to the wildlife trail, indicating that species whose observations were more often recorded directly on or very close to the trail were more likely to leave detectable eDNA in the sampled soil. In contrast to our waterhole model, larger species were more likely to be detected. Trails are usually formed by large herbivores and frequented by predator species (Neave et al., 2025). It is likely that smaller and cryptic species, such as mice and mongooses, are using the trails less due to higher exposure and predation risk. This aligns with the general assumption that local space use and proximity to sampled substrates strongly shape eDNA signals in terrestrial settings (Leempoel et al., 2020). As in the waterhole models, carnivores showed lower detection probabilities than herbivores. Together, these soil-based results reinforce our conclusion that species traits and fine-scale activity patterns are primary drivers of terrestrial mammal eDNA detection across substrates.

## 5. Conclusions

By combining extensive camera trapping with eDNA metabarcoding in waterholes in natural environments, we show that mammal eDNA detection is strongly structured by species traits, behavior and substrate type, with relatively weak, but substrate-dependent, effects of environmental conditions. Overall, our results demonstrate a significant influence of mammalian social and individual behavior on eDNA detectability, alongside previously reported drivers such as the number of individuals and the time elapsed between water interaction and eDNA sampling. Additionally, we also corroborated previous work that showed higher eDNA persistence in sediment over water samples, with eDNA in sediments being less susceptible to negative abiotic influences. From a monitoring perspective, this implies that water eDNA provides a high-resolution but short-memory record, whereas sediment eDNA can better reflect chronic space use and longer-term presence of species. Finally, the spatial heterogeneity of eDNA detections at individual sampling spots underscores the fine-scale spatial structure of eDNA in lentic systems and the significant value of sampling close to observed animal activity and/or spatial replication in eDNA sampling.

## Acknowledgments

We would like to thank the management of Botsalano Game Reserve, especially T. Ngubenkomo and D. Dinale for their support of this research and North West Parks Board, Particularly J. Power, and the Department of Economic Development, Environment, Conservation and Tourism, South Africa for providing the necessary permits. Sampling for the project has been approved by North West Parks Board and by the Department of Economic Development, Environment, Conservation and Tourism, South Africa (Permit No. NW 39881/05/2022), the research has been approved by SANBIs NZG Animal Research Ethics and Scientific Committee (P2022/15) and a permission in terms of section 20 of the animal diseases act 1984 was received from the Department of Agriculture, Land Reform and Rural Development, South Africa (DALRRD). A certificate of compliance in accordance with Article 17, paragraph 2, of the Nagoya Protocol has been issued by the Access and Benefit-Sharing Clearing-House (ABSCH) of the Department of Forestry, Fisheries and the Environment, South Africa (ABSCH-IRCC-ZA-262932-1). We thank S. Mahlangu for assistance in sample collection and J. Gladitsch for contributing to the laboratory work. Illumina sequencing was conducted through Novogene Europe. This research was funded in whole, or in part, by the Austrian Science Fund (FWF) [Grant DOI: 10.55776/35059-B]. For the purpose of Open Access, the author has applied a CC by public copyright licence to any Author-Accepted Manuscript (AAM) version arising from this submission. The authors acknowledge the financial support from the University of Graz.

## References

Adams, C. I. M., Hoekstra, L. A., Muell, M. R., & Janzen, F. J. (2019). A Brief Review of Non-Avian Reptile Environmental DNA (eDNA), with a Case Study of Painted Turtle (Chrysemys picta) eDNA Under Field Conditions. Diversity, 11(4), 50. 10.3390/d11040050

Allen, M. C., Kwait, R., Vastano, A., Kisurin, A., Zoccolo, I., Jaffe, B. D., Angle, J. C., Maslo, B., & Lockwood, J. L. (2023). Sampling environmental DNA from trees and soil to detect cryptic arboreal mammals. Scientific Reports, 13(1), 1–13. 10.1038/s41598-023-27512-8

Alonso, R. S., McClintock, B. T., Lyren, L. M., Boydston, E. E., & Crooks, K. R. (2015). Mark-recapture and mark-resight methods for estimating abundance with remote cameras: A carnivore case study. PloS One, 10(3), e0123032. 10.1371/journal.pone.0123032

Andruszkiewicz Allan, E., Zhang, W. G., C. Lavery, A., & F. Govindarajan, A. (2021). Environmental DNA shedding and decay rates from diverse animal forms and thermal regimes. Environmental DNA, 3(2), 492–514. 10.1002/edn3.141

Bai, R., Ji, Y., Hu, W., Hu, Z., He, S., Onditi, K. O., Wang, H., Xiong, Y., Zhang, J., Jiang, X., & Li, X. (2025). Seasonal differences and downstream transport of terrestrial vertebrate eDNA: Insights from integrated camera-trap survey and water eDNA metabarcoding. Global Ecology and Conservation, 62, e03854. 10.1016/j.gecco.2025.e03854

Barnes, M. A., & Turner, C. R. (2016). The ecology of environmental DNA and implications for conservation genetics. Conservation Genetics, 17(1), 1–17. 10.1007/s10592-015-0775-4

Barnes, M. A., Turner, C. R., Jerde, C. L., Renshaw, M. A., Chadderton, W. L., & Lodge, D. M. (2014). Environmental Conditions Influence eDNA Persistence in Aquatic Systems. Environmental Science & Technology, 48(3), 1819–1827. 10.1021/es404734p

Bates, D., Mächler, M., Bolker, B., & Walker, S. (2015). Fitting Linear Mixed-Effects Models Using lme4. Journal of Statistical Software, 67(1). 10.18637/jss.v067.i01

Beery, S., Morris, D., Yang, S., Hernandez, A., Miao, Z., Vargas, L., Dodhia, R., Lavista, J., & Microsoft AI for Good Lab. (2026). MegaDetector (Version 0.1.0) [Python]. https://github.com/microsoft/MegaDetector (Original work published 2026)

Broadhurst, H. A., Gregory, L. M., Bleakley, E. K., Perkins, J. C., Lavin V, J., Bolton, P., Browett, S. S., Howe V, C., Singleton, N., Tansley, D., Sales, N. G., & McDevitt, A. D. (2021). Mapping differences in mammalian distributions and diversity using environmental DNA from rivers. SCIENCE OF THE TOTAL ENVIRONMENT, 801. 10.1016/j.scitotenv.2021.149724

Burian, A., Mauvisseau, Q., Bulling, M., Domisch, S., Qian, S., & Sweet, M. (n.d.). Improving the reliability of eDNA data interpretation. Retrieved April 12, 2026, from https://onlinelibrary.wiley.com/doi/10.1111/1755-0998.13367

Burton, A. C., Neilson, E., Moreira, D., Ladle, A., Steenweg, R., Fisher, J. T., Bayne, E., & Boutin, S. (2015). REVIEW: Wildlife camera trapping: A review and recommendations for linking surveys to ecological processes. Journal of Applied Ecology, 52(3), 675–685. 10.1111/1365-2664.12432

Buxton, A. S., Groombridge, J. J., & Griffiths, R. A. (2017). Is the detection of aquatic environmental DNA influenced by substrate type? PLOS ONE, 12(8). 10.1371/journal.pone.0183371

Caravaggi, A., Burton, A. C., Clark, D. A., Fisher, J. T., Grass, A., Green, S., Hobaiter, C., Hofmeester, T. R., Kalan, A. K., Rabaiotti, D., & Rivet, D. (2020). A review of factors to consider when using camera traps to study animal behavior to inform wildlife ecology and conservation. Conservation Science and Practice, 2(8), e239. 10.1111/csp2.239

Deiner, K., Bik, H. M., Mächler, E., Seymour, M., Lacoursière-Roussel, A., Altermatt, F., Creer, S., Bista, I., Lodge, D. M., De Vere, N., Pfrender, M. E., & Bernatchez, L. (2017). Environmental DNA metabarcoding: Transforming how we survey animal and plant communities. Molecular Ecology, 26(21), 5872–5895. 10.1111/mec.14350

GBIF.org. (2025). GBIF Home Page.

Greenberg, S., Godin, T., & Whittington, J. (2019). Design patterns for wildlife-related camera trap image analysis. Ecology and Evolution, 9(24), 13706–13730. 10.1002/ece3.5767

Grolemund, G., & Wickham, H. (2011). Dates and Times Made Easy with {lubridate}. Journal of Statistical Software, 40(3), 1–25.

Harper, L. R., Lawson Handley, L., Carpenter, A. I., Ghazali, M., Di Muri, C., Macgregor, C. J., Logan, T. W., Law, A., Breithaupt, T., Read, D. S., McDevitt, A. D., & Hänfling, B. (2019). Environmental DNA (eDNA) metabarcoding of pond water as a tool to survey conservation and management priority mammals. Biological Conservation, 238, 108225. 10.1016/j.biocon.2019.108225

Hartig, F. (2024). DHARMa: Residual Diagnostics for Hierarchical (Multi-Level / Mixed) Regression Models.

Iacaruso, N. J., Reves, O. P., Merkelz, S. J., Waldrep, C. L., & Davis, M. A. (2026). A systematic review evaluating the performance of eDNA methods relative to conventional methods for biodiversity monitoring. Ecography, 2026(5), e07952. 10.1002/ecog.07952

Ionescu, D., Bizic, M., Karnatak, R., Musseau, C. L., Onandia, G., Kasada, M., Berger, S. A., Nejstgaard, J. C., Ryo, M., Lischeid, G., Gessner, M. O., Wollrab, S., & Grossart, H.-P. (2022). From microbes to mammals: Pond biodiversity homogenization across different land-use types in an agricultural landscape. Ecological Monographs. 10.1002/ecm.1523

IPBES. (2019). Global assessment report on biodiversity and ecosystem services of the Intergovernmental Science-Policy Platform on Biodiversity and Ecosystem Services. 10.5281/ZENODO.6417333

Ishige, T., Miya, M., Ushio, M., Sado, T., Ushioda, M., Maebashi, K., Yonechi, R., Lagan, P., & Matsubayashi, H. (2017). Tropical-forest mammals as detected by environmental DNA at natural saltlicks in Borneo. Biological Conservation, 210, 281–285. 10.1016/j.biocon.2017.04.023

Jones, K. E., Bielby, J., Cardillo, M., Fritz, S. A., O’Dell, J., Orme, C. D. L., Safi, K., Sechrest, W., Boakes, E. H., Carbone, C., Connolly, C., Cutts, M. J., Foster, J. K., Grenyer, R., Habib, M., Plaster, C. A., Price, S. A., Rigby, E. A., Rist, J., … Purvis, A. (2009). PanTHERIA: a species-level database of life history, ecology, and geography of extant and recently extinct mammals. Ecology, 90(9), 2648. 10.1890/08-1494.1

Joseph, C., Faiq, M. E., Li, Z., & Chen, G. (2022). Persistence and degradation dynamics of eDNA affected by environmental factors in aquatic ecosystems. Hydrobiologia, 849(19), 4119–4133. 10.1007/s10750-022-04959-w

Kalan, A. K., Hohmann, G., Arandjelovic, M., Boesch, C., McCarthy, M. S., Agbor, A., Angedakin, S., Bailey, E., Balongelwa, C. W., Bessone, M., Bocksberger, G., Coxe, S. J., Deschner, T., Després-Einspenner, M.-L., Dieguez, P., Fruth, B., Herbinger, I., Granjon, A.-C., Head, J., … Kühl, H. S. (2019). Novelty Response of Wild African Apes to Camera Traps. Current Biology, 29(7), 1211–1217.e3. 10.1016/j.cub.2019.02.024

Kingdon, J. (2015). The Kingdon Field Guide to African Mammals (2nd editio). Bloomsbury.

Larsson, J. (2024). eulerr: Area-Proportional Euler and Venn Diagrams with Ellipses.

Leempoel, K., Hebert, T., & Hadly, E. A. (2020). A comparison of eDNA to camera trapping for assessment of terrestrial mammal diversity. Proceedings of the Royal Society B: Biological Sciences, 287(1918), 20192353. 10.1098/rspb.2019.2353

Lele, S. R., Keim, J. L., & Solymos, P. (2023). ResourceSelection: Resource Selection (Probability) Functions for Use-Availability Data.

Leray, M., Knowlton, N., & Machida, R. J. (2022). MIDORI2: A collection of quality controlled, preformatted, and regularly updated reference databases for taxonomic assignment of eukaryotic mitochondrial sequences. Environmental DNA, 4(4), 894–907. 10.1002/edn3.303

Longmire, J. L., Maltbie, M., Baker, R. J., & Texas Tech University. (1997). Use of “Lysis Buffer” in DNA isolation and its implication for museum collections. Museum of Texas Tech University. 10.5962/bhl.title.143318

Lyet, A., Pellissier, L., Valentini, A., Dejean, T., Hehmeyer, A., & Naidoo, R. (2021). eDNA sampled from stream networks correlates with camera trap detection rates of terrestrial mammals. Scientific Reports, 11(1), 11362. 10.1038/s41598-021-90598-5

Lynggaard, C., Bertelsen, M. F., Jensen, C. V., Johnson, M. S., Frøslev, T. G., Olsen, M. T., & Bohmann, K. (2022). Airborne environmental DNA for terrestrial vertebrate community monitoring. Current Biology, 32(3), 701–707.e5. 10.1016/j.cub.2021.12.014

Martin, M. (2011). CUTADAPT removes adapter sequences from high-throughput sequencing reads. EMBnet.Journal, 17. 10.14806/ej.17.1.200

Mas-Carrió, E., Schneider, J., Nasanbat, B., Ravchig, S., Buxton, M., Nyamukondiwa, C., Stoffel, C., Augugliaro, C., Ceacero, F., Taberlet, P., Glaizot, O., Christe, P., & Fumagalli, L. (2022). Assessing environmental DNA metabarcoding and camera trap surveys as complementary tools for biomonitoring of remote desert water bodies. Environmental DNA, 4(3), 580–595. 10.1002/edn3.274

McCartin, L. J., Vohsen, S. A., Ambrose, S. W., Layden, M., McFadden, C. S., Cordes, E. E., McDermott, J. M., & Herrera, S. (2022). Temperature Controls eDNA Persistence across Physicochemical Conditions in Seawater. Environmental Science & Technology, 56(12), 8629–8639. 10.1021/acs.est.2c01672

Meek, P. D., Ballard, G. A., Fleming, P. J. S., Schaefer, M., Williams, W., & Falzon, G. (2014). Camera traps can be heard and seen by animals. PLoS ONE, 9(10). 10.1371/journal.pone.0110832

Morris, J. W. (2022). Botsalano Game Reserve—A Visitor Guide (p. 36).

Neave, G., Murphy, B. P., Rangers, T., & Davies, H. F. (2025). Exotic Megaherbivores as Ecosystem Engineers in Australian Savannas: Do They Facilitate Predator Movement? Ecology and Evolution, 15(7), e71622. 10.1002/ece3.71622

Newton, J. P., Allentoft, M. E., Bateman, P. W., van der Heyde, M., & Nevill, P. (2025). Targeting Terrestrial Vertebrates With eDNA: Trends, Perspectives, and Considerations for Sampling. Environmental DNA, 7(1). 10.1002/edn3.70056

Newton, J. P., Nevill, P., Bateman, P. W., Campbell, M. A., & Allentoft, M. E. (2024). Spider webs capture environmental DNA from terrestrial vertebrates. iScience, 27(2). 10.1016/j.isci.2024.108904

O’Brien, T. G., Kinnaird, M. F., & Wibisono, H. T. (2003). Crouching tigers, hidden prey: Sumatran tiger and prey populations in a tropical forest landscape. Animal Conservation, 6(2), 131–139. 10.1017/S1367943003003172

Osner, N. (2026). TrapTagger [Python]. https://github.com/WildEyeConservation/TrapTagger (Original work published 2022)

Parr, C. S., Wilson, N., Leary, P., Schulz, K. S., Lans, K., Walley, L., Hammock, J. A., Goddard, A., Rice, J., Studer, M., Holmes, J. T. G., & Corrigan Robert J., Jr. (2014). The Encyclopedia of Life v2: Providing Global Access to Knowledge About Life on Earth. Biodiversity Data Journal, 29, e1079. 10.3897/BDJ.2.e1079

Posit team. (2024). RStudio: Integrated Development Environment for R.

R Core Team. (2023). R: A language and environment for statistical computing. R Foundation for Statistical Computing.

Riaz, T., Shehzad, W., Viari, A., Pompanon, F., Taberlet, P., & Coissac, E. (2011). EcoPrimers: Inference of new DNA barcode markers from whole genome sequence analysis. Nucleic Acids Research, 39(21), 1–11. 10.1093/nar/gkr732

Robin, X., Turck, N., Hainard, A., Tiberti, N., Lisacek, F., Sanchez, J.-C., & Müller, M. (2011). pROC: an open-source package for R and S+ to analyze and compare ROC curves. BMC Bioinformatics, 12, 77.

Rodgers, T. W., & Mock, K. E. (2015). Drinking water as a source of environmental DNA for the detection of terrestrial wildlife species. Conservation Genetics Resources, 7(3), 693–696. 10.1007/s12686-015-0478-7

Rognes, T., Flouri, T., Nichols, B., Quince, C., & Mahé, F. (2016). VSEARCH: a versatile open source tool for metagenomics. PeerJ, 4, e2584.

Sales, N. G., McKenzie, M. B., Drake, J., Harper, L. R., Browett, S. S., Coscia, I., Wangensteen, O. S., Baillie, C., Bryce, E., Dawson, D. A., Ochu, E., Haenfling, B., Handley, L. L., Mariani, S., Lambin, X., Sutherland, C., & McDevitt, A. D. (2020). Fishing for mammals: Landscape-level monitoring of terrestrial and semi-aquatic communities using eDNA from riverine systems. JOURNAL OF APPLIED ECOLOGY, 57(4), 707–716. 10.1111/1365-2664.13592

Sauseng, G., & Schenekar, T. (2025). Water Interaction Type Affects Environmental DNA Shedding Rates of Terrestrial Mammal eDNA Into Surface Water Bodies. Environmental DNA, 7. 10.1002/edn3.70120

Schenekar, T., Baxter, J., Metlholo, Phukuntsi, M., Sedlmayr, I., Weckworth, B., & Mwale, M. (2024). Optimizing waterborne eDNA capture from waterholes in savanna systems under remote field conditions. Molecular Ecology Resources, 00, e13942. 10.1111/1755-0998.13942

Schenekar, T., Baxter, J., Sedlmayr, I., Gladitsch, J., Mahlangu, S., & Mwale, M. (2026). Unveiling Vertebrate Biodiversity in Arid and Semi-Arid Terrestrial Ecosystems Through eDNA Metabarcoding at Savanna Waterholes. Evolutionary Applications, 19(2), e70200. 10.1111/eva.70200

Sedlmayr, I., & Schenekar, T. (2024). eDNA state and medium affect DNA degradation patterns in semi-natural systems of Southern African waterholes. Environmental DNA, 1–14. 10.1002/edn3.70025

Sherub, K., Thurnheer, S., Lüthi, M., Marques, V., Lyet, A., Dhendup, T., Dorji, L., Albouy, C., & Pellissier, L. (2026). Comparing Watershed-Based eDNA Sampling and Camera Trapping for Assessing Mammal Diversity in North-Western Bhutan. Environmental DNA. 10.1002/edn3.70243

Strickler, K. M., Fremier, A. K., & Goldberg, C. S. (2015). Quantifying effects of UV-B, temperature, and pH on eDNA degradation in aquatic microcosms. Biological Conservation, 183, 85–92. 10.1016/j.biocon.2014.11.038

Taberlet, P., Coissac, E., Hajibabaei, M., & Rieseberg, L. H. (2012). Environmental DNA. Molecular Ecology, 21, 1789–1793. 10.1111/j.1365-294X.2012.05542.x

Tanwar, K. S., Sadhu, A., & Jhala, Y. V. (2021). Camera trap placement for evaluating species richness, abundance, and activity. Scientific Reports, 11(1), 23050. 10.1038/s41598-021-02459-w

Thomsen, P. F., & Willerslev, E. (2015). Environmental DNA – An emerging tool in conservation for monitoring past and present biodiversity. Biological Conservation, 183, 4–18. 10.1016/j.biocon.2014.11.019

Tsuji, S., Ushio, M., Sakurai, S., Minamoto, T., & Yamanaka, H. (2017). Water temperature-dependent degradation of environmental DNA and its relation to bacterial abundance. PLOS ONE, 12(4), e0176608. 10.1371/journal.pone.0176608

Turner, C. R., Uy, K. L., & Everhart, R. C. (2015). Fish environmental DNA is more concentrated in aquatic sediments than surface water. Biological Conservation, 183, 93–102. 10.1016/j.biocon.2014.11.017

Ushio, M., Fukuda, H., Inoue, T., Makoto, K., Kishida, O., Sato, K., Murata, K., Nikaido, M., Sado, T., Sato, Y., Takeshita, M., Iwasaki, W., Yamanaka, H., Kondoh, M., & Miya, M. (2017). Environmental DNA enables detection of terrestrial mammals from forest pond water. Molecular Ecology Resources, 17(6). 10.1111/1755-0998.12690

Vélez, J., McShea, W., Shamon, H., Castiblanco-Camacho, P. J., Tabak, M. A., Chalmers, C., Fergus, P., & Fieberg, J. (2023). An evaluation of platforms for processing camera-trap data using artificial intelligence. Methods in Ecology and Evolution, 14(2), 459–477. 10.1111/2041-210X.14044

Wickham, H. (2016). ggplot2: Elegant Graphics for Data Analysis. Springer-Verlag.

Wickham, H. (2023). stringr: Simple, Consistent Wrappers for Common String Operations.

Wickham, H., François, R., Henry, L., Müller, K., & Vaughan, D. (2023). dplyr: A Grammar of Data Manipulation (1.1.4).

Wickham, H., & Henry, L. (2023). purrr: Functional Programming Tools.

Wickham, H., Hester, J., & Bryan, J. (2024). readr: Read Rectangular Text Data.

Wickham, H., Vaughan, D., & Girlich, M. (2024). tidyr: Tidy Messy Data.

Williams, K. E., Huyvaert, K. P., Vercauteren, K. C., Davis, A. J., & Piaggio, A. J. (2018). Detection and persistence of environmental DNA from an invasive, terrestrial mammal. Ecology and Evolution, 8(1), 688–695. 10.1002/ece3.3698

Xiao, Z., Dong, S., Zhang, Z., Qi, S., Wan, Y., & Song, Z. (2025). Spatio-temporal distribution of environmental DNA from amphibian and turtle species in a pond ecosystem. Environmental Research, 279, 121834. 10.1016/j.envres.2025.121834

Yates, M. C., Wilcox, T. M., Kay, S., Peres-Neto, P., & Heath, D. D. (2025). A Framework to Unify the Relationship Between Numerical Abundance, Biomass, and Environmental DNA. Environmental DNA, 7(2), e70073. 10.1002/edn3.70073

Zimmermann, F., & Rovero, F. (2016). Field deployment of camera traps (pp. 22–32). 10.2307/jj.28833751.11

