## Supporting Information 1 for "Species ecology, behavior, and substrate type shape eDNA detection of mammals at savanna waterholes"

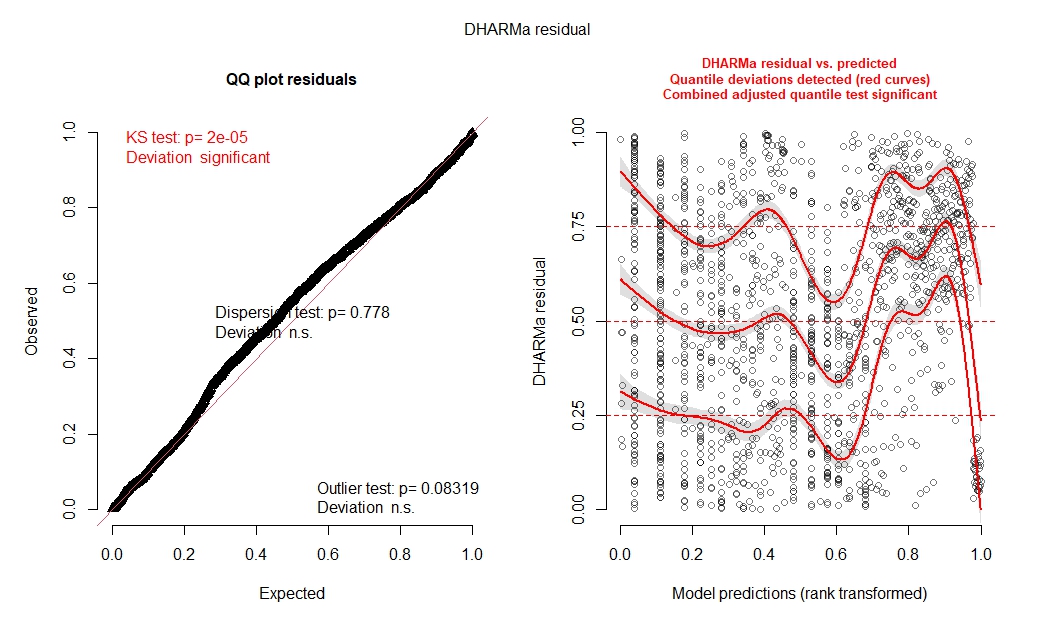

**Figure S1**. DHARMa residual diagnostics for the mixed-effects model of species traits, activity metrics and environmental variables on eDNA detection probability. DHARMa-scaled residuals for the binomial generalized linear mixed-effects model for modelling of species traits, activity metrics and environmental variables on eDNA detection probability. Left: quantile–quantile (QQ) plot of simulated versus expected residuals, including Kolmogorov–Smirnov, dispersion and outlier tests. Right: residuals plotted against model predictions (rank-transformed), with smoothed quantile curves (red) and the combined quantile test indicating significant deviations from perfect uniformity.

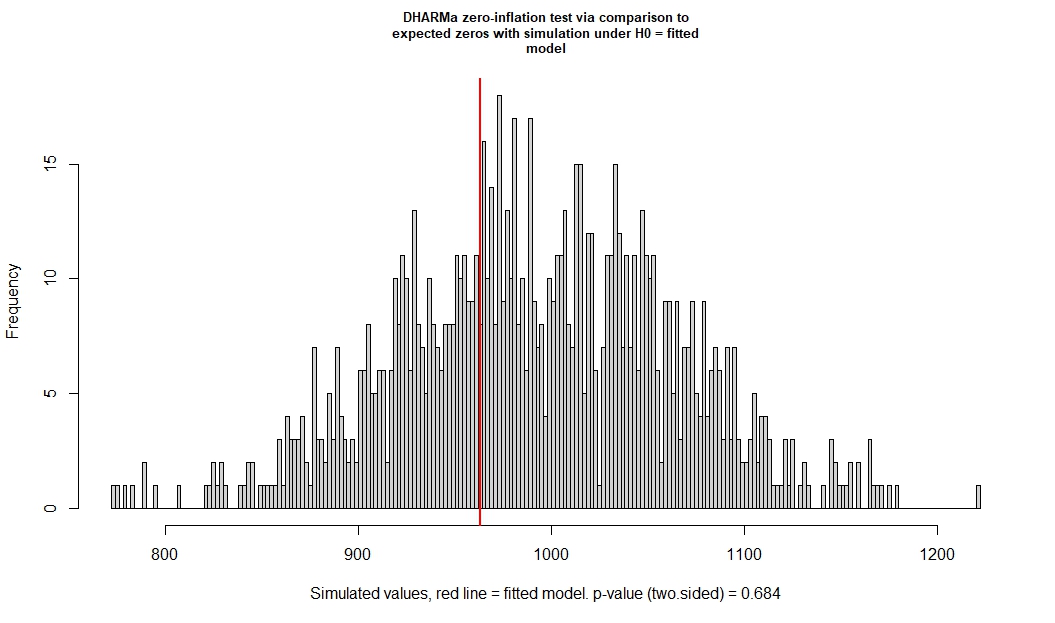
**Figure S2.** DHARMa zero-inflation test for the mixed-effects model of species traits, activity metrics and environmental variables on eDNA detection probability. Histogram of the number of zeros in simulated datasets generated from the fitted binomial generalized linear mixed-effects model (“Mixed-effects modelling of species traits, activity metrics and environmental variables on eDNA detection probability”). The red vertical line shows the observed number of zeros;
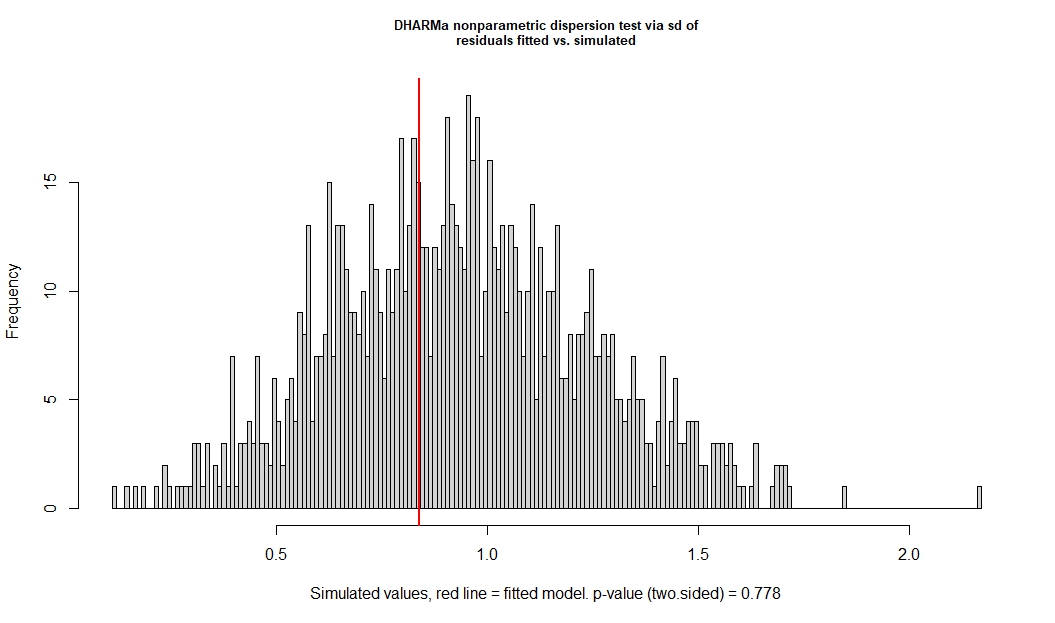
the associated two-sided DHARMa test indicates no evidence for zero inflation (p = 0.684).

**Figure S3**. DHARMa dispersion test for the mixed-effects model of species traits, activity metrics and environmental variables on eDNA detection probability. Histogram of the standard deviation of simulated residuals from the fitted binomial generalized linear mixed-effects model (“Mixed-effects modelling of species traits, activity metrics and environmental variables on eDNA detection probability”), used for the DHARMa nonparametric dispersion test. The red vertical line indicates the standard deviation of the observed residuals; the two-sided test shows no evidence of over- or underdispersion (p = 0.778).

**Table S1**. Fixed effects of the base mixed-effects model without spatial terms for eDNA detection probability at the sample level. Estimates are on the logit scale. Std. Error is the standard error of the estimated coefficient, z value is the Wald test statistic (estimate divided by its standard error), and P-value is the corresponding probability of obtaining a test statistic at least as extreme under the null hypothesis that the true coefficient is zero. *: p < 0.05, **: p < 0.01, *** p < 0.001

| **Predictor** | **Estimate** | **Std. Error** | **z value** | **P-value** |
| --- | --- | --- | --- | --- |
| (Intercept) | -3.15 | 0.49 | -6.47 | < 0.001*** |
| Adult body mass | 0.24 | 0.34 | 0.70 | 0.487 |
| Adult head body length | 0.31 | 0.46 | 0.67 | 0.503 |
| Social group size | 0.71 | 0.33 | 2.14 | 0.033 |
| Trophic level 2 (omnivore) | -1.86 | 0.73 | -2.55 | 0.011* |
| Trophic level 3 (carnivore) | -2.19 | 0.83 | -2.64 | 0.008** |
| Number of visitation events | 0.12 | 0.03 | 3.50 | < 0.001*** |
| Number of individual observations | 0.05 | 0.02 | 2.00 | 0.045* |
| Water interaction intensity | 0.14 | 0.05 | 2.98 | 0.003** |
| time since last visit - same period | -0.05 | 0.04 | -1.30 | 0.194 |
| time since last visit - overall | -0.28 | 0.09 | -3.17 | 0.002** |
| No previous visit - same period | 0.54 | 0.27 | 1.98 | 0.048* |
| No previous visit - overall | -1.11 | 0.30 | -3.67 | < 0.001*** |
| Season (wet) | 0.09 | 0.29 | 0.31 | 0.757 |
| Temperature | -0.08 | 0.12 | -0.65 | 0.516 |
| pH | -0.00 | 0.08 | -0.06 | 0.952 |
| Light intensity | 0.01 | 0.08 | 0.18 | 0.855 |
| Conductivity | -0.05 | 0.08 | -0.63 | 0.528 |
| Oxygen concentration | -0.01 | 0.06 | -0.21 | 0.835 |

**Table S2.** Summary of fixed effects of the binomial mixed‑effects model of eDNA detection probability also incorporating substrate type. Sediment was used as the reference level for the medium effect, therefore, main effects represent relationships in sediment samples, while interaction terms (water × predictor) indicate how these relationships changed in water samples relative to sediment. For each predictor, the estimate (on logit scale), the standard error of the estimated coefficient (Std. Error), Wald test statistic (z-value) and P-value are given. *: p < 0.05, **: p < 0.01, *** p < 0.001 A negative Estimate (β) indicates a decreased, a positive an increased probability of detection.

| **Predictor** | **Estimate (β)** | **Std. Error** | **z** | **p-value** |  |
| --- | --- | --- | --- | --- | --- |
| Intercept | -2.540 | 0.512 | -4.961 | <0.001 | *** |
| medium (water) | -1.558 | 0.151 | -10.336 | <0.001 | *** |
| Adult Body Mass | 0.258 | 0.353 | 0.730 | 0.465 |  |
| Adult Head-Body Length | 0.165 | 0.482 | 0.342 | 0.732 |  |
| Social Group Size | 0.931 | 0.341 | 2.728 | 0.006 | ** |
| Trophic Level 2 (Omnivore) | -2.575 | 0.762 | -3.379 | <0.001 | *** |
| Trophic Level 3 (Cernivore) | -3.615 | 0.937 | -3.856 | <0.001 | *** |
| individual observations | 0.008 | 0.032 | 0.244 | 0.807 |  |
| number of visits | 0.109 | 0.044 | 2.451 | 0.014 | * |
| time passed (same sampling period) | -0.044 | 0.049 | -0.907 | 0.364 |  |
| time passed (overall) | -0.347 | 0.121 | -2.878 | 0.004 | ** |
| No. of previous visits (same sampling period) | 0.597 | 0.372 | 1.606 | 0.108 |  |
| No. of previous visits overall | -1.485 | 0.397 | -3.742 | <0.001 | *** |
| behaviour percentile | 0.060 | 0.059 | 1.012 | 0.312 |  |
| Temperature (mean) | -0.186 | 0.133 | -1.395 | 0.163 |  |
| pH | 0.084 | 0.085 | 0.990 | 0.322 |  |
| lux | 0.220 | 0.096 | 2.299 | 0.022 | * |
| conductivity | 0.077 | 0.087 | 0.886 | 0.375 |  |
| O2 saturation | -0.103 | 0.072 | -1.434 | 0.152 |  |
| season (wet) | 0.082 | 0.307 | 0.267 | 0.790 |  |
| water × Adult Body Mass | -0.019 | 0.049 | -0.396 | 0.692 |  |
| water × Adult Head-Body Length | 0.328 | 0.106 | 3.094 | 0.002 | ** |
| water × Social Group Size | -0.460 | 0.056 | -8.207 | <0.001 | *** |
| water × Trophic Level 2 (omnivore) | 1.506 | 0.217 | 6.927 | <0.001 | *** |
| water × Trophic Level 3 (carnivore) | 2.477 | 0.498 | 4.971 | <0.001 | *** |
| water × individual observations | 0.088 | 0.042 | 2.082 | 0.037 | * |
| water × No. visitation events | 0.029 | 0.056 | 0.517 | 0.605 |  |
| water × time passed (same period) | -0.026 | 0.073 | -0.360 | 0.719 |  |
| water × time passed (overall) | 0.167 | 0.174 | 0.957 | 0.338 |  |
| water × No. previous visits (same sampling period) | -0.173 | 0.564 | -0.306 | 0.760 |  |
| water × No. previous visits overall | 0.967 | 0.582 | 1.660 | 0.097 | . |
| water × behaviour percentile | 0.194 | 0.081 | 2.407 | 0.016 | * |
| water × Temperature | 0.245 | 0.087 | 2.808 | 0.005 | ** |
| water × pH | -0.241 | 0.068 | -3.552 | <0.001 | *** |
| water × lux | -0.519 | 0.088 | -5.895 | <0.001 | *** |
| water × conductivity | -0.310 | 0.060 | -5.171 | <0.001 | *** |
| water × O2 saturation | 0.207 | 0.065 | 3.168 | 0.002 | ** |

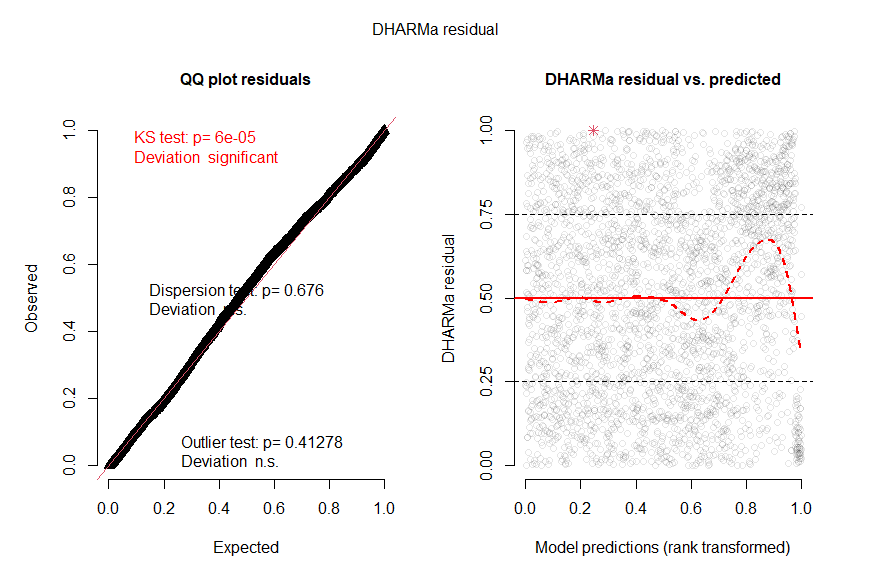

**Figure S4.** Residuals for the binomial generalized linear mixed-effects model for modelling of species traits, activity metrics, substrate and environmental variables on eDNA detection probability. Left: quantile–quantile (QQ) plot of simulated versus expected residuals, including Kolmogorov–Smirnov, dispersion and outlier tests. Right: residuals plotted against model predictions (rank-transformed).

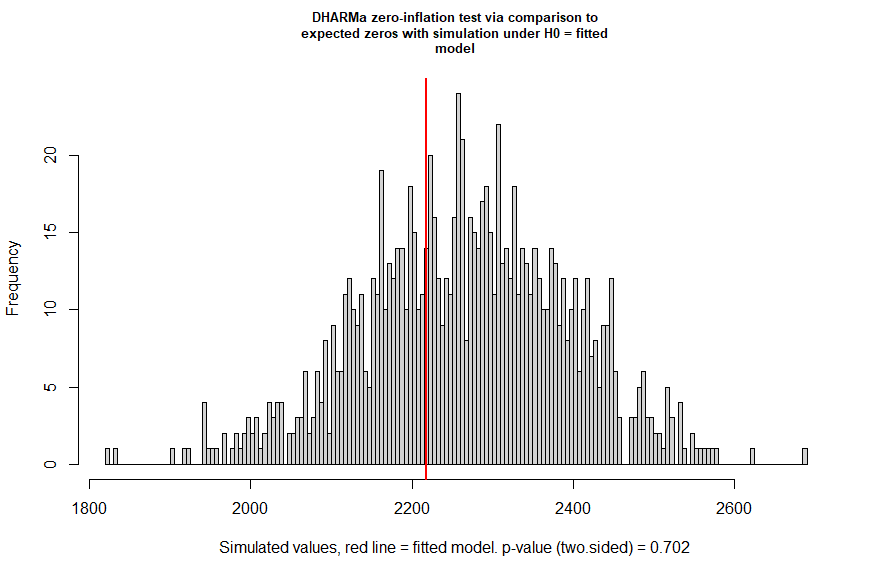

**Figure S5.** DHARMa zero-inflation test for the mixed-effects model incorporating substrate. Histogram of the number of zeros in simulated datasets generated from the fitted binomial generalized linear mixed-effects model (“Mixed-effects modelling of species traits, activity metrics, substrate and environmental variables on eDNA detection probability”). The red vertical line shows the observed number of zeros; the associated two-sided DHARMa test indicates no evidence for zero inflation (p = 0.702).

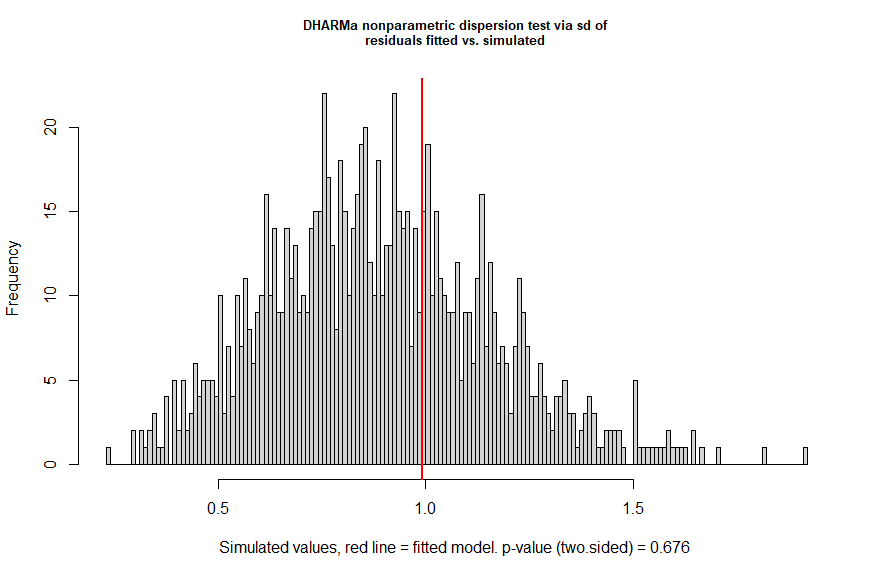

**Figure S6.** DHARMa dispersion test for the mixed-effects model incorporating substrate. Histogram of the standard deviation of simulated residuals from the fitted binomial generalized linear mixed-effects model (“Mixed-effects modelling of species traits, activity metrics, substrate and environmental variables on eDNA detection probability”), used for the DHARMa nonparametric dispersion test. The red vertical line indicates the standard deviation of the observed residuals; the two-sided test shows no evidence of over- or underdispersion (p = 0.676).

**Table S3.** Fixed effects of the spatial mixed-effects model including spatial terms for eDNA detection probability at the sample level. Estimates are on the logit scale. Std. Error is the standard error of the estimated coefficient, z value is the Wald test statistic (estimate divided by its standard error), and P-value is the corresponding probability of obtaining a test statistic at least as extreme under the null hypothesis that the true coefficient is zero. *: p < 0.05, **: p < 0.01, *** p < 0.001

| **Predictor** | **Estimate** | **Std. Error** | **z value** | **P-value** |
| --- | --- | --- | --- | --- |
| (Intercept) | -3.15 | 0.48 | -6.51 | < 0.001*** |
| Adult body mass | 0.24 | 0.34 | 0.69 | 0.487 |
| Adult head body length | 0.31 | 0.46 | 0.68 | 0.500 |
| Social group size | 0.71 | 0.33 | 2.15 | 0.032 |
| Trophic level 2 (omnivore) | -1.86 | 0.72 | -2.57 | 0.010* |
| Trophic level 3 (carnivore) | -2.19 | 0.82 | -2.65 | 0.008** |
| Number of visitation events | 0.11 | 0.03 | 3.38 | < 0.001*** |
| Number of individual observations | -0.03 | 0.03 | -0.94 | 0.347 |
| Water interaction intensity | 0.14 | 0.05 | 3.03 | 0.002** |
| time since last visit - same period | -0.05 | 0.04 | -1.35 | 0.179 |
| time since last visit - overall | -0.28 | 0.09 | -3.17 | 0.002** |
| No previous visit - same period | 0.53 | 0.27 | 1.95 | 0.052 |
| No previous visit - overall | -1.10 | 0.30 | -3.66 | < 0.001*** |
| Season (wet) | 0.10 | 0.29 | 0.34 | 0.737 |
| Temperature | -0.08 | 0.12 | -0.70 | 0.484 |
| pH | -0.00 | 0.08 | -0.06 | 0.956 |
| Light intensity | 0.02 | 0.08 | 0.20 | 0.843 |
| Conductivity | -0.06 | 0.08 | -0.71 | 0.476 |
| Oxygen concentration | -0.01 | 0.06 | -0.18 | 0.854 |
| Number of water interactions – local | 0.02 | 0.04 | 0.44 | 0.661 |
| Number of water interactions – same sampling spot | 0.10 | 0.03 | 3.48 | < 0.001*** |

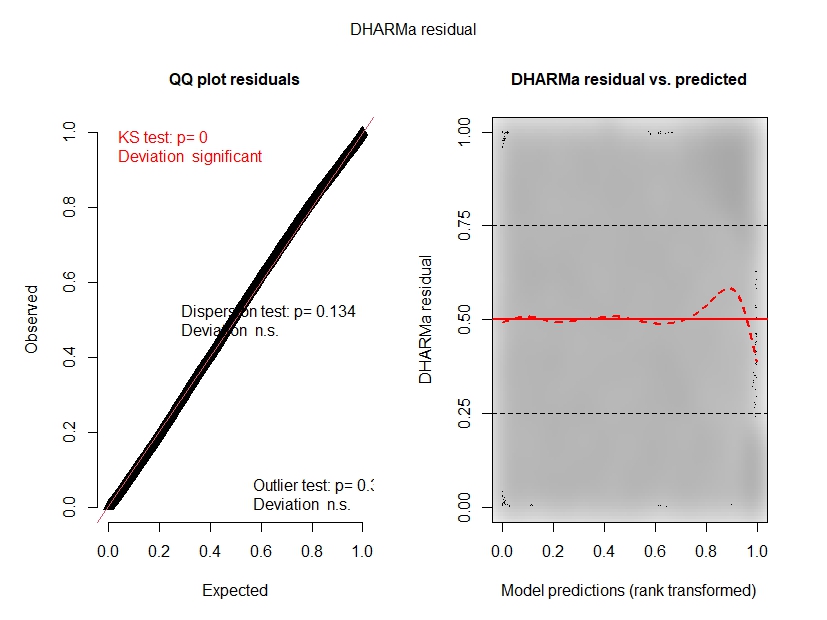

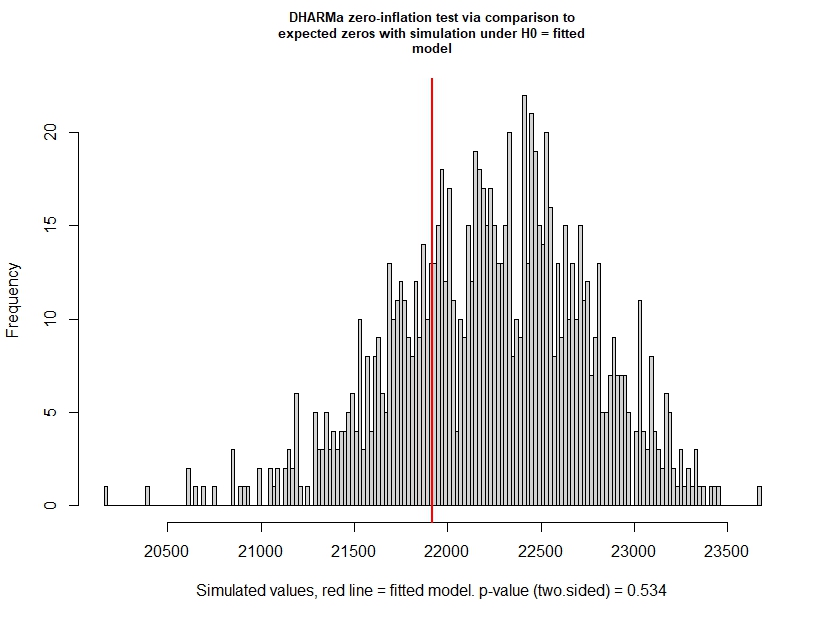
**Figure S7**.DHARMa residual diagnostics for the non-spatial mixed-effects model on detection at the sample level. DHARMa-scaled residuals for the binomial generalized linear mixed-effects model for modelling of species traits, activity metrics and environmental variables on eDNA detection probability. Left: quantile–quantile (QQ) plot of simulated versus expected residuals, including Kolmogorov–Smirnov, dispersion and outlier tests. Right: residuals plotted against model predictions (rank-transformed), with smoothed quantile curves (red) and the combined quantile test indicating significant deviations from perfect uniformity.

**Figure S8.** DHARMa zero-inflation test for the non-spatial mixed-effects model on detection at the sample level. Histogram of the number of zeros in simulated datasets generated from the fitted binomial generalized linear mixed-effects model (“Mixed-effects modelling of species traits, activity metrics and environmental variables on eDNA detection probability”). The red vertical line shows the observed number of zeros; the associated two-sided DHARMa test indicates no evidence for zero inflation (p = 0.534).

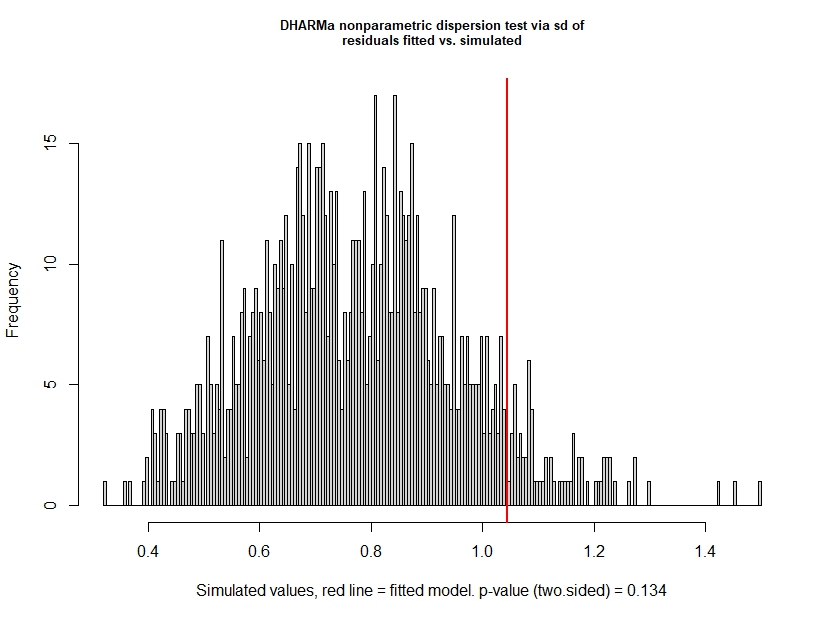

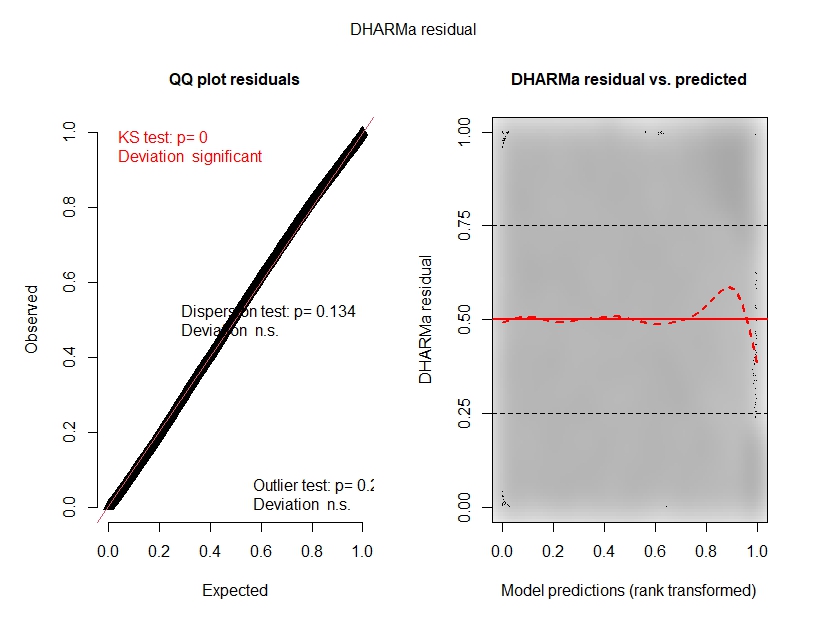
**Figure S9.** DHARMa dispersion for the non-spatial mixed-effects model on detection at the sample level. Histogram of the standard deviation of simulated residuals from the fitted binomial generalized linear mixed-effects model (“Mixed-effects modelling of species traits, activity metrics and environmental variables on eDNA detection probability”), used for the DHARMa nonparametric dispersion test. The red vertical line indicates the standard deviation of the observed residuals; the two-sided test shows no evidence of over- or underdispersion (p = 0.134).

**Figure S10**. DHARMa residual diagnostics for the spatial mixed-effects model on detection at the sample level. DHARMa-scaled residuals for the binomial generalized linear mixed-effects model for modelling of species traits, activity metrics, spatial information and environmental variables on eDNA detection probability. Left: quantile–quantile (QQ) plot of simulated versus expected residuals, including Kolmogorov–Smirnov, dispersion and outlier tests. Right: residuals plotted against model predictions (rank-transformed), with smoothed quantile curves (red).

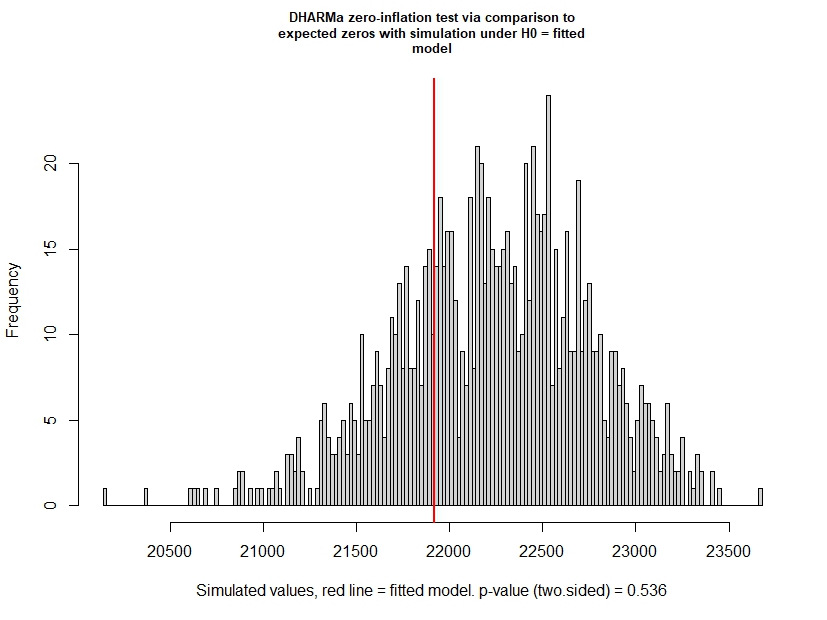

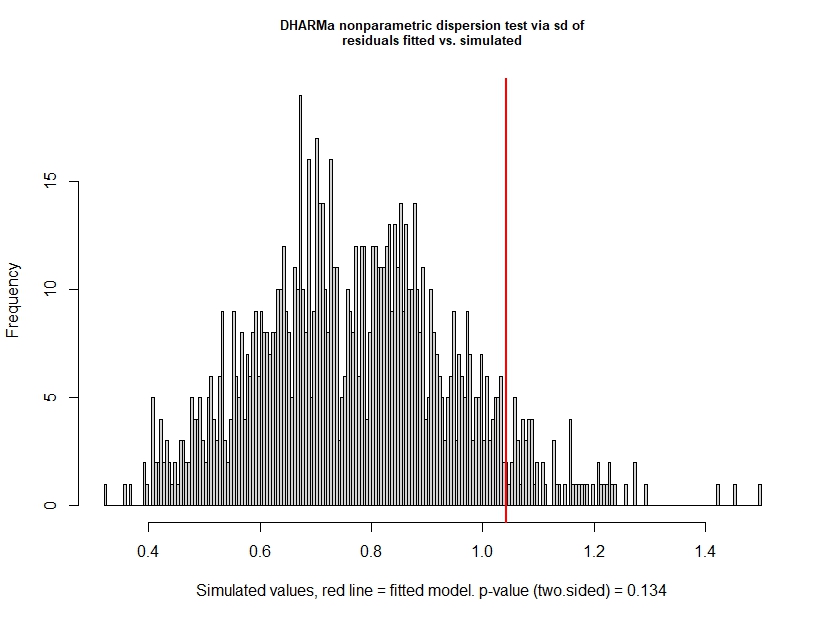
**Figure S11**. DHARMa zero-inflation test for the spatial mixed-effects model on detection at the sample level. Histogram of the number of zeros in simulated datasets generated from the fitted binomial generalized linear mixed-effects model (“Mixed-effects modelling of species traits, activity metrics, spatial information and environmental variables on eDNA detection probability”). The red vertical line shows the observed number of zeros; the associated two-sided DHARMa test indicates no evidence for zero inflation (p = 0.534).

**Figure S12**. DHARMa dispersion for the spatial mixed-effects model on detection at the sample level. Histogram of the standard deviation of simulated residuals from the fitted binomial generalized linear mixed-effects model (“Mixed-effects modelling of species traits, activity metrics, spatial information and environmental variables on eDNA detection probability”), used for the DHARMa nonparametric dispersion test. The red vertical line indicates the standard deviation of the observed residuals; the two-sided test shows no evidence of over- or underdispersion (p = 0.134).

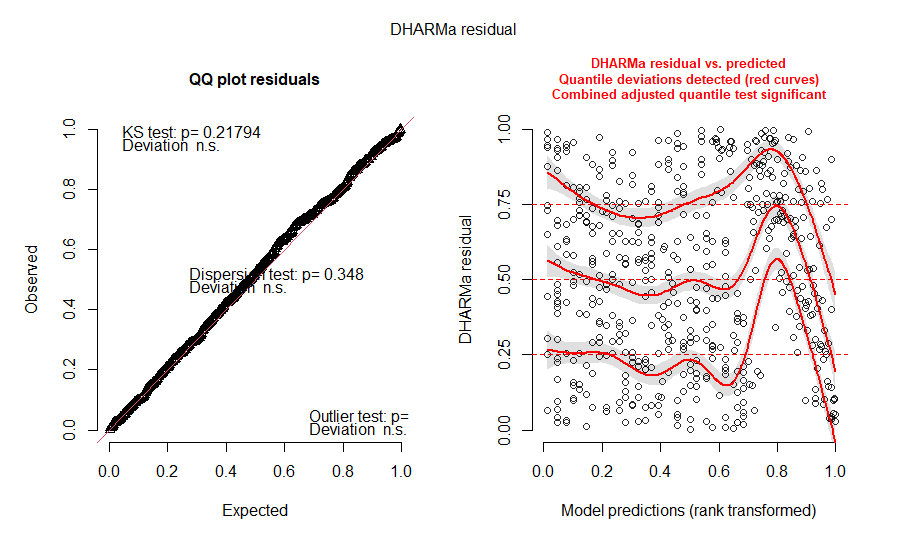

**Figure S13.** DHARMa residual diagnostics for the mixed-effects model for soil samples collected at wildlife trails. DHARMa-scaled residuals for the binomial generalized linear mixed-effects model for modelling of species traits and activity metrics on eDNA detection probability. Left: quantile–quantile (QQ) plot of simulated versus expected residuals, including Kolmogorov–Smirnov, dispersion and outlier tests. Right: residuals plotted against model predictions (rank-transformed), with smoothed quantile curves (red) and the combined quantile test indicating significant deviations from perfect uniformity.

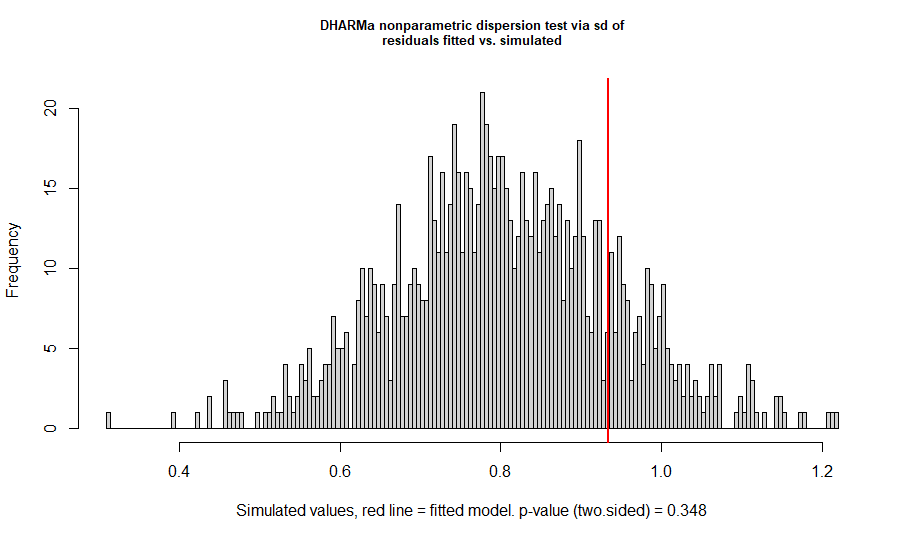

**Figure S14.** DHARMa zero-inflation test for the mixed-effects model for soil samples collected at wildlife trails. Histogram of the number of zeros in simulated datasets generated from the fitted binomial generalized linear mixed-effects model (“Mixed-effects modelling of species traits and activity metrics on eDNA detection probability”). The red vertical line shows the observed number of zeros; the associated two-sided DHARMa test indicates no evidence for zero inflation (p = 0.348).

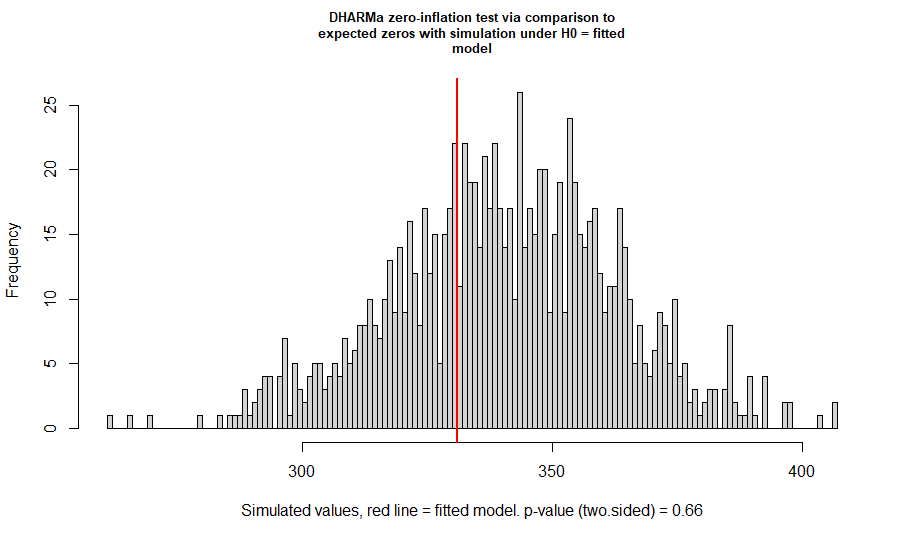

**Figure S15.** DHARMa dispersion test for the mixed-effects model for soil samples collected at wildlife trails. Histogram of the standard deviation of simulated residuals from the fitted binomial generalized linear mixed-effects model (“Mixed-effects modelling of species traits and activity metrics on eDNA detection probability”), used for the DHARMa nonparametric dispersion test. The red vertical line indicates the standard deviation of the observed residuals; the two-sided test shows no evidence of over- or underdispersion (p = 0.66).
