## Supporting Information 2 for "Species ecology, behavior, and substrate type shape eDNA detection of mammals at savanna waterholes": 02_multiple_sps.R

```
# Script to separate entries in which multiple sampling points were checked

setwd(dirname(rstudioapi::getActiveDocumentContext()$path))


library(lubridate)
library(dplyr)
library(readr)
library(tidyr)


# Import csv from  as a dataframe
raw_data <- read_csv("../01_raw_data/01_CT_data_all.csv")  %>% select(-`...1`)


# Split date and time and remove the seconds from DateTime column

raw_data <- raw_data %>%
  mutate(
    DateTime = as.POSIXct(DateTime, format = "%Y-%m-%d %H:%M:%S"),
    Date = as.Date(DateTime),                      
    Time = format(DateTime, format = "%H:%M")     
  )


# Compute the number of checked sampling points for each entry
raw_data <- raw_data %>%
  rowwise() %>%
  mutate(sp_count = sum(c_across(sp1:sp8))) %>%
  ungroup()

### Convert to long format. Needs a workaround for non-valid data because the Trails do not have sps and therefore "NA" values in sp_count
# Handle valid rows (where sp_count is not NA)
valid_data <- raw_data %>%
  filter(!is.na(sp_count)) %>%
  pivot_longer(cols = sp1:sp8, names_to = "sp", values_to = "Presence") %>%
  filter(Presence == TRUE) %>%
  mutate(Individuals_divided_by_sp = Individuals / sp_count) %>%
  select(-Presence)

# Handle NA rows (keep them with one entry and carry Individuals over)
na_data <- raw_data %>%
  filter(is.na(sp_count)) %>%
  mutate(
    sp = NA,
    Individuals_divided_by_sp = Individuals
  ) %>%
  select(-c(sp1, sp2, sp3, sp4, sp5, sp6, sp7, sp8))  # remove sp1:sp8 to match the long format

# Combine both
raw_data_sp_multiplied <- bind_rows(valid_data, na_data)


write.csv(raw_data_sp_multiplied, "02_raw_data_sp_multiplied.csv")
```
