## Supporting Information 2 for "Species ecology, behavior, and substrate type shape eDNA detection of mammals at savanna waterholes": 03_remove_duplicates.R

```
# Script to remove duplicates (same animal species at same sp covered by two cameras at same time) from dataset

setwd(dirname(rstudioapi::getActiveDocumentContext()$path))


library(readr)
library(stringr)
library(dplyr)

# Import csv from  TimeLapse into a dataframe
raw_data_sp_multiplied <- read_csv("../02_multiple_sps/02_raw_data_sp_multiplied.csv")  %>% select(-`...1`)


# Extract camera name from RelativePath and create a new 'camera' column

raw_data_sp_multiplied <- raw_data_sp_multiplied %>%
  mutate(camera = str_extract(str_replace(RelativePath, "Cam\\s", "Cam"), "Cam\\w{2,}"))

# Excluding trails data for this step as the data is different: there are no sps and there are multiple entries per minute due to motion trigger
# 1. All rows with Waterhole == "Trails" to the new dataframe  
trails_data <- raw_data_sp_multiplied %>%  
  filter(Waterhole == "Trails")  
# 2. Original df without those rows  
raw_data_sp_multiplied <- raw_data_sp_multiplied %>%  
  filter(Waterhole != "Trails")  


# Correct the values in "Trails" to the exact locations Trail1, Trail2 and Trail3 based on the camera
trails_data$Waterhole <- with(trails_data, ifelse(  
  Waterhole == "Trails" & camera == "CamA1", "Trail1",  
  ifelse(  
    Waterhole == "Trails" & camera == "CamA2", "Trail2",  
    ifelse(  
      Waterhole == "Trails" & camera == "CamA3", "Trail3",  
      Waterhole    
    )  
  )  
))  


# Make a new variable that counts the number of different activities for each case
raw_data_sp_multiplied <- raw_data_sp_multiplied %>%
  rowwise() %>%
  mutate(behaviour_count = sum(c_across(all_of(c("standingclose", "drinking", "feetinwater", "feetdrink"))))) %>%
  ungroup()


# Check for duplicates
duplicates <- raw_data_sp_multiplied %>%
  group_by(DateTime, Species, Waterhole, sp) %>%
  filter(n() > 1) %>%  # Keep only groups with more than one entry
  ungroup()

# Check for triplicates
triplicates <- raw_data_sp_multiplied %>%
  group_by(DateTime, Species, Waterhole, sp) %>%
  filter(n() > 2) %>%  # Keep only groups with more than 2 entries
  ungroup()


# Check for unique entries
unique_entries <- raw_data_sp_multiplied %>%
  group_by(DateTime, Species, Waterhole, sp) %>%
  filter(n() == 1) %>%  # Keep only unique entries
  ungroup()


# Check for duplicates based on DateTime, Species, Waterhole, and sp and keep only those with the higher number of Individuals
duplicates_01_highest_N <- raw_data_sp_multiplied %>%
  group_by(DateTime, Species, Waterhole, sp) %>%
  filter(n() > 1) %>%  # Keep only groups with more than one entry
  slice_max(Individuals_divided_by_sp, n = 1, with_ties = TRUE) %>%  # Keep only the entry with the highest Individuals
  ungroup()


duplicates_02_highest_N_behaviour <- duplicates_01_highest_N %>%
  group_by(DateTime, Species, Waterhole, sp) %>%
  slice_max(behaviour_count, n = 1, with_ties = TRUE) %>%  # Keep only the entry with the highest behaviour count
  ungroup()

duplicates_03_highest_N_behaviour_random <- duplicates_02_highest_N_behaviour %>%  
  group_by(DateTime, Species, Waterhole, sp) %>%  
  # Randomly select one row per group  
  slice_sample(n = 1) %>%  
  ungroup()    


# Combine filtered data with unique entries and trails data
CT_data_duplicates_removed <- bind_rows(duplicates_03_highest_N_behaviour_random, unique_entries, trails_data)

write.csv(CT_data_duplicates_removed, "03_CT_data_duplicates_removed.csv")
```
