## Supporting Information 2 for "Species ecology, behavior, and substrate type shape eDNA detection of mammals at savanna waterholes": 03b_behaviour_split.R

```
# Script to separate entries in which multiple behaviours were checked at the same sampling point

setwd(dirname(rstudioapi::getActiveDocumentContext()$path))


library(lubridate)
library(dplyr)
library(readr)
library(tidyr)


# Import csv from  TimeLapse into a dataframe
data <- read_csv("../03_remove_duplicates/03_CT_data_duplicates_removed.csv")

# Compute the count of TRUE values for each row
data <- data %>%
  rowwise() %>%
  mutate(behaviour_count = sum(c_across(all_of(c("standingclose", "drinking", "feetinwater", "feetdrink"))))) %>%
  ungroup()

# Convert wide format to long format using the new variables and divide Individuals
data_behaviour_multiplied <- data %>%
  pivot_longer(cols = c("standingclose", "drinking", "feetinwater", "feetdrink"), 
               names_to = "behaviour", values_to = "Presence") %>%
  filter(Presence == TRUE) %>%
  mutate(Individuals_divided_by_sp_and_behaviour = Individuals_divided_by_sp / behaviour_count) %>%
  select(-Presence)  # Keep behaviour_count in the final dataframe
write.csv(data_behaviour_multiplied , "data_behaviour_split.csv")
```
