## Supporting Information 2 for "Species ecology, behavior, and substrate type shape eDNA detection of mammals at savanna waterholes": 04_species_names.R

```
setwd(dirname(rstudioapi::getActiveDocumentContext()$path))

library(lubridate)
library(dplyr)
library(readr)
library(tidyr)

#upload last csv resulting from "remove duplicates
CT_data_duplicates_removed <- read_csv("../03b_multiple_behaviour/data_behaviour_split.csv") %>% select(-`...1`)


#Remove all "notsures" from the dataset
CT_data_notsures_removed <- CT_data_duplicates_removed %>%
  filter(notsure != TRUE)

# Remove all "Mammals"
CT_data_mammals_removed <- CT_data_duplicates_removed %>%
  filter(Species !="Mammal")

unique_species <- unique(CT_data_mammals_removed$Species)
print(unique_species)

write.csv(unique_species, "unique_species.csv")


# Import the species Latin name lookup CSV file to make all names homogenous across the entire dataset
species_latin_lookup <- read.csv2("04_Species_names_latin.csv")

# Merge the lookup table with CT_data_incl_latin
CT_data_incl_latin <- merge(
  CT_data_mammals_removed,
  species_latin_lookup,
  by = "Species",
  all.x = TRUE
)

#rename the column Species to "Old_name"
colnames(CT_data_incl_latin)[colnames(CT_data_incl_latin) == "Species"] <- "Old_name"

#rename the colums species_Latin to Species
colnames(CT_data_incl_latin)[colnames(CT_data_incl_latin) == "Species_Latin"] <- "Species"

write.csv(CT_data_incl_latin , "04_CT_data_incl_latin.csv")
```
