## Supporting Information 2 for "Species ecology, behavior, and substrate type shape eDNA detection of mammals at savanna waterholes": 05_adding_observation_periods_CT.R

```
# Sccript to add the observation periods to the camera trap data
# Setwd
setwd(dirname(rstudioapi::getActiveDocumentContext()$path))

# Load required packages
library(lubridate)
library(dplyr)
library(readr)
library(tidyr)

#Load the csv with the period assignment 
period_assignment <- read_csv2('period_assignment_CT.csv')  


#Load the csv
CT_data <- read_csv('../04_species_names/04_CT_data_incl_latin.csv')   %>% select(-`...1`)


#Check uniqueness of (location, date) in period_assignment  
dups <- period_assignment %>%  
  count(Waterhole, RelativePath) %>%  
  filter(n > 1)  

if (nrow(dups) > 0) {  
  warning("In 'period_assignment', (Waterhole, RelativePath) is not unique for the following combinations:\n",  
          paste0("  - ", dups$Water, " | ", dups$RelativePath, " (n = ", dups$n, ")", collapse = "\n"))  
}  

# Join period and regular_period to CT_data
CT_data_periods_added <- CT_data  %>%  
  left_join(  
    period_assignment %>%  
      select(Waterhole, RelativePath, period, regular_period),  
    by = c("Waterhole", "RelativePath")  
  )  

# Rename "Waterhole" to "location" to match with eDNA data
names(CT_data_periods_added)[names(CT_data_periods_added) == "Waterhole"] <- "location"  

# Write csv
write.csv(CT_data_periods_added, "CT_data_periods_added.csv")
```
