## Supporting Information 2 for "Species ecology, behavior, and substrate type shape eDNA detection of mammals at savanna waterholes": 06_adding_visitation_events.R

```
# Script to individual animal visitations into visitations events using 30 minute intervals

# Setwd

setwd(dirname(rstudioapi::getActiveDocumentContext()$path))
# Load required packages
library(lubridate)
library(dplyr)
library(readr)
library(tidyr)
library(stringr)  

#Load the csv
CT_data <- read_csv('../05 period assignment/CT_data_periods_added.csv')   %>% select(-`...1`)

# Define the interval threshold of 30 min
event_threshold <- dminutes(30)

# 
# For naming the visitation events - Helper function: create species_code  
# -----------------------------  
make_species_code <- function(species) {  
 
  
  # Handle NAs safely  
  species <- ifelse(is.na(species), "", species)  
  
  # Split into words  
  words_list <- str_split(species, "\\s+")  
  
  codes <- vapply(words_list, function(words) {  
    words <- words[words != ""]   # remove empty elements  
    
    if (length(words) >= 2) {  
            w1 <- words[1]  
      w2 <- words[2]  
      code <- paste0(  
        substr(w1, 1, 1),  
        substr(w2, 1, 2)  
      )  
    } else if (length(words) == 1) {  
      w1 <- words[1]  
      code <- substr(w1, 1, min(3, nchar(w1)))  
    } else {  
      code <- "UNK"  
    }  

    
    code  
  }, FUN.VALUE = character(1))  
  
  codes  
}  


# Main processing  

CT_data_vis <- CT_data %>%  
  mutate(  
    DateTime = as.POSIXct(DateTime, tz = "UTC")    
  ) %>%  
  arrange(location, Species, DateTime) %>%  
  group_by(location, Species) %>%  
  mutate(  
    # Time difference to previous record within same location & Species  
    time_diff = DateTime - lag(DateTime),  
    
    # Start a new visit if:  
    # - this is the first record for that location & species (is.na(time_diff))  
    # - or the gap is > 30 minutes  
    new_event_flag = if_else(is.na(time_diff) | time_diff > event_threshold, 1L, 0L),  
    
    # Running event number per location & Species  
    event_number = cumsum(new_event_flag)  
  ) %>%  
  ungroup() %>%  
  # Create season code: "w" for wet, "d" for dry, "u" otherwise  
  mutate(  
    season_code = case_when(  
      Season == "wet" ~ "w",  
      Season == "dry" ~ "d",  
      TRUE            ~ "u"   # unknown / other  
    ),  
    # Create species code  
    species_code = make_species_code(Species),  
    # Build the visitation event ID:  
    # location_seasonCode_speciesCode_eventNumber  
    vis_event = paste(location, season_code, species_code, event_number, sep = "_")  
  )  

# Delete all unnecessary columns in final dataset
CT_data_vis$time_diff<- NULL
CT_data_vis$new_event_flag<- NULL
CT_data_vis$event_number<- NULL
CT_data_vis$season_code<- NULL
CT_data_vis$species_code<- NULL


# Write csv
write.csv(CT_data_vis , "CT_data_visitation_events_added.csv")
```
