## Supporting Information 2 for "Species ecology, behavior, and substrate type shape eDNA detection of mammals at savanna waterholes": 07_Venn_diagram.R

```
# Script to calculate some basic statistics and to produce the Venn (Euler) diagram
# Set wd
setwd(dirname(rstudioapi::getActiveDocumentContext()$path))


# Load the required packages

library(dplyr)
library(readr)
library(tidyr)
library(eulerr)  


#load CT data
CT_data <- read_csv("../06 adding visitation events/CT_data_visitation_events_added.csv")  %>% select(-`...1`)

# Load eDNA data per sample
eDNA_data_per_sample <- read_csv("eDNA_data_samples_incl_latin.csv")   %>% select(-`...1`)
# Load eDNA data per sampling evevnt
eDNA_data_per_sampling_event <- read_csv("eDNA_data_sampling_events_incl_latin.csv")   %>% select(-`...1`)


# Extract unique species from each dataset. For eDNA data, either dataset can be used
CT_species <- unique(CT_data$Species)
eDNA_species <- unique(eDNA_data_per_sampling_event$Species)

# Get species lists
species_list <- list(  
  CT   = CT_species,  
  eDNA = eDNA_species  
)  


# Creating a presence-absence matrix for all species
all_species <- sort(unique(c(CT_species, eDNA_species)))  

species_presence <- data.frame(  
  Species     = all_species,  
  CT_species  = ifelse(all_species %in% CT_species, 1, 0),  
  eDNA_species = ifelse(all_species %in% eDNA_species, 1, 0)  
)

# Write out presence absence matrix
write.csv(species_presence, "species_presence_CT_eDNA.csv", row.names = FALSE)  

# Quick check of set sizes  
length(CT_species)      # number of CT species  
length(eDNA_species)    # number of eDNA species  
length(intersect(CT_species, eDNA_species))  # number of shared species 

# Species lists of sets
only_CT   <- setdiff(CT_species, eDNA_species)  
only_eDNA <- setdiff(eDNA_species, CT_species)  
both      <- intersect(CT_species, eDNA_species) 


venn_data <- c(  
  "CT"        = length(only_CT),  
  "eDNA"      = length(only_eDNA),  
  "CT&eDNA"   = length(both)  
)  

# Fit proportional Euler diagram  
fit <- euler(venn_data) 

palette_colors <- c("#F9DEE0","#D3EFFC","#D6D1DD" )

Plot_CT_eDNA_species <- plot(  
  fit,  
  labels = list(  
    fontsize = 16,  
    position = "outside"  
  ),  
  quantities = list(  
    fontsize = 16  
  ),  
  edges = FALSE,  
  fills = palette_colors[1:2]   # use first 2 colors  
)

Plot_CT_eDNA_species
```
