## Supporting Information 2 for "Species ecology, behavior, and substrate type shape eDNA detection of mammals at savanna waterholes": 07ba calculating mean datalogger values.R

```
# Set wd
setwd(dirname(rstudioapi::getActiveDocumentContext()$path))

## Load required package  

library(dplyr)  
library(lubridate)  

## 1. Import data -------------------------------------------------------------  

periods <- read.csv(  
  "periods_start_end_times.csv",  
  sep = ";",  
  dec = ",",  
  stringsAsFactors = FALSE  
)  

datalogger <- read.csv(  
  "datalogger_data.csv",  
  sep = ";",  
  dec = ",",  
  stringsAsFactors = FALSE  
)  

## 2. Convert date-time columns ----------------------------------------------  

# Periods: "Start" and "End" in format "24.03.2023 07:46:00"  
periods <- periods %>%  
  mutate(  
    Start = dmy_hms(Start),  
    End   = dmy_hms(End)  
  )  

# Datalogger: "datetime" in the same format  
datalogger <- datalogger %>%  
  mutate(  
    datetime = dmy_hms(datetime)  
  )  

## 3. Compute mean temp and light per period and location ---------------------  

# For safety, ensure consistent type of "location" in both dataframes  
periods$location    <- as.character(periods$location)  
datalogger$location <- as.character(datalogger$location)  

# For each row in `periods`, find matching datalogger rows (same location and  
# datetime between Start and End), and compute mean temp and light  
periods_with_means <- periods %>%  
  rowwise() %>%  
  mutate(  
    mean_temp  = mean(datalogger$temp[datalogger$location == location &  
                                        datalogger$datetime >= Start &  
                                        datalogger$datetime <= End],  
                      na.rm = TRUE),  
    mean_light = mean(datalogger$light[datalogger$location == location &  
                                         datalogger$datetime >= Start &  
                                         datalogger$datetime <= End],  
                      na.rm = TRUE)  
  ) %>%  
  ungroup()  

## 4. Inspect result ----------------------------------------------------------  

head(periods_with_means)  
# Write csv

write.csv(periods_with_means , "temp_light_means.csv")
```
