## Supporting Information 2 for "Species ecology, behavior, and substrate type shape eDNA detection of mammals at savanna waterholes": 08 testing for influence of species properties onto detection overall.R

```
# Script for Logistic Regression on the effect of species traits on detection in the camera trapping or eDNA dataset
# Set wd
setwd(dirname(rstudioapi::getActiveDocumentContext()$path))

## Load required package  

library(dplyr)  
library(ResourceSelection)  

# Read  the  CSV files  
detections_df <- read.csv("species_presence_CT_eDNA.csv",  
                       stringsAsFactors = FALSE)  

pantheria_df <- read.csv2("Pantheria database/PanTHERIA_1-0_WR05_Aug2008.csv",  
                         stringsAsFactors = FALSE)  

## Adding species traits: 
#  Select the columns you want from PanTHERIA  
pantheria_sel <- pantheria_df %>%  
  select(  
    MSW05_Binomial,  
    `X5.1_AdultBodyMass_g`,  
    `X13.1_AdultHeadBodyLen_mm`,  
    `X10.1_PopulationGrpSize`,  
    `X10.2_SocialGrpSize`,  
    `X6.2_TrophicLevel`  
  )  

#Join:  
##    Species (in species_df) <-> MSW05_Binomial (in pantheria_df)  
detections_df <- detections_df %>%  
  left_join(  
    pantheria_sel,  
    by = c("Species" = "MSW05_Binomial")  
  )  

## 4. Optional: save the result  
write.csv(  
  detections_df,  
  "species_detections_df_CT_eDNA_with_Pantheria.csv",  
  row.names = FALSE  
)  


### Missing data was complemented with data from Kingdom Field guide and gbif

# Load the complemented csv
detections_characteristics <- read.csv2("species_detections_species_characteristics.csv",  
                          stringsAsFactors = FALSE)  


## Logistic regression on species characteristics on detection and non detection of CT and eDNA

#  Ensure TrophLev is a factor  
detections_characteristics$TrophLev <- factor(detections_characteristics$TrophLev)  


#  Standardize continuous predictors 
detections_characteristics$AdultBodyMass_sc    <- scale(detections_characteristics$AdultBodyMass)  
detections_characteristics$AdultHeadBodyLen_sc <- scale(detections_characteristics$AdultHeadBodyLen)  
detections_characteristics$SocialGrpSize_sc    <- scale(detections_characteristics$SocialGrpSize)                           


# Model for CT
m_CT <- glm(  
  CT_species ~ AdultBodyMass_sc + AdultHeadBodyLen_sc +  
    SocialGrpSize_sc + TrophLev ,  
  family = binomial,  
  data   = detections_characteristics  
)  

summary(m_CT)  


# Using ANOVA to see whether the trophic level overall has an significant effect, not just the individual levels. 
car::Anova(m_CT, type = 3)  # overall tests for each predictor  

# Quantify collinearity between body weight and body length
cor(detections_characteristics$AdultBodyMass,  
    detections_characteristics$AdultHeadBodyLen,  
    use = "complete.obs")  
m_CT_full <- glm(  
  CT_species ~ AdultBodyMass_sc + AdultHeadBodyLen_sc +  
               SocialGrpSize_sc + TrophLev,  
  family = binomial,  
  data   = detections_characteristics  
)  
  
car::vif(m_CT_full)

## Collinearity between body length and body mass is not a big issue


# Model for eDNA

# Changing the reference level for Trophic Level 
detections_characteristics$TrophLev <- relevel(detections_characteristics$TrophLev,  
                                               ref = "2")  
m_eDNA <- glm(  
  eDNA_species ~ AdultBodyMass_sc + AdultHeadBodyLen_sc +  
    SocialGrpSize_sc + TrophLev,  
  family = binomial,  
  data   = detections_characteristics  
)  

summary(m_eDNA)  
car::Anova(m_eDNA, type = 3)  


### Goodness of fit tests of models  ###############


## Ensure response variables are numeric 0/1  
detections_characteristics$CT_species   <- as.numeric(detections_characteristics$CT_species)  
detections_characteristics$eDNA_species <- as.numeric(detections_characteristics$eDNA_species)  

# If they are 0/1 coded but became 1/2 during coercion, fix that:  
if (all(na.omit(detections_characteristics$CT_species) %in% c(1, 2))) {  
  detections_characteristics$CT_species <- detections_characteristics$CT_species - 1  
}  
if (all(na.omit(detections_characteristics$eDNA_species) %in% c(1, 2))) {  
  detections_characteristics$eDNA_species <- detections_characteristics$eDNA_species - 1  
}  

## ---- CT model ----  
# Create a complete-case dataset for the CT model  
dat_CT <- detections_characteristics[complete.cases(  
  detections_characteristics[, c("CT_species",  
                                 "AdultBodyMass_sc",  
                                 "AdultHeadBodyLen_sc",  
                                 "SocialGrpSize_sc",  
                                 "TrophLev")]  
), ]  

# Refit the CT model on complete cases (ensures y and fitted have same length)  
m_CT_cc <- glm(  
  CT_species ~ AdultBodyMass_sc + AdultHeadBodyLen_sc +  
    SocialGrpSize_sc + TrophLev,  
  family = binomial,  
  data   = dat_CT  
)  
```
