## Supporting Information 2 for "Species ecology, behavior, and substrate type shape eDNA detection of mammals at savanna waterholes": 09 Data prep and modelling detections.R

```
# Script to prepare the data for GLMM modelling 

# Set wd
setwd(dirname(rstudioapi::getActiveDocumentContext()$path))

## Load required package  

library(dplyr)  
library(tidyr)  
library(stringr) 
library(purrr) 
library(lubridate) 

# Read  the  CSV files  
CT_data <- read.csv("CT_data_visitation_events_added.csv",  
                          stringsAsFactors = FALSE)  

eDNA_data_samples <- read.csv("eDNA_data_samples_incl_latin.csv",  
                    stringsAsFactors = FALSE)  %>% select(-`X`) %>% select(-`Old_name`)

eDNA_data_sampling_events <- read.csv("eDNA_data_sampling_events_incl_latin.csv",  
                              stringsAsFactors = FALSE)  %>% select(-`X`) %>% select(-`Old_name`)

# Preparation for modelling of detection events: 

# exclude all non-regular periods
eDNA_data_sampling_events_sel <- subset(eDNA_data_sampling_events, regular_period > 0)

CT_data_sel <- subset(CT_data, regular_period > 0)

eDNA_data_samples_sel <- subset(eDNA_data_samples, regular_period > 0)

### Create the abstract variable "ind_obs", which is a combination of number of pictures and number of individuals (per sp per behavior) 

# Initialize the ind_obs variable in eDNA_data_sampling_events_sel
eDNA_data_sampling_events_sel$ind_obs <- NA

# Loop through each unique combination of Species and period in eDNA_data_sampling_events_sel
for(i in 1:nrow(eDNA_data_sampling_events_sel)) {
  species <- eDNA_data_sampling_events_sel$Species[i]
  period <- eDNA_data_sampling_events_sel$period[i]
  
  # Sum the Individuals_divided_by_sp_and_behaviour in CT_data_sel where the Species and period match
  matching_sum <- sum(CT_data_sel$Individuals_divided_by_sp_and_behaviour[
    CT_data_sel$Species == species & CT_data_sel$period == period], na.rm = TRUE)
  
  # If no match is found, give a warning
  if(matching_sum == 0) {
    warning(paste("No matching combination of Species:", species, "and period:", period, "found in CT_data_inclperiod_filtered"))
  }
  
  # Assign the sum to the ind_obs column
  eDNA_data_sampling_events_sel$ind_obs[i] <- matching_sum
}

# Add the number of visitation events for this species in this period
# Calculate visitation evens
ct_summary <- CT_data_sel %>%  
  group_by(Species, period) %>%  
  summarise(  
    n_vis_events = n_distinct(vis_event),  
    .groups = "drop"  
  )

# Join with table
eDNA_data_sampling_events_joined <- eDNA_data_sampling_events_sel %>%  
  left_join(ct_summary, by = c("Species", "period"))  

# replace NAs with zeros
eDNA_data_sampling_events_joined$n_vis_events[is.na(eDNA_data_sampling_events_joined$n_vis_events)] <- 0  


write.csv(ct_summary , "visitation_events_per_period.csv")

# Add the species properties to the dataframe

# Load the complemented csv
detections_characteristics <- read.csv2("../08 LogRes of species properties influencing detection of eDNA and CT/species_detections_species_characteristics.csv",  
                                        stringsAsFactors = FALSE)  

# Join the characteristics into main data frame
eDNA_data_sampling_events_joined <- eDNA_data_sampling_events_joined %>%  
  left_join(  
    detections_characteristics %>%  
      select(Species, AdultBodyMass, AdultHeadBodyLen, SocialGrpSize, TrophLev),  
    by = "Species"  
  ) 

# Adding the environmental variables temperature, light, pH, conductivity
env_variables <- read.csv2("../07b adding env variables visitation events and ind obs/water measurements and vis events for modelling.csv",  
                                        stringsAsFactors = FALSE) 

eDNA_data_sampling_events_joined <- eDNA_data_sampling_events_joined %>%  
  left_join(  
    env_variables %>%  
      select(period, pH, Temp_mean, lux_mean, conductivity, O2),  
    by = "period"  
  ) 

# Adding behaviour
# Create the behav_num column in CT data 
CT_data_sel <- CT_data_sel  %>%
  mutate(behav_num = case_when(
    behaviour == "standingclose" ~ 1,
    behaviour == "drinking" ~ 2,
    behaviour == "feetinwater" ~ 3,
    behaviour == "feetdrink" ~ 4,
    TRUE ~ NA_real_  # Assign NA for other behaviors
  ))

# Check for any NAs in the new 'behav_num' column
if (any(is.na(CT_data_sel$behav_num))) {
  warning("NA values have been created in 'behav_num'.")
}

###Calculate percentiles and include them in the main df  . Percentile is rounded
# Step 1: Calculate the 75th percentile for each combination of species and period in CT_data_sel 
percentiles <- CT_data_sel  %>%
  group_by(Species, period) %>%
  summarize(behav_perz = round(quantile(behav_num, 0.75, na.rm = TRUE))) %>%
  ungroup()

# Step 2: Merge the percentiles with the eDNA_data_sampling_events_joined dataframe
eDNA_data_sampling_events_joined <- eDNA_data_sampling_events_joined %>%
  left_join(percentiles, by = c("Species"="Species", "period"="period"))


## Calculation of time since last visit  

# Create a variable for the time point of eDNA sampling for each period

# Create date 2 by replacing periods with underlines
eDNA_data_sampling_events_joined$date2 <- gsub("_", ".", eDNA_data_sampling_events_joined$date) 
# Replace periods with dashes in date2 and convert to Date class with the correct format (dd.mm.yyyy)
eDNA_data_sampling_events_joined$date2 <- as.Date(gsub("\\.", "-", eDNA_data_sampling_events_joined$date2), format="%d-%m-%Y")

# Combine date2 with the time 07:45 and convert to POSIXct
eDNA_data_sampling_events_joined$eDNA_sampling_timepoint <- as.POSIXct(paste(eDNA_data_sampling_events_joined$date2, "07:45"), format="%Y-%m-%d %H:%M")

# Importing the DateTime of the latest picture for each species in each period from the CT data into the eDNA data

# Create the latest_picture dataframe
latest_picture_sameperiod <- CT_data_sel %>%
  group_by(Species, period) %>%               # Group by Species and period
  filter(DateTime == max(DateTime, na.rm = TRUE)) %>%  # Filter for the latest DateTime in each group
  select(Species, period, DateTime) %>%       # Select only the relevant columns
  ungroup() 

# Ungroup the dataframe for future operations
latest_picture_sameperiod <- unique(latest_picture_sameperiod)
latest_picture_sameperiod$species <- latest_picture_sameperiod$Species

# Perform a left join to add the DateTime from latest_picture to eDNA_data_sampling_events_joined
eDNA_data_sampling_events_joined <- eDNA_data_sampling_events_joined %>%
  left_join(latest_picture_sameperiod, by = c("Species", "period")) %>%
  rename(latest_CT_picture_sameperiod = DateTime)
#Remove species column
eDNA_data_sampling_events_joined$species <- NULL


# Calculate the time difference between eDNA_sampling_timepoint and latest_CT_picture

eDNA_data_sampling_events_joined <- eDNA_data_sampling_events_joined %>%
  mutate(
    eDNA_sampling_timepoint_POSIX = ymd_hms(eDNA_sampling_timepoint),
    latest_CT_picture_sameperiod_POSIX = ymd_hms(latest_CT_picture_sameperiod)
  )

# Calculate time difference in minutes (rounded) and replace negative/zero values with 1
eDNA_data_sampling_events_joined <- eDNA_data_sampling_events_joined %>%
  mutate(
    time_since_last_visit_sameperiod = round(as.numeric(difftime(eDNA_sampling_timepoint_POSIX, latest_CT_picture_sameperiod_POSIX, units = "mins"))),
    time_since_last_visit_sameperiod = if_else(time_since_last_visit_sameperiod <= 0, 1, time_since_last_visit_sameperiod)  # Replace negatives/zero with 1
  )


## Calculate time since last visit overall 

#####start with this: 
# Create a list of periods
periods <- unique(CT_data_sel$period)

# Initialize the new dataframe with unique species
latest_pictures_periods <- CT_data_sel %>%
  select(Species) %>%
  distinct()


# Loop through each period and calculate the latest DateTime for each species
for (period in periods) {
  latest_picture_period <- CT_data_sel %>%
    filter(period == !!period) %>%
    group_by(Species) %>%
    summarize(!!period := max(DateTime, na.rm = TRUE)) %>%
    ungroup()
  
  # Join the calculated DateTime for the current period to the new dataframe
  latest_pictures_periods <- latest_pictures_periods %>%
    left_join(latest_picture_period, by = c("Species" = "Species"))
}

# Pivot the latest picture period from wide to long and add variable about season and hierarchy (= order within a period)
latest_pictures_periods_long <- latest_pictures_periods %>%  
  # 1. Pivot from wide to long: all columns from 2nd to last  
  pivot_longer(  
    cols      = 2:ncol(.),  
    names_to  = "period",  
    values_to = "date"      
  ) %>%  
  # 2. Create season and period_hierarchy from column names  
  mutate(  
    # season: based on presence of "w" or "d" in the name  
    season = case_when(  
      str_detect(period, "_w_") ~ "wet",  
      str_detect(period, "_d_") ~ "dry",  
      TRUE                                  ~ NA_character_  
    ),  
    # period_hierarchy: last two digits of the name  
    period_hierarchy = str_extract(period, "[0-9]{2}$")  
  )  


latest_pictures_overall <- latest_pictures_periods_long %>%  
  # Derive base period and hierarchy from period string  
  mutate(  
    period_base = str_sub(period, 1, nchar(period) - 2),  
    hierarchy   = as.integer(str_sub(period, -2, -1))    
  ) %>%  
  # Work within Species + period_base groups  
  arrange(Species, period_base, hierarchy) %>%  
  group_by(Species, period_base) %>%  
  mutate(  
    filled_date = {  
      last_non_na <- as.Date(NA)  
      out <- date  
      for (i in seq_along(date)) {  
        if (!is.na(date[i])) {  

          last_non_na <- date[i]  
          out[i] <- date[i]  
        } else {  
          out[i] <- last_non_na  
        }  
      }  
      out  
    }  
  ) %>%  
  ungroup()  

latest_pictures_lookup <- latest_pictures_overall %>%  
  select(Species, period, filled_date) %>%  
  rename(latest_picture_overall = filled_date)  

# Join with main df
eDNA_data_sampling_events_joined <- eDNA_data_sampling_events_joined %>%  
  left_join(latest_pictures_lookup, by = c("Species", "period"))  


# Calculate the time difference between eDNA_sampling_timepoint and latest_CT_picture
# Convert to POSIXct and create new variables
eDNA_data_sampling_events_joined <- eDNA_data_sampling_events_joined %>%
  mutate(
    eDNA_sampling_timepoint_POSIX = ymd_hms(eDNA_sampling_timepoint),
    latest_picture_overall_POSIX = ymd_hms(latest_picture_overall)
  )

# Calculate time difference in minutes (rounded) and replace negative/zero values with 1
eDNA_data_sampling_events_joined <- eDNA_data_sampling_events_joined %>%
  mutate(
    time_since_last_visit_overall = round(as.numeric(difftime(eDNA_sampling_timepoint_POSIX, latest_picture_overall_POSIX, units = "mins"))),
    time_since_last_visit_overall = if_else(time_since_last_visit_overall <= 0, 1, time_since_last_visit_overall)  # Replace negatives/zero with 1
  )


# glmm can't handle NAs. So we need to create an additional variable defining whether there was no visit in this period or overall

# Handle time-since-last-visit NAs with indicators  
eDNA_data_sampling_events_joined <- eDNA_data_sampling_events_joined %>%  
  mutate(  
    no_prev_visit_sameperiod   = ifelse(is.na(time_since_last_visit_sameperiod), 1, 0),  
    no_prev_visit_overall= ifelse(is.na(time_since_last_visit_overall),    1, 0),  
    
    # Replace NAs with 0 (or another constant) so we can still use the variable numerically  
    time_since_last_visit_sameperiod  = ifelse(is.na(time_since_last_visit_sameperiod),  0, time_since_last_visit_sameperiod),  
    time_since_last_visit_overall     = ifelse(is.na(time_since_last_visit_overall),     0, time_since_last_visit_overall)  
  )


# remove helper columns
eDNA_data_sampling_events_joined$eDNA_sampling_timepoint_POSIX <- NULL 
eDNA_data_sampling_events_joined$latest_CT_picture_sameperiod_POSIX <- NULL
eDNA_data_sampling_events_joined$latest_CT_picture_sameperiod <- NULL
eDNA_data_sampling_events_joined$latest_picture_overall_POSIX<- NULL 
eDNA_data_sampling_events_joined$latest_picture_overall<- NULL 
eDNA_data_sampling_events_joined$eDNA_sampling_timepoint<- NULL 

###timedifference is done!!


## Calculating the most frequently used sp as well as counts on all sps
# Converting sp to sp num
CT_data_sel <- CT_data_sel %>%  
  mutate(  
    sp_num = as.integer(str_remove(sp, "^sp"))  
  ) 


# For each Species + period, find the most frequent sp in CT_data_sel  
sp_mode_by_group <- CT_data_sel %>%  
  group_by(Species, period, sp_num) %>%  
  summarise(n = n(), .groups = "drop_last") %>%   # count how often each sp appears  
  slice_max(n, with_ties = FALSE) %>%             # keep the most frequent sp per Species+period  
  ungroup() %>%  
  select(Species, period, sp_most_freq = sp_num)  

# Join this info into eDNA_data_sampling_events_joined  
eDNA_data_sampling_events_joined <- eDNA_data_sampling_events_joined %>%  
  left_join(sp_mode_by_group, by = c("Species", "period"))  


## Count observation counts at each sp for each Species and period


# Count how many times each sp occurs per Species + period  
sp_counts_long <- CT_data_sel %>%  
  group_by(Species, period, sp) %>%  
  summarise(n = n(), .groups = "drop")  

# Pivot to wide format, one column per sp, named "<sp>_count"  
sp_counts_wide <- sp_counts_long %>%  
  mutate(sp_col = paste0(sp, "_count")) %>%  
  select(Species, period, sp_col, n) %>%  
  pivot_wider(  
    names_from   = sp_col,  
    values_from  = n,  
    values_fill  = 0  
  ) %>%  
  # 2b) Order the *_count columns alphabetically  
  select(  
    Species,  
    period,  
    sort(setdiff(names(.), c("Species", "period")))  
  )  

# Join the counts into eDNA_data_sampling_events_joined  
eDNA_data_sampling_events_joined <- eDNA_data_sampling_events_joined %>%  
  left_join(sp_counts_wide, by = c("Species", "period"))  


# also social group size and behav perz. have NAs, the need to be handled like the NAs in the time variables
eDNA_data_sampling_events_joined <- eDNA_data_sampling_events_joined %>%  
  mutate(  
    # Indicators for missingness  
    SocialGrpSize_missing = ifelse(is.na(SocialGrpSize), 1, 0),  
    behav_perz_missing    = ifelse(is.na(behav_perz),    1, 0),  
    
    # Replace NAs with a constant (e.g. 0 or mean), then scale  
    SocialGrpSize = ifelse(is.na(SocialGrpSize), 0, SocialGrpSize),  
    behav_perz    = ifelse(is.na(behav_perz),    0, behav_perz)  
  ) 


# Rename dataframe and export as csv

eDNA_CT_combined_sampling_events  <- eDNA_data_sampling_events_joined

write.csv(eDNA_CT_combined_sampling_events , "eDNA_CT_combined_sampling_events.csv")


# add the relevant variables also to the sample dataframe


eDNA_CT_combined_samples <- eDNA_data_samples_sel %>%  
  left_join(  
    eDNA_CT_combined_sampling_events %>%  
      select(  
        Species,  
        period,
        date2,
        ind_obs,  
        AdultBodyMass,  
        AdultHeadBodyLen,  
        SocialGrpSize,
        SocialGrpSize_missing,
        TrophLev,
        n_vis_events,
        pH,
        Temp_mean,
        lux_mean,
        conductivity,
        O2,
        behav_perz,  
        behav_perz_missing,
        time_since_last_visit_sameperiod,  
        time_since_last_visit_overall,
        no_prev_visit_sameperiod,
        no_prev_visit_overall,
        sp_most_freq,
        sp1_count,
        sp2_count,
        sp3_count,
        sp4_count, 
        sp5_count,
        sp6_count,
        sp7_count,
        sp8_count
      ),  
    by = c("Species", "period")  
  )  


write.csv(eDNA_CT_combined_samples , "eDNA_CT_combined_samples.csv")
```
