## Supporting Information 2 for "Species ecology, behavior, and substrate type shape eDNA detection of mammals at savanna waterholes": splitting_sampling_event_data_by_substrate.R

```
# Set working directory
setwd(dirname(rstudioapi::getActiveDocumentContext()$path))

library(dplyr)

# Import data
df <- read.csv("eDNA_CT_combined_samples.csv", stringsAsFactors = FALSE)


### ------------------------------------------------------------
### SPLIT BY MEDIUM
### ------------------------------------------------------------

df_water <- df %>% filter(medium == "water")
df_sediment <- df %>% filter(medium == "sediment")

### ------------------------------------------------------------
### FUNCTION: DEREPLICATION PIPELINE
### ------------------------------------------------------------

dereplicate_df <- function(data) {
  
  data %>%
    group_by(period, Species) %>%
    summarise(
      number.of.samples = n(),
      num_detection = sum(detection == 1, na.rm = TRUE),
      
      across(
        -c(X, sample.name, medium, sampling.spot, detection),
        ~first(.)
      ),
      
      .groups = "drop"
    ) %>%
    mutate(
      detection = ifelse(num_detection > 0, 1, 0)
    )
}

### ------------------------------------------------------------
### APPLY DEREPLICATION and add medium column
### ------------------------------------------------------------

df_water_summary <- dereplicate_df(df_water) %>%
  mutate(medium = "water")

df_sediment_summary <- dereplicate_df(df_sediment) %>%
  mutate(medium = "sediment")


### ------------------------------------------------------------
### EXPORT RESULTS
### ------------------------------------------------------------


df_combined <- bind_rows(df_water_summary, df_sediment_summary)

write.csv(df_combined,
          "eDNA_CT_combined_sampling_events_split_by_substrate.csv",
          row.names = FALSE)
```
