## Supporting Information 2 for "Species ecology, behavior, and substrate type shape eDNA detection of mammals at savanna waterholes": 10_modelling.R

```
# Script for modelling model influence of CT (and species trait) predictors onto eDNA dectection

# Set wd
setwd(dirname(rstudioapi::getActiveDocumentContext()$path))

## Load required package  

library(dplyr)  
library(tidyr)  
library(stringr) 
library(purrr) 
library(lubridate) 
library(lme4)
library(DHARMa)

# Read  the  CSV files  
data_sampling_events <- read.csv("eDNA_CT_combined_sampling_events.csv", )  


# Ensure that factor predictors are factors and scale continuous variables

data_sampling_events <- data_sampling_events %>%  
  mutate(  
    # Factors  
    Species  = factor(Species),  
    location = factor(location),  
    period   = factor(period),  
    season   = factor(season, levels = c("dry", "wet")),  # adjust reference if needed  
    TrophLev = factor(TrophLev),  
    
    # Scale species traits  
    AdultBodyMass_sc    = as.numeric(scale(AdultBodyMass)),  
    AdultHeadBodyLen_sc = as.numeric(scale(AdultHeadBodyLen)),  
    SocialGrpSize_sc    = as.numeric(scale(SocialGrpSize)),  
    
    # Scale activity metrics
    ind_obs_sc          = as.numeric(scale(ind_obs)),  
    n_vis_sc            = as.numeric(scale(n_vis_events)), 
    time_same_sc        = as.numeric(scale(time_since_last_visit_sameperiod)),  
    time_overall_sc     = as.numeric(scale(time_since_last_visit_overall)),
    behav_perz_sc       = as.numeric(scale(behav_perz)),  

    #Scale environmental variables
    Temp_mean_sc            = as.numeric(scale(Temp_mean)),
    lux_sc            = as.numeric(scale(lux_mean)),
    cond_sc            = as.numeric(scale(conductivity)),
    O2_sc            = as.numeric(scale(O2)),
    pH_sc            = as.numeric(scale(pH)),
    
  ) 


# 1.b Modelling the effects of species traits, activity parameters,  and environmental variables onto positive samples of species

m_full <- glmer(  
  cbind(num_detection, number.of.samples - num_detection) ~  
    # Species traits
    AdultBodyMass_sc +  
    AdultHeadBodyLen_sc +  
    SocialGrpSize_sc +  
    SocialGrpSize_missing +  
    TrophLev + 
    # Activity metrics
    ind_obs_sc +  
    n_vis_sc +
    time_same_sc +  
    time_overall_sc + 
    no_prev_visit_sameperiod +  
    no_prev_visit_overall + 
    behav_perz_sc +  
    behav_perz_missing +  
    # Environmental variables
    season +
    Temp_mean_sc +
    pH_sc +
    lux_sc +
    cond_sc +
    O2_sc +
    (1 | Species) +  
    (1 | location) +  
    (1 | period),  
  data   = data_sampling_events,  
  family = binomial(link = "logit"),  
  control = glmerControl(  
    optimizer = "bobyqa",  
    optCtrl   = list(maxfun = 1e5)  
  )  
)  

summary(m_full)  

# Dropping the  behav_perz_missing and SocialGrpSize_missing indicator because they are fully correlated with Species

m_reduced <- glmer(  
  cbind(num_detection, number.of.samples - num_detection) ~  
    # Species traits
    AdultBodyMass_sc +  
    AdultHeadBodyLen_sc +  
    SocialGrpSize_sc +  
    TrophLev +  
    # Activity metrics
    n_vis_sc +
    ind_obs_sc +  
    time_same_sc +  
    time_overall_sc +  
    no_prev_visit_sameperiod +  
    no_prev_visit_overall +  
    behav_perz_sc +  
    # Environmental variables
    season +  
    Temp_mean_sc +
    pH_sc +
    lux_sc +
    cond_sc +
    O2_sc +
    (1 | Species) +  
    (1 | location) +  
    (1 | period),  
  data   = data_sampling_events,  
  family = binomial(link = "logit"),  
  control = glmerControl(  
    optimizer = "bobyqa",  
    optCtrl   = list(maxfun = 1e5)  
  )  
)  

summary(m_reduced)  


res_rate <- simulateResiduals(m_reduced, n = 1000)  
plot(res_rate)  
testDispersion(res_rate)  
testZeroInflation(res_rate) 


## Stepwise simplification of model 

### removing gorups of predictors that have little to no influence:
# 1. Environemntal variables
m1 <- update(m_reduced,  
             . ~ . - Temp_mean_sc - lux_sc - cond_sc - O2_sc - pH_sc)  


AIC(m_reduced, m1)  
anova(m_reduced, m1, test = "Chisq")  

# M1 is new working model
# 2. season
m2 <- update(m1, . ~ . - season)  

AIC(m1, m2)  
anova(m1, m2, test = "Chisq")  

# M2 is new working model
m3 <- update(m2, . ~ . - AdultBodyMass_sc - AdultHeadBodyLen_sc)  

AIC(m2, m3)  
anova(m2, m3, test = "Chisq") 

# M3 is new working model
m4 <- update(m3, . ~ . - time_same_sc)  

AIC(m3, m4)  
anova(m3, m4, test = "Chisq")  

# M4 is the final model

m_final <- m4

summary(m_final)  

res_rate <- simulateResiduals(m_final, n = 1000)  
plot(res_rate)  
testDispersion(res_rate)  
testZeroInflation(res_rate) 
```
