## Supporting Information 2 for "Species ecology, behavior, and substrate type shape eDNA detection of mammals at savanna waterholes": 10a_modelling_split_by_substrate.R

```
# Script for modelling model influence of CT (and species trait) predictors onto eDNA detection with assessing the effect for sediment and water

# Set wd
setwd(dirname(rstudioapi::getActiveDocumentContext()$path))

## Load required package  

library(dplyr)  
library(tidyr)  
library(stringr) 
library(purrr) 
library(lubridate) 
library(lme4)
library(DHARMa)

# Read  the  CSV files  
data_sampling_events_split_by_medium <- read.csv("eDNA_CT_combined_sampling_events_split_by_substrate.csv", )  


# Ensure that factor predictors are factors and scale continuous variables

data_sampling_events_split_by_medium <- data_sampling_events_split_by_medium %>%  
  mutate(  
    # Factors  
    Species  = factor(Species),  
    location = factor(location),  
    period   = factor(period),  
    season   = factor(season, levels = c("dry", "wet")),  # adjust reference if needed  
    TrophLev = factor(TrophLev),  
    
    # Scale species traits  
    AdultBodyMass_sc    = as.numeric(scale(AdultBodyMass)),  
    AdultHeadBodyLen_sc = as.numeric(scale(AdultHeadBodyLen)),  
    SocialGrpSize_sc    = as.numeric(scale(SocialGrpSize)),  
    
    # Scale activity metrics
    ind_obs_sc          = as.numeric(scale(ind_obs)),  
    n_vis_sc            = as.numeric(scale(n_vis_events)), 
    time_same_sc        = as.numeric(scale(time_since_last_visit_sameperiod)),  
    time_overall_sc     = as.numeric(scale(time_since_last_visit_overall)),
    behav_perz_sc       = as.numeric(scale(behav_perz)),  

    #Scale environmental variables
    Temp_mean_sc            = as.numeric(scale(Temp_mean)),
    lux_sc            = as.numeric(scale(lux_mean)),
    cond_sc            = as.numeric(scale(conductivity)),
    O2_sc            = as.numeric(scale(O2)),
    pH_sc            = as.numeric(scale(pH)),
    
  ) 


# Modelling the effects of species traits, activity parameters,  and environmental variables onto positive samples of species now with an interaction term "medium"

m_full_split_by_medium <- glmer(
  
  cbind(num_detection, number.of.samples - num_detection) ~
    
    medium * (
      
      # Species traits
      AdultBodyMass_sc +
        AdultHeadBodyLen_sc +
        SocialGrpSize_sc +
        TrophLev +
        
        # Activity metrics
        ind_obs_sc +
        n_vis_sc +
        time_same_sc +
        time_overall_sc +
        no_prev_visit_sameperiod +
        no_prev_visit_overall +
        behav_perz_sc +
        
        # Environmental variables
        Temp_mean_sc +
        pH_sc +
        lux_sc +
        cond_sc +
        O2_sc
    ) +
    
    season +
    
    (1 | Species) +
    (1 | location) +
    (1 | period),
  
  data = data_sampling_events_split_by_medium,
  family = binomial(link = "logit"),
  control = glmerControl(
    optimizer = "bobyqa",
    optCtrl = list(maxfun = 1e5)
  )
)

summary(m_full_split_by_medium)  


## Simplifying the model

m_reduced_split_by_medium <- glmer(
  
  cbind(num_detection, number.of.samples - num_detection) ~
    
    medium * (
      SocialGrpSize_sc +
        TrophLev +
        ind_obs_sc +
        behav_perz_sc +
        Temp_mean_sc +
        pH_sc +
        lux_sc +
        cond_sc +
        O2_sc
    ) +
    
    n_vis_sc +
    time_overall_sc +
    no_prev_visit_overall +
    

    (1 | Species) +
    (1 | location) +
    (1 | period),
  
  data = data_sampling_events_split_by_medium,
  
  family = binomial(link = "logit"),
  
  control = glmerControl(
    optimizer = "bobyqa",
    optCtrl = list(maxfun = 1e5)
  )
)

summary(m_reduced_split_by_medium)

# Comparing the full and reduced model 

anova(m_full_split_by_medium, m_reduced_split_by_medium, test = "Chisq")


# Difference is significant! We should keep the full model!


# DHARMA diagnostics for full model

res_rate <- simulateResiduals(m_full_split_by_medium, n = 1000)  
plot(res_rate)  
testDispersion(res_rate)  
testZeroInflation(res_rate) 

summary(m_full_split_by_medium)  
```
