## Supporting Information 2 for "Species ecology, behavior, and substrate type shape eDNA detection of mammals at savanna waterholes": 11 spatial GLMM modelling.R

```
# Script for spatial GLMM modelling (testing for an effect of sampling point location)


# Set wd
setwd(dirname(rstudioapi::getActiveDocumentContext()$path))
# Load packages
library(lme4) 
library(dplyr)  
library(tidyr) 
library(DHARMa)

# Load csv
dat <- read.csv("eDNA_CT_combined_samples.csv ", 
                    stringsAsFactors = FALSE) 

# Calculate counts for same, neighboring and local sps
dat <- dat %>%  
  mutate(  
    # same interaction count: sp<sampling.spot>_count  
    same_count = case_when(  
      sampling.spot == 1 ~ sp1_count,  
      sampling.spot == 2 ~ sp2_count,  
      sampling.spot == 3 ~ sp3_count,  
      sampling.spot == 4 ~ sp4_count,  
      sampling.spot == 5 ~ sp5_count,  
      sampling.spot == 6 ~ sp6_count,  
      sampling.spot == 7 ~ sp7_count,  
      sampling.spot == 8 ~ sp8_count,  
      TRUE ~ NA_real_  
    ),  
    # neighbours in circular arrangement  
    left_neigh = case_when(  
      sampling.spot == 1 ~ sp8_count,  
      sampling.spot == 2 ~ sp1_count,  
      sampling.spot == 3 ~ sp2_count,  
      sampling.spot == 4 ~ sp3_count,  
      sampling.spot == 5 ~ sp4_count,  
      sampling.spot == 6 ~ sp5_count,  
      sampling.spot == 7 ~ sp6_count,  
      sampling.spot == 8 ~ sp7_count,  
      TRUE ~ NA_real_  
    ),  
    right_neigh = case_when(  
      sampling.spot == 1 ~ sp2_count,  
      sampling.spot == 2 ~ sp3_count,  
      sampling.spot == 3 ~ sp4_count,  
      sampling.spot == 4 ~ sp5_count,  
      sampling.spot == 5 ~ sp6_count,  
      sampling.spot == 6 ~ sp7_count,  
      sampling.spot == 7 ~ sp8_count,  
      sampling.spot == 8 ~ sp1_count,  
      TRUE ~ NA_real_  
    ),  
    neigh_count = left_neigh + right_neigh,  
    local_count  = same_count + neigh_count  
  )  

#Replacing NAs with 0
dat <- dat %>%  
  mutate(  
    same_count  = replace_na(same_count, 0),  
    local_count = replace_na(local_count, 0)  
  )  

# Make sure predictors are factor & scale continuous predictors
dat <- dat %>%  
  mutate(  
    # Factors  
    Species  = factor(Species),  
    location = factor(location),  
    period   = factor(period),  
    season   = factor(season, levels = c("dry", "wet")),  # adjust reference if needed  
    TrophLev = factor(TrophLev),  
    
    # Scale species traits  
    AdultBodyMass_sc    = as.numeric(scale(AdultBodyMass)),  
    AdultHeadBodyLen_sc = as.numeric(scale(AdultHeadBodyLen)),  
    SocialGrpSize_sc    = as.numeric(scale(SocialGrpSize)),  
    
    # Scale behavioral predictors  
    ind_obs_sc          = as.numeric(scale(ind_obs)),  
    behav_perz_sc       = as.numeric(scale(behav_perz)),  
    
    # Scale time-since-last-visit variables  
    time_same_sc        = as.numeric(scale(time_since_last_visit_sameperiod)),  
    time_overall_sc     = as.numeric(scale(time_since_last_visit_overall)),
    n_vis_sc            = as.numeric(scale(n_vis_events)), 
    
    #Scale environmental predictors
    Temp_mean_sc        = as.numeric(scale(Temp_mean)),
    lux_sc              = as.numeric(scale(lux_mean)),
    cond_sc             = as.numeric(scale(conductivity)),
    O2_sc               = as.numeric(scale(O2)),
    pH_sc               = as.numeric(scale(pH)),    
    # Scale sp count data
    neigh_count_sc      = as.numeric(scale(neigh_count)),
    local_count_sc      = as.numeric(scale(local_count)),
    same_count_sc      = as.numeric(scale(same_count)),
  ) 


# Base model without spatial distribution terms  
m_no_spatial <- glmer(  
  detection ~   
    AdultBodyMass_sc +  
    AdultHeadBodyLen_sc +  
    SocialGrpSize_sc +  
    TrophLev +  
    ind_obs_sc +  
    behav_perz_sc +  
    time_same_sc +  
    time_overall_sc +  
    no_prev_visit_sameperiod +  
    no_prev_visit_overall +  
    n_vis_sc +
    season +  
    Temp_mean_sc +
    pH_sc +
    lux_sc +
    cond_sc +
    O2_sc +
    (1 | Species) +  
    (1 | location) +  
    (1 | period),    
  data = dat,  
  family = binomial,  
  control = glmerControl(optimizer = "bobyqa")  
)  

summary(m_no_spatial)
res_no_spatial <- simulateResiduals(m_no_spatial, n = 1000)  
plot(res_no_spatial)  
testDispersion(res_no_spatial)  
testZeroInflation(res_no_spatial) 

# Model with spatial terms  
m_spatial <- glmer(  
  detection ~   
    AdultBodyMass_sc +  
    AdultHeadBodyLen_sc +  
    SocialGrpSize_sc +  
    TrophLev +  
    ind_obs_sc +  
    behav_perz_sc +  
    time_same_sc +  
    time_overall_sc +  
    no_prev_visit_sameperiod +  
    no_prev_visit_overall +  
    n_vis_sc +
    season +  
    Temp_mean_sc +
    pH_sc +
    lux_sc +
    cond_sc +
    O2_sc +
    local_count_sc +
    same_count_sc +
    (1 | Species) +  
    (1 | location) +  
    (1 | period),  
  data = dat,  
  family = binomial,  
  control = glmerControl(optimizer = "bobyqa")  
)  

summary(m_spatial)

res_spatial <- simulateResiduals(m_spatial, n = 1000)  
plot(res_spatial)  
testDispersion(res_spatial)  
testZeroInflation(res_spatial) 

anova(m_no_spatial, m_spatial, test = "Chisq")  
```
