## Supporting Information 2 for "Species ecology, behavior, and substrate type shape eDNA detection of mammals at savanna waterholes": 12 calculate missing camera trap time of surveys.R

```
# Script for calculating the camera trap times 

## ------------------------------------------------------------  
## Interactive script: camera‑trapping effort per survey  
## ------------------------------------------------------------  

# Helper: read datetime from user  
read_datetime <- function(prompt_text, format = "%Y-%m-%d %H:%M") {  
  repeat {  
    cat(prompt_text, "(format:", format, ")\n> ")  
    x <- trimws(readline())  
    dt <- as.POSIXct(x, format = format, tz = "UTC")  
    if (!is.na(dt)) return(dt)  
    cat("Could not parse date-time. Please try again.\n\n")  
  }  
}  

# Helper: read yes/no  
read_yes_no <- function(prompt_text) {  
  repeat {  
    cat(prompt_text, "(y/n)\n> ")  
    x <- tolower(trimws(readline()))  
    if (x %in% c("y", "yes"))  return(TRUE)  
    if (x %in% c("n", "no"))   return(FALSE)  
    cat("Please answer y or n.\n\n")  
  }  
}  

results_list  <- list()  
survey_index  <- 1  

cat("Camera‑trapping survey effort calculator\n")  
cat("========================================\n\n")  

repeat {  
  cat("\n--- Survey", survey_index, "---------------------------------\n")  
  cat("Enter survey name (or leave empty to finish):\n> ")  
  survey_name <- trimws(readline())  
  
  if (survey_name == "") {  
    cat("No survey name entered. Finishing input.\n")  
    break  
  }  
  
  # Start / end of survey  
  start_dt <- read_datetime("Enter survey START date-time")  
  end_dt   <- read_datetime("Enter survey END date-time")  
  
  if (end_dt <= start_dt) {  
    cat("WARNING: End time is not after start time. Please re-enter this survey.\n\n")  
    next  
  }  
  
  total_secs    <- as.numeric(difftime(end_dt, start_dt, units = "secs"))  
  total_days    <- total_secs / (60 * 60 * 24)  
  total_minutes <- total_secs / 60  
  
  # Missing periods  
  missing_periods <- list()  
  if (read_yes_no("Are there any missing periods (camera not working)?")) {  
    repeat {  
      cat("\n--- Missing period", length(missing_periods) + 1, "-------------------\n")  
      miss_start <- read_datetime("  Enter MISSING PERIOD START date-time")  
      miss_end   <- read_datetime("  Enter MISSING PERIOD END date-time")  
      
      if (miss_end <= miss_start) {  
        cat("  WARNING: Missing period end is not after start. Ignoring this period.\n")  
      } else if (miss_start >= end_dt || miss_end <= start_dt) {  
        cat("  WARNING: Missing period lies outside the survey interval. Ignoring.\n")  
      } else {  
        miss_start_clipped <- max(miss_start, start_dt)  
        miss_end_clipped   <- min(miss_end, end_dt)  
        
        missing_periods[[length(missing_periods) + 1]] <-  
          c(start = miss_start_clipped, end = miss_end_clipped)  
        
        cat("  Missing period recorded from", as.character(miss_start_clipped),  
            "to", as.character(miss_end_clipped), "\n")  
      }  
      
      if (!read_yes_no("Do you want to add another missing period?")) break  
    }  
  }  
  
  # Compute total missing time (merge overlaps)  
  total_missing_secs <- 0  
  if (length(missing_periods) > 0) {  
    mp <- do.call(rbind, missing_periods)  
    starts <- as.numeric(mp[, "start"])  
    ends   <- as.numeric(mp[, "end"])  
    o <- order(starts)  
    starts <- starts[o]  
    ends   <- ends[o]  
    
    merged_starts <- c(starts[1])  
    merged_ends   <- c(ends[1])  
    
    if (length(starts) > 1) {  
      for (i in 2:length(starts)) {  
        if (starts[i] <= merged_ends[length(merged_ends)]) {  
          merged_ends[length(merged_ends)] <- max(merged_ends[length(merged_ends)], ends[i])  
        } else {  
          merged_starts <- c(merged_starts, starts[i])  
          merged_ends   <- c(merged_ends, ends[i])  
        }  
      }  
    }  
    
    total_missing_secs <- sum(merged_ends - merged_starts)  
  }  
  
  total_missing_days    <- total_missing_secs / (60 * 60 * 24)  
  total_missing_minutes <- total_missing_secs / 60  
  
  net_secs    <- total_secs - total_missing_secs  
  net_days    <- net_secs / (60 * 60 * 24)  
  net_minutes <- net_secs / 60  
  
  perc_missing <- if (total_secs > 0) {  
    100 * total_missing_secs / total_secs  
  } else {  
    NA_real_  
  }  
  
  # Store result  
  results_list[[survey_index]] <- data.frame(  
    Survey           = survey_name,  
    Start            = start_dt,  
    End              = end_dt,  
    Total_days       = total_days,  
    Total_minutes    = total_minutes,  
    Missing_days     = total_missing_days,  
    Missing_minutes  = total_missing_minutes,  
    Net_days         = net_days,  
    Net_minutes      = net_minutes,  
    Perc_missing     = perc_missing,  
    stringsAsFactors = FALSE  
  )  
  
  cat("\nSurvey", survey_name, "processed.\n")  
  
  if (!read_yes_no("Do you want to enter another survey?")) break  
  survey_index <- survey_index + 1  
}  

# Combine all results  
if (length(results_list) > 0) {  
  survey_effort <- do.call(rbind, results_list)  
  cat("\nAll surveys processed. Resulting data frame: 'survey_effort'\n\n")  
  print(survey_effort)  
  
  ## ---- NEW: save to CSV -----------------------------------------------  
  # Choose a file name (change path if you want)  
  out_file <- "camera_trap_survey_effort.csv"  
  
  # Write the data frame to CSV (without row names)  
  write.csv(survey_effort, file = out_file, row.names = FALSE)  
  
  cat("\nSurvey effort table saved to:", normalizePath(out_file), "\n")  
  ## ---------------------------------------------------------------------  
  
} else {  
  cat("\nNo surveys were entered.\n")  
}  


### Correction of errors ########

## ============================================================  
## ADD-ON: Interactive editor for survey names and times  
## ============================================================  

if (exists("survey_effort") && nrow(survey_effort) > 0) {  
  
  cat("\n============================================================\n")  
  cat("Survey editor\n")  
  cat("You can now correct survey names, start/end times,\n")  
  cat("and missing periods (camera not working) for each survey.\n")  
  cat("============================================================\n\n")  
  
  # Keep a copy of the original for reference if needed  
  survey_effort_original <- survey_effort  
  
  repeat {  
    
    cat("\nCurrent survey table:\n")  
    print(survey_effort)  
    cat("\n")  
    
    # Ask if user wants to edit anything  
    if (!read_yes_no("Do you want to EDIT any survey?")) {  
      cat("No further edits requested. Exiting editor.\n")  
      break  
    }  
    
    # Show the index and survey names for easier reference  
    cat("\nAvailable surveys:\n")  
    for (i in seq_len(nrow(survey_effort))) {  
      cat("  [", i, "]", survey_effort$Survey[i],  
          " (", format(survey_effort$Start[i]),  
          " to ", format(survey_effort$End[i]), ")\n", sep = "")  
    }  
    cat("\n")  
    
    # Choose survey index to edit  
    repeat {  
      cat("Enter the NUMBER of the survey you want to edit:\n> ")  
      idx_input <- trimws(readline())  
      
      if (idx_input == "" || is.na(suppressWarnings(as.integer(idx_input)))) {  
        cat("Please enter a valid integer index.\n\n")  
        next  
      }  
      
      idx <- as.integer(idx_input)  
      if (idx < 1 || idx > nrow(survey_effort)) {  
        cat("Index out of range. Must be between 1 and", nrow(survey_effort), "\n\n")  
      } else {  
        break  
      }  
    }  
    
    # Display chosen survey  
    cat("\nYou selected survey [", idx, "]:\n", sep = "")  
    print(survey_effort[idx, , drop = FALSE])  
    cat("\n")  
    
    ## ---- Edit survey name -----------------------------------  
    if (read_yes_no("Do you want to EDIT the survey NAME?")) {  
      cat("Enter new survey name (leave empty to keep current: '",  
          survey_effort$Survey[idx], "'):\n> ", sep = "")  
      new_name <- trimws(readline())  
      
      if (new_name != "") {  
        survey_effort$Survey[idx] <- new_name  
        cat("Survey name updated to: '", new_name, "'\n", sep = "")  
      } else {  
        cat("Survey name left unchanged.\n")  
      }  
    }  
    
    ## ---- Edit start/end times -------------------------------  
    edit_times <- read_yes_no("Do you want to EDIT the survey START/END times?")  
    if (edit_times) {  
      
      # Show current values  
      cat("\nCurrent START:", format(survey_effort$Start[idx]), "\n")  
      cat("Current END  :", format(survey_effort$End[idx]), "\n\n")  
      
      # Read new times, but allow user to keep existing  
      cat("Enter new START date-time (or press ENTER to keep current):\n")  
      cat("Format: %Y-%m-%d %H:%M  (example: 2024-06-01 12:00)\n> ")  
      start_input <- trimws(readline())  
      
      if (start_input == "") {  
        new_start <- survey_effort$Start[idx]  
      } else {  
        new_start <- as.POSIXct(start_input, format = "%Y-%m-%d %H:%M", tz = "UTC")  
        if (is.na(new_start)) {  
          cat("Could not parse new start time. Keeping previous start.\n")  
          new_start <- survey_effort$Start[idx]  
        }  
      }  
      
      cat("\nEnter new END date-time (or press ENTER to keep current):\n")  
      cat("Format: %Y-%m-%d %H:%M  (example: 2024-06-10 18:30)\n> ")  
      end_input <- trimws(readline())  
      
      if (end_input == "") {  
        new_end <- survey_effort$End[idx]  
      } else {  
        new_end <- as.POSIXct(end_input, format = "%Y-%m-%d %H:%M", tz = "UTC")  
        if (is.na(new_end)) {  
          cat("Could not parse new end time. Keeping previous end.\n")  
          new_end <- survey_effort$End[idx]  
        }  
      }  
      
      if (new_end <= new_start) {  
        cat("\nWARNING: New END time is not after START time.\n",  
            "Times NOT changed for this survey.\n\n", sep = "")  
      } else {  
        survey_effort$Start[idx] <- new_start  
        survey_effort$End[idx]   <- new_end  
        cat("Start/End times updated for this survey.\n")  
      }  
    }  
    
    ## ---- Re-enter missing periods for this survey -----------  
    # We will ask the user if they want to re-enter missing periods.  
    # These will be used to recompute Missing_days, Net_days, etc.  
    
    if (read_yes_no("Do you want to RE-ENTER missing periods for this survey?")) {  
      
      # Use current start/end  
      start_dt <- survey_effort$Start[idx]  
      end_dt   <- survey_effort$End[idx]  
      
      if (end_dt <= start_dt) {  
        cat("ERROR: Survey END is not after START. Cannot enter missing periods.\n")  
      } else {  
        
        total_secs <- as.numeric(difftime(end_dt, start_dt, units = "secs"))  
        
        # Collect missing periods  
        missing_periods <- list()  
        
        if (read_yes_no("Are there any missing periods (camera not working) for this survey?")) {  
          
          repeat {  
            cat("\n--- Missing period", length(missing_periods) + 1, "-------------------\n")  
            miss_start <- read_datetime(" Enter MISSING PERIOD START date-time")  
            miss_end   <- read_datetime(" Enter MISSING PERIOD END date-time")  
            
            if (miss_end <= miss_start) {  
              cat(" WARNING: Missing period end is not after start. Ignoring this period.\n")  
            } else if (miss_start >= end_dt || miss_end <= start_dt) {  
              cat(" WARNING: Missing period lies outside the survey interval. Ignoring.\n")  
            } else {  
              miss_start_clipped <- max(miss_start, start_dt)  
              miss_end_clipped   <- min(miss_end, end_dt)  
              
              missing_periods[[length(missing_periods) + 1]] <-  
                c(start = miss_start_clipped, end = miss_end_clipped)  
              
              cat(" Missing period recorded from", as.character(miss_start_clipped),  
                  "to", as.character(miss_end_clipped), "\n")  
            }  
            
            if (!read_yes_no("Do you want to add another missing period?")) break  
          }  
        }  
        
        # Compute total missing time (merge overlaps)  
        total_missing_secs <- 0  
        
        if (length(missing_periods) > 0) {  
          mp <- do.call(rbind, missing_periods)  
          starts <- as.numeric(mp[, "start"])  
          ends   <- as.numeric(mp[, "end"])  
          o <- order(starts)  
          starts <- starts[o]  
          ends   <- ends[o]  
          
          merged_starts <- c(starts[1])  
          merged_ends   <- c(ends[1])  
          
          if (length(starts) > 1) {  
            for (i in 2:length(starts)) {  
              if (starts[i] <= merged_ends[length(merged_ends)]) {  
                # overlap -> extend last  
                merged_ends[length(merged_ends)] <-  
                  max(merged_ends[length(merged_ends)], ends[i])  
              } else {  
                # no overlap -> new segment  
                merged_starts <- c(merged_starts, starts[i])  
                merged_ends   <- c(merged_ends,   ends[i])  
              }  
            }  
          }  
          
          total_missing_secs <- sum(merged_ends - merged_starts)  
        }  
        
        total_missing_days    <- total_missing_secs / (60 * 60 * 24)  
        total_missing_minutes <- total_missing_secs / 60  
        
        net_secs    <- total_secs - total_missing_secs  
        net_days    <- net_secs / (60 * 60 * 24)  
        net_minutes <- net_secs / 60  
        
        perc_missing <- if (total_secs > 0) {  
          100 * total_missing_secs / total_secs  
        } else {  
          NA_real_  
        }  
        
        # Update this survey's row  
        survey_effort$Total_days[idx]       <- total_secs / (60 * 60 * 24)  
        survey_effort$Total_minutes[idx]    <- total_secs / 60  
        survey_effort$Missing_days[idx]     <- total_missing_days  
        survey_effort$Missing_minutes[idx]  <- total_missing_minutes  
        survey_effort$Net_days[idx]         <- net_days  
        survey_effort$Net_minutes[idx]      <- net_minutes  
        survey_effort$Perc_missing[idx]     <- perc_missing  
        
        cat("\nMissing‑period information updated for this survey.\n")  
      }  
    } else {  
      # If missing periods are *not* re-entered but Start/End changed,  
      # we still need to recompute totals and net based on existing values.  
      # Note: we have no record of original missing periods here,  
      # so we must assume Missing_* and Perc_missing are now invalid unless  
      # the user explicitly re-entered them.  
      if (edit_times) {  
        cat("\nNOTE: You changed START/END times but did NOT re-enter missing periods.\n")  
        cat("      Missing_days, Missing_minutes, and Perc_missing MAY be inconsistent.\n")  
        cat("      Consider re-entering missing periods for this survey later.\n")  
      }  
    }  
    
    cat("\nUpdated entry for this survey:\n")  
    print(survey_effort[idx, , drop = FALSE])  
    
    # Loop back to possibly edit another survey  
  }  
  
  # After editing, offer to save updated CSV  
  if (read_yes_no("\nDo you want to SAVE the UPDATED survey table to CSV?")) {  
    cat("Enter file name for updated CSV (leave empty to overwrite 'camera_trap_survey_effort.csv'):\n> ")  
    new_file <- trimws(readline())  
    if (new_file == "") {  
      out_file_updated <- "camera_trap_survey_effort.csv"  
    } else {  
      out_file_updated <- new_file  
    }  
    
    write.csv(survey_effort, file = out_file_updated, row.names = FALSE)  
    cat("Updated survey effort table saved to:", normalizePath(out_file_updated), "\n")  
  } else {  
    cat("Updated table NOT saved. You can still save it manually from 'survey_effort'.\n")  
  }  
  
} else {  
  cat("\nNo survey_effort object found or it has zero rows. Nothing to edit.\n")  
}  


############# Recompiling  #############
## ============================================================  
## ADD-ON: Recompute all derived columns in survey_effort  
## ============================================================  

if (exists("survey_effort") && nrow(survey_effort) > 0) {  
  
  cat("\n============================================================\n")  
  cat("Recompute survey-effort metrics\n")  
  cat("This will recompute Total_* and Net_* columns for ALL surveys\n")  
  cat("based on current Start and End times.\n")  
  cat("Existing Missing_* and Perc_missing values will be overwritten.\n")  
  cat("============================================================\n\n")  
  
  print(survey_effort)  
  cat("\n")  
  
  if (read_yes_no("Do you want to RECOMPUTE all metrics now?")) {  
    
    # Ask user how to treat missing periods  
    cat("\nYou have two options for missing periods:\n")  
    cat("  1) Assume NO missing time for any survey\n")  
    cat("  2) Use a SINGLE global percentage of missing time for all surveys\n\n")  
    
    choice <- NA_integer_  
    repeat {  
      cat("Enter 1 or 2:\n> ")  
      inp <- trimws(readline())  
      if (inp %in% c("1", "2")) {  
        choice <- as.integer(inp)  
        break  
      } else {  
        cat("Please enter either 1 or 2.\n\n")  
      }  
    }  
    
    global_perc_missing <- 0  
    
    if (choice == 1) {  
      cat("\nYou selected: assume NO missing time for any survey.\n")  
      global_perc_missing <- 0  
    } else if (choice == 2) {  
      cat("\nYou selected: use a SINGLE global percentage missing for all surveys.\n")  
      repeat {  
        cat("Enter percentage of missing time (0–100), e.g. 5 for 5%:\n> ")  
        p_inp <- trimws(readline())  
        p_val <- suppressWarnings(as.numeric(p_inp))  
        if (!is.na(p_val) && p_val >= 0 && p_val <= 100) {  
          global_perc_missing <- p_val  
          break  
        } else {  
          cat("Please enter a numeric value between 0 and 100.\n\n")  
        }  
      }  
    }  
    
    # Loop over rows and recompute  
    for (i in seq_len(nrow(survey_effort))) {  
      
      start_dt <- survey_effort$Start[i]  
      end_dt   <- survey_effort$End[i]  
      
      if (is.na(start_dt) || is.na(end_dt) || end_dt <= start_dt) {  
        cat("WARNING: Invalid Start/End for row", i,  
            "- setting all derived metrics to NA.\n")  
        survey_effort$Total_days[i]       <- NA_real_  
        survey_effort$Total_minutes[i]    <- NA_real_  
        survey_effort$Missing_days[i]     <- NA_real_  
        survey_effort$Missing_minutes[i]  <- NA_real_  
        survey_effort$Net_days[i]         <- NA_real_  
        survey_effort$Net_minutes[i]      <- NA_real_  
        survey_effort$Perc_missing[i]     <- NA_real_  
      } else {  
        
        # Total duration  
        total_secs    <- as.numeric(difftime(end_dt, start_dt, units = "secs"))  
        total_days    <- total_secs / (60 * 60 * 24)  
        total_minutes <- total_secs / 60  
        
        # Missing based on global_perc_missing  
        perc_missing         <- global_perc_missing  
        total_missing_secs   <- total_secs * (perc_missing / 100)  
        total_missing_days   <- total_missing_secs / (60 * 60 * 24)  
        total_missing_minute <- total_missing_secs / 60  
        
        # Net duration  
        net_secs    <- total_secs - total_missing_secs  
        net_days    <- net_secs / (60 * 60 * 24)  
        net_minutes <- net_secs / 60  
        
        # Store back into data frame  
        survey_effort$Total_days[i]       <- total_days  
        survey_effort$Total_minutes[i]    <- total_minutes  
        survey_effort$Missing_days[i]     <- total_missing_days  
        survey_effort$Missing_minutes[i]  <- total_missing_minute  
        survey_effort$Net_days[i]         <- net_days  
        survey_effort$Net_minutes[i]      <- net_minutes  
        survey_effort$Perc_missing[i]     <- perc_missing  
      }  
    }  
    
    cat("\nRecomputed survey_effort table:\n")  
    print(survey_effort)  
    
    # Offer to save updated CSV  
    if (read_yes_no("\nDo you want to SAVE the UPDATED table (after recomputation) to CSV?")) {  
      cat("Enter file name for updated CSV (leave empty to overwrite 'camera_trap_survey_effort.csv'):\n> ")  
      new_file <- trimws(readline())  
      if (new_file == "") {  
        out_file_updated <- "camera_trap_survey_effort.csv"  
      } else {  
        out_file_updated <- new_file  
      }  
      
      write.csv(survey_effort, file = out_file_updated, row.names = FALSE)  
      cat("Updated survey effort table saved to:", normalizePath(out_file_updated), "\n")  
    } else {  
      cat("Updated table NOT saved. You can still save it manually from 'survey_effort'.\n")  
    }  
    
  } else {  
    cat("Metrics not recomputed.\n")  
  }  
  
} else {  
  cat("\nNo survey_effort object with data found. Nothing to recompute.\n")  
}  


## ============================================================  
## Create clean summary data frame and save to CSV  
## ============================================================  
  
if (exists("survey_effort") && is.data.frame(survey_effort) && nrow(survey_effort) > 0) {  
  
  # Pick only the desired columns (if they exist)  
  wanted_cols <- c(  
    "Survey",        # optional but usually present  
    "Start",  
    "End",  
    "Total_days",  
    "Total_minutes",  
    "Missing_days",  
    "Missing_minutes",  
    "Net_days",  
    "Net_minutes",  
    "Perc_missing"  
  )  
  
  # Keep only columns that exist in survey_effort  
  used_cols <- intersect(wanted_cols, names(survey_effort))  
  
  summary_df <- survey_effort[, used_cols, drop = FALSE]  
  
  cat("\nSummary data frame to be saved:\n")  
  print(summary_df)  
  
  # Ask for file name  
  cat("\nEnter file name for the summary CSV (leave empty for 'camera_trap_survey_summary.csv'):\n> ")  
  out_file <- trimws(readline())  
  if (out_file == "") {  
    out_file <- "camera_trap_survey_summary.csv"  
  }  
  
  # Save to CSV  
  write.csv(summary_df, file = out_file, row.names = FALSE)  
  cat("Summary table saved to:", normalizePath(out_file), "\n")  
  
} else {  
  cat("\nNo valid 'survey_effort' object found (or it has zero rows). Nothing to export.\n")  
}  
```
