## Supporting Information 2 for "Species ecology, behavior, and substrate type shape eDNA detection of mammals at savanna waterholes": trophlev_test.R

```
# Script to calculate the effect of the trophic level onto eDNA detection rates

# Set wd
setwd(dirname(rstudioapi::getActiveDocumentContext()$path))

# Load required packages

library(dplyr)
library(FSA)
## Aggregating the individual observations first on the species level


data <- read.csv2("eDNA_CT_combined_sampling_events.csv")

# Calculating the means:
# 1) Species-level means
species_summary <- data %>%
  group_by(Species, TrophLev) %>%
  summarise(
    Species_Mean_Detection = mean(detection_rate, na.rm = TRUE),
    Species_N = sum(!is.na(detection_rate)),
    .groups = "drop"
  )

# 2) Trophic-level summary based on species means
summary_table <- species_summary %>%
  group_by(TrophLev) %>%
  summarise(
    Mean_Detection_Rate = mean(Species_Mean_Detection, na.rm = TRUE),
    SD_Detection_Rate   = sd(Species_Mean_Detection, na.rm = TRUE),
    N_Species           = n(),
    .groups = "drop"
  ) %>%
  rename(Trophic_Level = TrophLev)

print(summary_table)


species_data <- data %>%
  group_by(Species, TrophLev) %>%
  summarise(
    mean_detection = mean(detection_rate, na.rm = TRUE),
    n = sum(!is.na(detection_rate)),
    .groups = "drop"
  )

# Kruskal–Wallis on species-level means
kruskal.test(mean_detection ~ TrophLev, data = species_data)

#  Dunn's test for pairwise comparisons

dunnTest(mean_detection ~ TrophLev, data = species_data, method = "holm")
```
