## Supporting Information 2 for "Species ecology, behavior, and substrate type shape eDNA detection of mammals at savanna waterholes": 14_modelling_detection_Rate_traildata.R

```
# Script for modelling model influence of CT (and species trait) predictors onto eDNA dectection

# Set wd
setwd(dirname(rstudioapi::getActiveDocumentContext()$path))

## Load required package  

library(dplyr)  
library(tidyr)  
library(stringr) 
library(purrr) 
library(lubridate) 
library(lme4)
library(DHARMa)


# 1. Read and prepare data 
data_sampling_events <- read.csv("eDNA_CT_combined_sampling_events_trailsonly.csv")
data_sampling_events$ind_obs[is.na(data_sampling_events$ind_obs)] <- 0

data_sampling_events <- data_sampling_events %>%  
  mutate(  
    Species  = factor(Species),  
    location = factor(location),  
    period   = factor(period),  
    season   = factor(season, levels = c("dry", "wet")),  
    TrophLev = factor(TrophLev),
    
    AdultBodyMass_sc    = as.numeric(scale(AdultBodyMass)),  
    AdultHeadBodyLen_sc = as.numeric(scale(AdultHeadBodyLen)),  
    SocialGrpSize_sc    = as.numeric(scale(SocialGrpSize)),  
    
    ind_obs_sc      = as.numeric(scale(ind_obs)),  
    n_vis_sc        = as.numeric(scale(n_vis_events)), 
    time_same_sc    = as.numeric(scale(time_since_last_visit_sameperiod)),  
    time_overall_sc = as.numeric(scale(time_since_last_visit_overall)),
    behav_perz_sc   = as.numeric(scale(behav_perz)),  
    
  )


# 2. Full model
m_full <- glmer(
  cbind(num_detection, number.of.samples - num_detection) ~
    AdultBodyMass_sc +
    AdultHeadBodyLen_sc +
    SocialGrpSize_sc +
    SocialGrpSize_missing +
    TrophLev +
    ind_obs_sc +
    n_vis_sc +
    time_same_sc +
    time_overall_sc +
    no_prev_visit_sameperiod +
    no_prev_visit_overall +
    behav_perz_sc +
    behav_perz_missing +
    (1 | Species) +
    (1 | location) +
    (1 | period),
  data   = data_sampling_events,
  family = binomial(link = "logit"),
  control = glmerControl(optimizer = "bobyqa",
                         optCtrl   = list(maxfun = 1e5))
)


summary(m_full)


m_red <- glmer(
  cbind(num_detection, number.of.samples - num_detection) ~
    # species traits
    AdultHeadBodyLen_sc +
    SocialGrpSize_sc +
    TrophLev +
    
    # CT activity metrics (reduced set)
    n_vis_sc +
    time_same_sc +
    behav_perz_sc +
    
    # random intercepts
    (1 | Species) +
    (1 | location) +
    (1 | period),
  data   = data_sampling_events,
  family = binomial(link = "logit"),
  control = glmerControl(optimizer = "bobyqa",
                         optCtrl   = list(maxfun = 1e5))
)

summary(m_red)

sim_red <- simulateResiduals(m_red, n = 1000)
plot(sim_red)
testDispersion(sim_red)
testZeroInflation(sim_red)


anova(m_red, m_full, test = "Chisq")
AIC(m_full, m_red)
```
