## Supporting Information 2 for "Species ecology, behavior, and substrate type shape eDNA detection of mammals at savanna waterholes": README.md

The raw data of the camera trapping survey exported from Timelapse can be found in the directory 01_raw_data

The subsequent directories contain the R scripts, input files and generated output files of the statistical analyses.

The numbering of the directories follows the chonological order described in the manuscript.

The following files had to be compressed into zip files due to size restrictions of github and needs to be unzipped before usage:

04_species_names/04_CT_data_incl_latin.zip ->  04_species_names/04_CT_data_incl_latin.csv 
05 period assignment/CT_data_periods_added.zip -> 05 period assignment/CT_data_periods_added.csv
06 adding visitation events/CT_data_visitation_events_added.zip -> 06 adding visitation events/CT_data_visitation_events_added.csv
09 data prep for modelling/CT_data_visitation_events_added.zip -> 09 data prep for modelling/CT_data_visitation_events_added.csv 
